# High-throughput screening identifies small molecule inhibitors of thioesterase superfamily member 1: Implications for the management of non-alcoholic fatty liver disease

**DOI:** 10.1101/2022.10.15.512369

**Authors:** Christopher S. Krumm, Lavoisier Ramos-Espiritu, Renée S. Landzberg, Carolina Adura, Xu Liu, Mariana Acuna, Yang Xie, Xu Xu, Matthew C. Tillman, Yingxia Li, J. Fraser Glickman, Eric A. Ortlund, John D. Ginn, David E. Cohen

## Abstract

Thioesterase superfamily member 1 (Them1; synonyms Acyl-CoA thioesterase 11 (Acot11) and steroidogenic acute regulatory protein-related lipid transfer (START) domain 14 (StarD14) is a long chain acyl-CoA thioesterase comprising two N-terminal hot-dog fold enzymatic domains linked to a C-terminal lipid-sensing START domain, which allosterically modulates enzymatic activity. Them1 is highly expressed in thermogenic adipose tissue, where it functions to suppress energy expenditure by limiting rates of fatty acid oxidation. Its expression is also induced markedly in liver in response to high fat feedings, where it suppresses fatty acid oxidation and promotes hepatic glucose production. Mice lacking the gene (*Them1^-/-^*) are protected against diet-induced non-alcoholic fatty liver disease (NAFLD), suggesting Them1 as a therapeutic target. The current study was designed to develop small molecule inhibitors of Them1 and to establish their activities *in vitro* and in cell culture. High-throughput screening combined with counter screening assays were leveraged to identify two lead allosteric inhibitors that selectively inhibited Them1 by binding the START domain. In primary mouse brown adipocytes, these inhibitors promoted fatty acid oxidation, as evidence by increased rates of oxygen consumption. In primary mouse hepatocytes, they similarly promoted fatty acid oxidation, but also reduced glucose production. Optimized Them1 inhibitors could provide an attractive modality for the pharmacologic management of NAFLD and obesity-associated metabolic disorders.

## INTRODUCTION

Non-alcoholic fatty liver disease (NAFLD) is a common multigenic disorder that manifests in response to overnutrition (Younossi et al., 2016). Despite an alarming rise in the prevalence of NAFLD, current therapies are limited. Following uptake into cells, fatty acids are activated by esterification to coenzyme A (CoA) molecules by acyl-CoA synthetases. Fatty acyl-CoAs may then be oxidized or incorporated into complex lipids. Mammalian cells express acyl-CoA thioesterases (Acots), which hydrolyze fatty acyl-CoAs to form fatty acids and CoA. Emerging data implicate Acots in the pathogenesis of NAFLD (Tillander et al., 2017).

Acots are divided into two subfamilies consisting of type I isoforms (i.e. 1 – 6), which contain a conserved α/β hydrolase catalytic domain as well as high sequence homology, and type II isoforms (i.e. 7 – 15), which share common structural features including a characteristic ’HotDog domain’ motif, but exhibit low sequence similarity (Tillander et al., 2017). Thioesterase superfamily member 1 (Them1) [synonyms: Acot11 and steroidogenic acute regulatory protein-related lipid transfer (START) domain 14 (StarD14)] is a type II long-chain fatty acyl-CoA thioesterase comprising two thioesterase, as well as one regulatory lipid binding domain. It is highly expressed in thermogenic adipose tissue including brown adipose tissue (BAT) and beige adipose tissue (BeAT), and is robustly upregulated in response to cold ambient temperatures (Zhang et al., 2012 and unpublished findings). BAT and BeAT in rodents and humans are rich in mitochondria and promote energy expenditure by non-shivering thermogenesis. Mice with genetic ablation of Them1 (*Them1^-/-^*) exhibit increased thermogenesis and are resistant to diet-induced obesity and NAFLD (Zhang et al., 2012), indicating that Them1 functions biologically to conserve energy, even when doing so becomes pathologic. Mechanistically, Them1 limits hydrolysis of long-chain fatty acyl-CoAs, which are destined for mitochondrial fatty acid oxidation to enable thermogenesis (Okada et al., 2016). In response to high-fat feeding, Them1 expression is also induced in liver and white adipose tissue (WAT), and appears to contribute toward NAFLD pathogenesis (Desai et al., 2018; Zhang et al., 2012). Consistent with a maladaptive role in response to overnutrition, *Them1* maps to syntenic regions of human chromosome 1 and mouse chromosome 4 that are associated with adiposity and diet-induced obesity (Adams et al., 2001).

Them1 self-assembles as a homotrimer and is one of two Acots that comprises tandem N-terminal thioesterase domains fused to a C-terminal lipid-sensing START domain (Han and Cohen, 2012; Tillander et al., 2017; Tillman et al., 2020). START domains often reside within multi-domain proteins, suggesting that lipid binding may regulate protein activity (Alpy and Tomasetto, 2005). The Them1 START domain regulates the enzymatic domains in response to binding long-chain fatty acids, as well as lysophosphatidylcholines, which activate and suppress acyl-CoA thioesterase activity, respectively (Han and Cohen, 2012; Tillman et al., 2020).

Based on the premise that Them1 might represent an attractive target in the management of NAFLD, we undertook an effort to develop small molecule Them1 inhibitors. Leveraging a high-throughput screen (HTS) followed by counter screening targeting related Acot isoforms, we identified selective inhibitors of Them1 activity with binding sites that include the enzymatic and START domains. Preliminary structure activity relationship (SAR) analyses of selected small molecules revealed molecular features that influenced Them1 enzymatic activity and binding interactions. The most promising inhibitors reversed Them1-mediated increases in rates of glucose production in cultured hepatocytes, and reversed Them1’s suppressive effects on rates of fatty acid oxidation in both cultured brown adipocytes and hepatocytes.

## RESULTS

### Assay for high-throughput screening against Them1 activity

A fluorometric Acot activity assay was developed in 384-well microplate format: Free CoA, liberated by Them1 activity, is conjugated to a non-fluorescent reagent to generate a fluorescent product (Supplemental figure 1A). We utilized full-length human Them1 for the HTS to capture either competitive and allosteric small molecule inhibitors with binding sites on the thioesterase or START domains. Expression followed by affinity chromatography yielded recombinant His-tagged human Them1 of 91% purity (Supplemental figure 1B), with protein identity verified by native mass spectrometry. Because of its relatively high monomeric solubility, myristoyl (C14)-CoA was selected as the substrate for this assay to avoid micelle formation at concentrations desired for the screen (Wei et al., 2009). Preliminary experiments established optimized conditions of 125 nM recombinant Them1 and 25 µM C14-CoA for 60 min at 22 °C, which were assessed based on product formation as a function of recombinant Them1 concentration (Supplemental figure 1C), as well as reaction time (Supplemental figure 1D). Steady state kinetic constants (K_m_: 6.39 μM, V_max_: 44.4 nmol/min/mg) (Supplemental figure 1E) were in good agreement with our previously published values (Han and Cohen, 2012). To determine the suitability of Them1 as a druggable target and the performance and reproducibility of the assay, we employed a test library of 1,056 pharmacologically active compounds (LOPAC) revealing several candidate inhibitors (Supplemental figures 1F - G).

### Small molecule inhibitors with selectivity toward Them1 activity

Given the promising results from the LOPAC screen, we conducted a HTS of 360,705 compounds from a highly diverse small molecule library in 384-well microplates. Z’ factors ≥ 0.7 (Figure 1A) exceeded the 0.5 threshold of an ‘excellent’ assay (Zhang et al., 1999) and 330 compounds (0.09% hit rate) fulfilled the screening criteria of a normalized percentage of inhibition (NPI) ≥ 30 (Figures 2B - C). From this hit set, 174 / 300 (53%) of compounds demonstrated reproducibility and concentration-dependent inhibition with IC_50_ values falling in the low micromolar range (Figure 1D).

**Figure 1.**
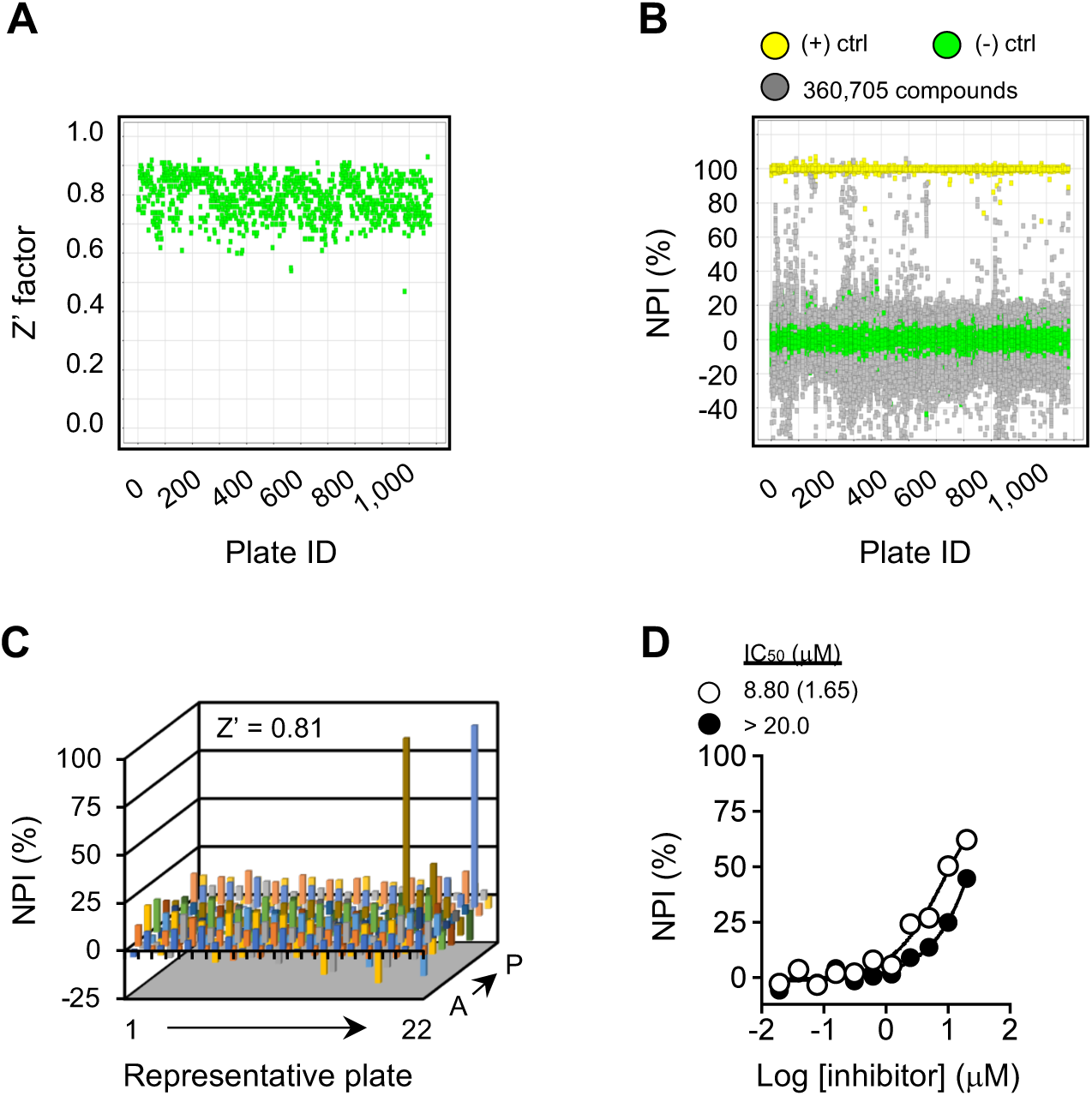
Small molecule inhibitors targeting Them1 activity identified by high-throughput screening. Reactions were performed in 384-well microplates with recombinant His-tagged human Them1 (Them1; 125 nM), myristoyl-CoA (C14-CoA; 25 μM) and compounds (12.5 μM), and incubated for 60 min at 22°C. **(A)** Z’ factors were calculated from reactions containing no enzyme [(+) control] or C14-CoA [(-) control; 25 μM]. Plate ID indicates the unique number assigned for each plate in the compound library. **(B)** Scatter plot for the NPI of 360,705 library compounds (yellow dots [(+) control)]; green dots [(-) control] and gray dots [compounds (12.5 μM)]). **(C)** Representative 384-well microplate from the high-throughput screen. Compounds were considered as potential inhibitors based on a normalized percent inhibition (NPI) ≥ 30%. Negative values are attributable to the absorbance of a minority of compounds. The Z’ factor is shown. **(D)** Representative compounds that exhibited inhibition of Them1 activity (open circles) or exceeded the maximum IC_50_ screening criteria of 20 μM (closed circles). Data represent mean (s.e.m.) of triplicate determinations. Where not visible, standard error bars are contained within the symbol sizes.

**Figure 2.**
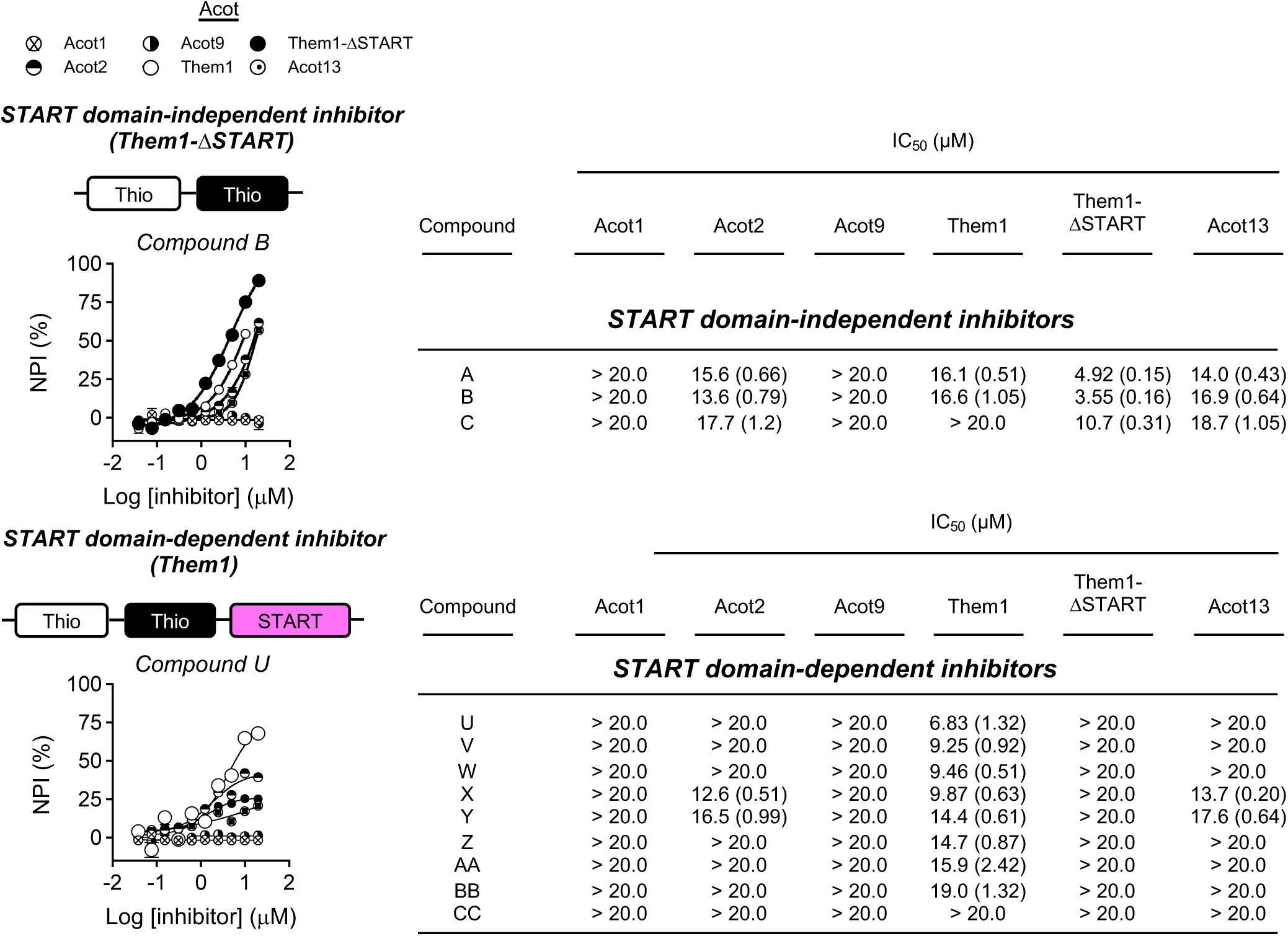
Small molecule inhibitors selectively targeting Them1 activity. Reactions were performed in 384-well microplates with recombinant Acot isoforms (125 nM), myristoyl-CoA (C14-CoA; 25 μM) and compounds (19.5 nM to 20 μM), and incubated for 60 min at 22 °C. Recombinant Acot isoforms included truncated Them1 containing only the 2 thioesterase (Thio) domains but lacking the START domain (Them1-ΔSTART), and type I [Acot1, Acot2] and type II [Acot9, Them1, Acot13] enzymes. *Left panel*: Representative compounds selectively inhibiting Them1 activity either through the thioesterase domains (Compound **B**) or the START domain (Compound **U**). Where not visible, standard error bars are contained within the symbol sizes. *Right panel*: Table containing compounds exhibiting START domain-independent or -dependent Them1 inhibition. Data represent mean (s.e.m.) of triplicate determinations.

A counter screen of the 174 compounds was performed to determine specificity for Them1 relative to other Acot isoforms [type I: Acot1 and Acot2, type II: Acot9 and Acot13] (Tillander et al., 2017). Expression and purification of these recombinant isoforms in *E. coli* yielded highly pure and active enzymes (Supplemental figures 2A - G). Comparable values of steady state kinetic constants among recombinant Acot isoforms excluded the possibility that differences in selectivity and potency of compounds instead represented an artifact due to differences in catalytic efficiency during the time course of the Acot activity assay (Supplemental figure 2H). Compounds targeting recombinant Them1 with NPI > 40 and recombinant Acot isoforms with NPI ≤ 40 were considered to be selective for Them1. Whether inhibition was dependent upon the thioesterase domains or the START domain was assessed by comparing Them1 to a truncated Them1 comprising only the thioesterase domains (Them1-βSTART). Of the 12 Them1-selective compounds, those with comparable NPI for both recombinant Them1 and Them1-βSTART enzymes were considered START domain-independent. By contrast, compounds with significantly higher NPI for recombinant Them1 versus Them1-βSTART enzymes were considered START domain-dependent.

This counter screen identified 12 / 174 compounds that exhibited START domain-independent (3 compounds) or -dependent (9 compounds) inhibition (Figure 2). Five START domain-dependent (compounds **U**, **V**, **W**, **Y** and **AA**) inhibitors were available for purchase from commercial vendors and yielded IC_50_ values comparable to those obtained during the validation and counter screens (data not shown).

### Structural-based optimization of lead small molecule inhibitors

Of the 9 START domain-dependent compounds (Figure 2), **U** and **W** were selected for structure-activity relationship (SAR) expansion based on their drug-like properties and tractability for medicinal chemistry. As a basis for SAR, these compounds were designated as **U1** and **W1**, respectively. Because **W1** contained acetate as a functional group within its structure, we first excluded acetylation as an irreversible mechanism of Them1 inhibition. We used dialysis to assess reversibility of inhibition (Supplemental figure 3A). **U1** was used as a control because it does not contain a functional acetate group (Supplemental figure 3B). Consistent with reversible inhibition by both compounds, dialysis restored Them1 activity. By contrast, Them1 activity remained suppressed in reactions containing the covalent modifier Ebselen as a control (Dahlin and Walters, 2016). Anticipating use in cultured mouse cells and *in vivo*, we further verified that compounds **U1** and **W1** inhibited the activity of purified recombinant mouse Them1 with comparable potencies (Supplemental figure 3C) to their inhibition of human Them1 activity (Supplemental figure 3D).

Commercially available structural analogs of compounds **U1** and **W1** were identified and used to reveal molecular features important for inhibition of Them1 activity (IC_50_) and binding (K_d_) (Figure 3). Compound **U1** is characterized by a central hydrocarbon decorated with thiophene and piperazine heterocycle rings, and an indole moiety. Structural analogs of compound **U1** for which the piperazine heterocycle was replaced improved (**U3**) or retained comparable (**U2**, **U4** and **U5**) IC_50_ and K_d_ values to the parent **U1** compound. Compound **W1** is characterized by a benzoic acid core with acetate and phenylpropanamide functional groups. When compared to the parent **W1** compound, shortening of the alkyl chain (**W2**), truncation of the isopropyl ester (**W3**) or alkyl substitution (**W4**) yielded similar IC_50_ and K_d_ values. However, removal of the acetate (**W5**) to the free phenol resulted in complete loss in both activity and binding affinity.

**Figure 3.**
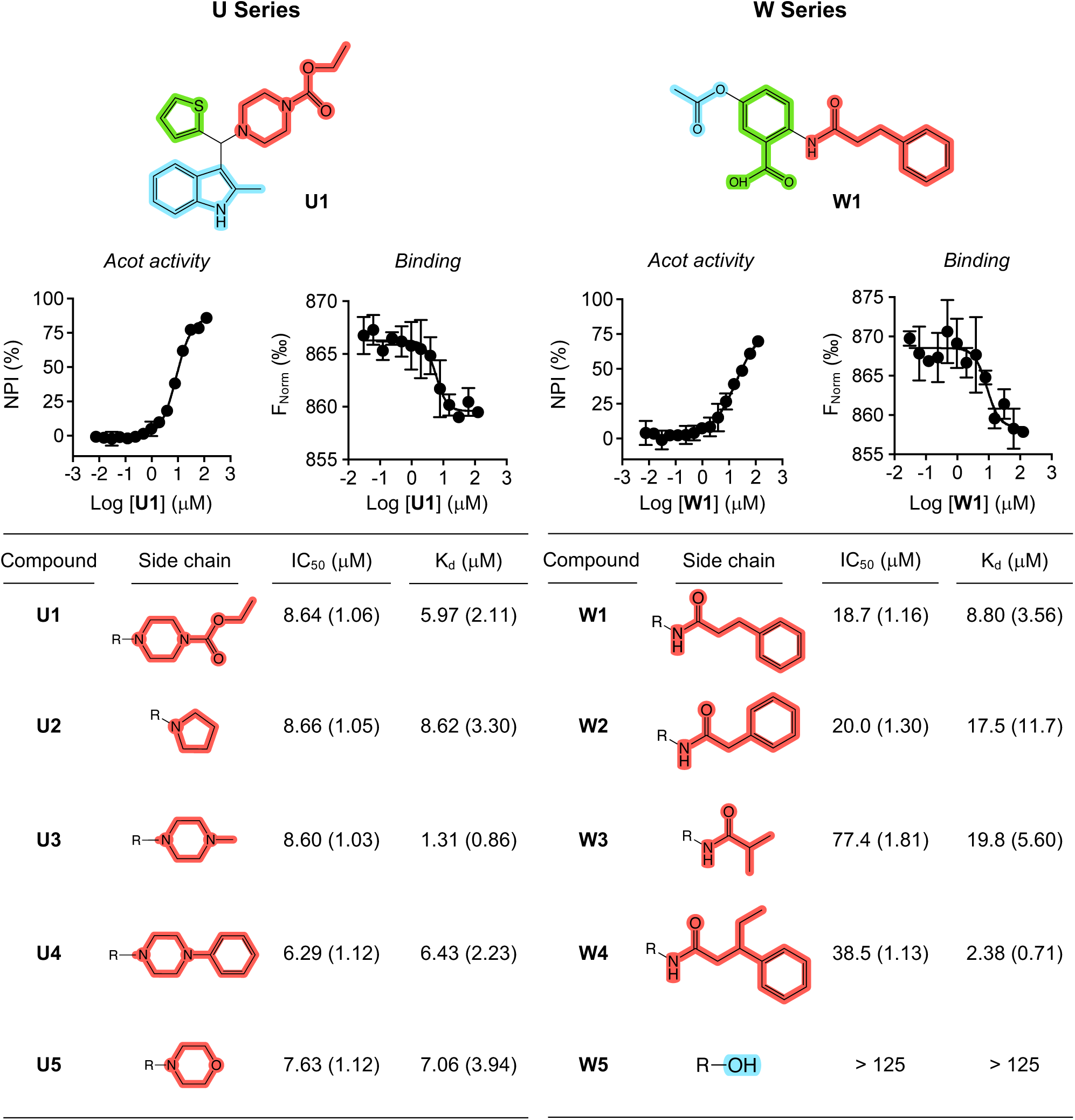
Structure activity relationship (SAR) expansion of commercially available small molecule inhibitors with START-domain dependent Them1 activity. Structural features of **U** and **W** series small molecules that selectively inhibit Them1 activity through its START domain. *Acot activity* was determined in 384-well microplates with recombinant His-tagged human Them1 (Them1; 125 nM), myristoyl-CoA (C14-CoA; 25 μM) for 60 min at 22 °C. IC_50_ values are listed. *Binding interactions* were quantified as K_d_ values by microscale thermophoresis using recombinant Them1 labeled with Monolith RED-Tris-NTA (100 nM). Data represent mean (s.e.m.) of triplicate determinations. Where not visible, standard error bars are contained within the symbol sizes.

### Them1 small molecule inhibitors reverse Them1-mediated suppression of oxygen consumption rates in primary brown adipocytes

Further development of compounds **U1** and **W1** as potential early lead compounds required a cell culture system with robust expression of active Them1. In this connection, we previously showed that endogenous Them1 expression was lost in cultured brown adipocytes, necessitating transduction with recombinant adenovirus in order to achieve expression and to demonstrate Them1-mediated suppression of fatty acid oxidation (Okada et al., 2016; Zhang et al., 2012). By contrast, cultured hepatocytes continued to express Them1 (Supplemental Figure 4A), although the level of expression diminished with time. To address these limitations, we generated a transgenic Them1 mouse from which we created mice that overexpress FLAG-Them1 in adipose tissue (*A-Them1Tg)* or in liver (*L-Them1Tg*). Mice harbored single copies of the transgene Them1 in the absence (wild type; Supplemental figure 4B) or the presence of a single copy of the transgene adiponectin-Cre (*A-Them1Tg;* Supplemental Figure 4C) or human thyroxine binding globulin (H1)-Cre (*L-Them1Tg*; Supplemental figure 4D). Adipose tissue-specific Them1 transgene overexpression was confirmed in *A*-*Them1Tg* mice according to increased *Them1* mRNA and protein abundance (Supplemental figure 4E - F) in WAT and BAT, but not other tissues including brain, heart, lung, skeletal muscle and kidney. FLAG protein abundance was detected exclusively in WAT and BAT of *A-Them1Tg* mice. In cultured brown adipocytes, Them1 protein was readily detected in brown adipocytes from *A-Them1Tg* mice (Supplemental figure 4G). Them1 protein was also robustly expressed in hepatocytes cultured from *L-Them1Tg* mice (Supplemental figure 4H).

A cell-based assay using primary brown adipocytes (Figure 4A) was optimized to measure OCR as a surrogate marker for fatty acid oxidation (Okada et al., 2016). Indicative of Them1 activity, OCR values following NE stimulation were blunted by Them1 expression in *A-Them1Tg* brown adipocytes. Treatment with compound **U1** in *A-Them1Tg* brown adipocytes reversed the suppressive effects of Them1 on values of OCR (Figure 4B). By contrast, treatment of cells with compounds **U2 – U5,** which exhibit similar IC_50_ and K_d_ values as the parent compound **U1** (Figure 3), demonstrated similar or reduced capacities to reverse the suppressive effects of Them1 on OCR (Figure 4C; Supplemental figure 5). Compound **W1** had no effect on reversing the suppressive effects of Them1 on OCR values in *A-Them1Tg* brown adipocytes (Figure 4D). Chemical modification with structurally modified analogs (**W2**; Figure 4E) were insufficient to completely reverse the suppressive effects of Them1 on OCR values in *A-Them1Tg* brown adipocytes. However, it is important to note that in these preliminary studies that compound **W1** and **W2** were tested at lower concentrations relative to their IC_50_ values *in vitro*.

**Figure 4.**
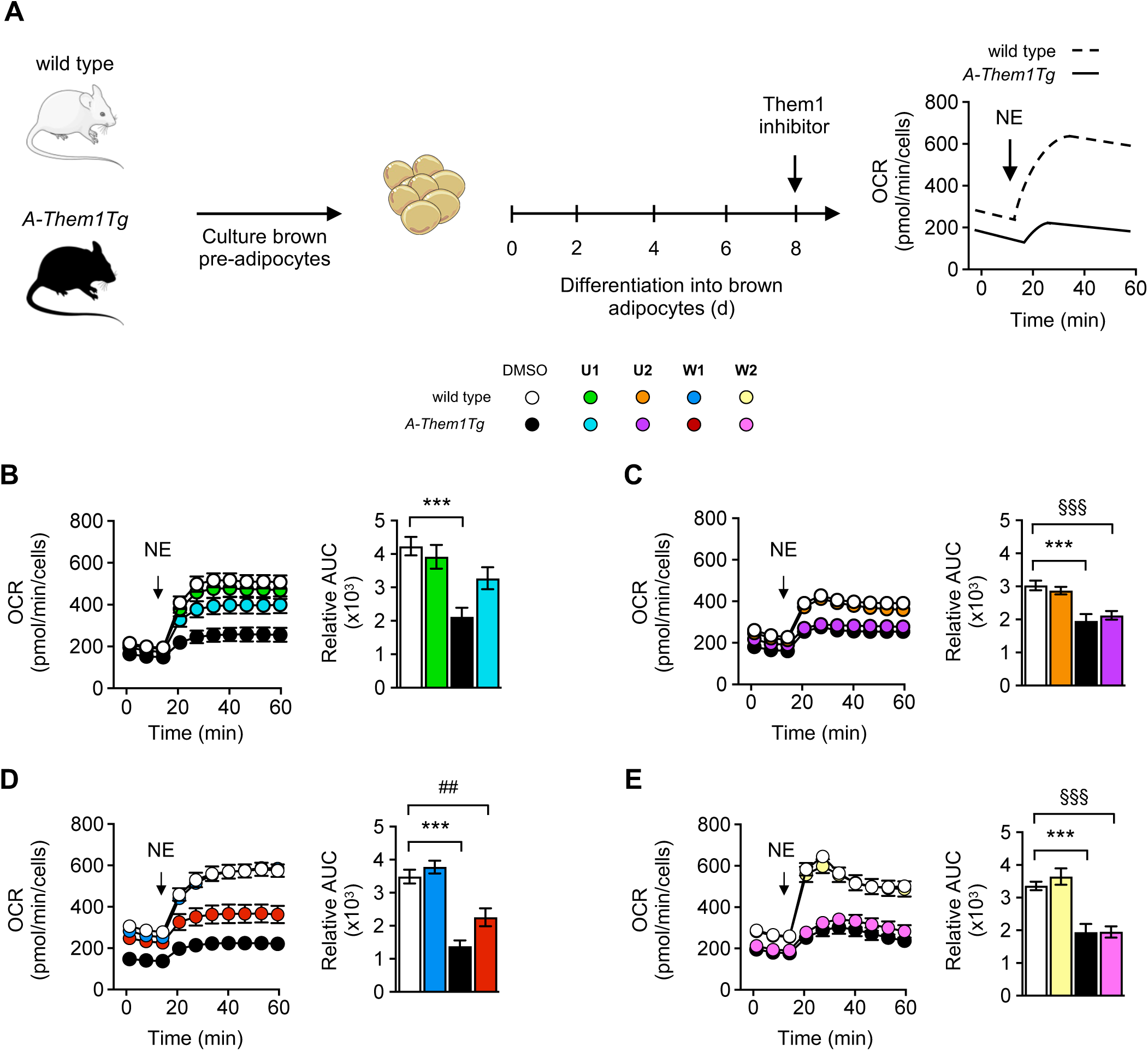
Them1 inhibitors increase fatty acid oxidation rates in cultured primary brown adipocytes. **(A)** Experimental design to test Them1 small molecule inhibitors in primary brown adipocytes cultured from wild type and transgenic adipose tissue-specific Them1 overexpression (*A-Them1Tg)* mice. **(B – E)** Oxygen consumption rate (OCR) values in primary brown adipocytes cultured from wild type and *A-Them1Tg* mice following stimulation with norepinephrine (NE; 1 μM), and treatment for 30 min at 10 μM with compounds **(B) U1**, **(C) U2**, **(D) W1** and **(E) W2**. The bar graphs in each panel represent relative values of area under the curve (AUC) following NE stimulation. Data in the graphs represent the mean ± s.e.m. of 2 to 3 independent experiments. Where not visible, standard error bars are contained within the symbol sizes. *P < 0.05; wild type (DMSO) vs. *A-Them1Tg* (DMSO); ***P < 0.001; wild type (DMSO) vs. *A-Them1Tg* (DMSO); _##_P < 0.01; wild type (DMSO) vs. *A-Them1Tg* (**W1**); ^§§§^P < 0.001; wild type (DMSO) vs. *A-Them1Tg* (**U2** or **W2**).

In cultured hepatocytes, cell-based assays were designed to measure OCR as a surrogate marker for fatty acid oxidation of exogenous fatty acids, and glucose production from gluconeogenic substrates (Figure 5A) (Desai et al., 2018; Kawano et al., 2014). OCR values following palmitate stimulation were reduced in *L-Them1Tg* hepatocytes (Figure 5B). Treatment with compound **U1** reversed this suppressive effect of Them1. Consistent with Them1-specific inhibition, compound **U1** did not increase OCR values in *Them1^-/-^*hepatocytes, which exhibited higher OCR values than wild type or *L-Them1Tg* hepatocytes (Figure 5B). Hepatic glucose production was increased by transgenic Them1 overexpression, and this was reduced by treatment with compounds **U1** and **W1**. Them1 inhibitors had no effect in *Them1^-/-^* hepatocytes, which exhibited lower glucose production rates compared with hepatocytes from wild type or *L-Them1Tg* mice (Figure 5C).

**Figure 5.**
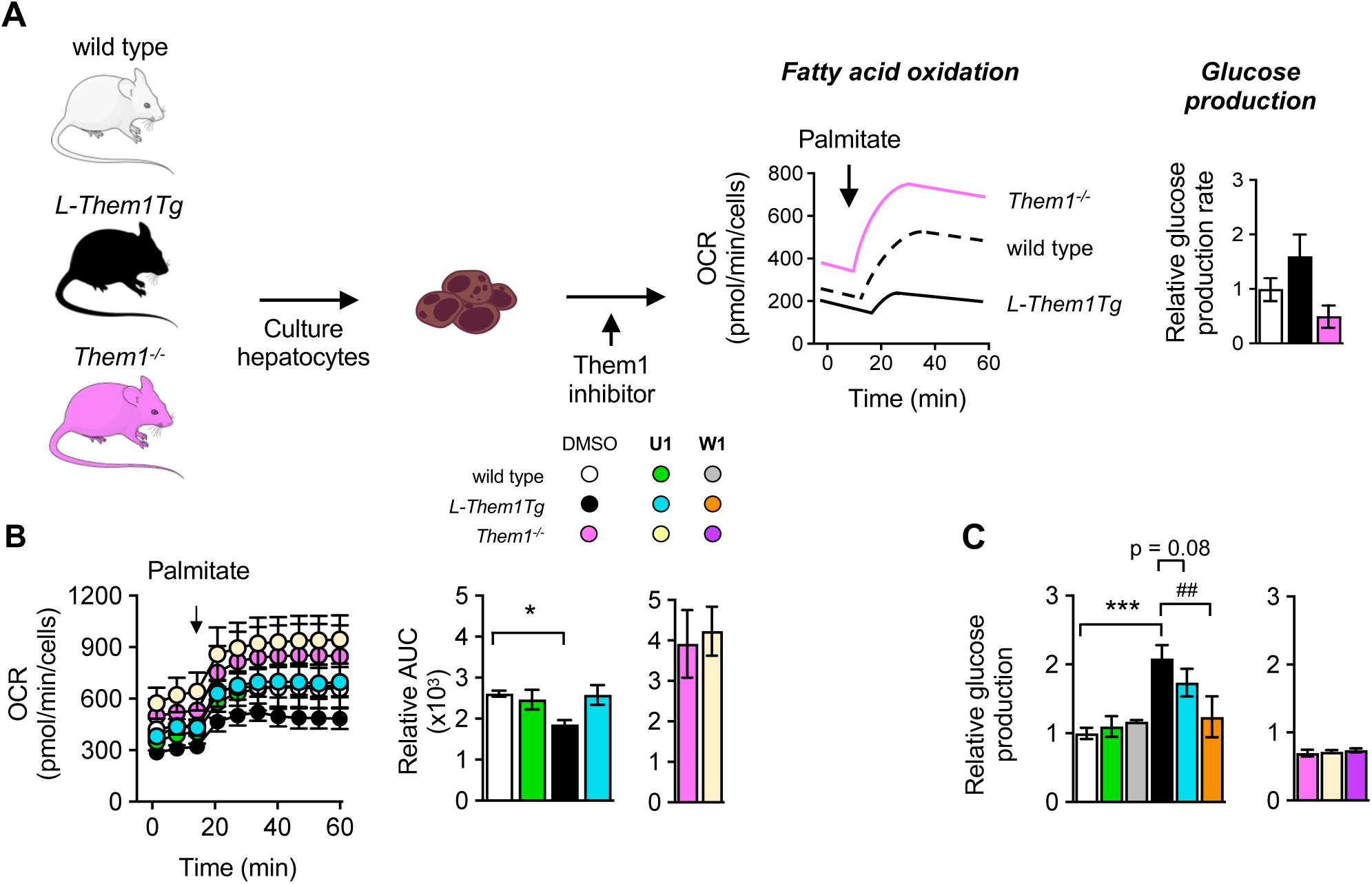
Them1 inhibitors increase fatty acid oxidation rates and suppress glucose production in cultured primary hepatocytes. **(A)** Experimental design to test Them1 small molecule inhibitors in primary hepatocytes cultured from wild type, transgenic liver-specific Them1 overexpression (*L-Them1Tg*) and Them1 knockout (*Them1*^-/-^) mice. **(B)** Oxygen consumption rate (OCR) values in primary hepatocytes following stimulation with palmitic acid conjugated with fatty acid-free BSA (300 μM) and treatment with compound **U1** (33 μM). The bar graphs in each panel represent relative values of area under the curve (AUC) following stimulation with palmitate. Data in the graphs represent the mean ± s.e.m. of 2 independent experiments. Where not visible, standard error bars are contained within the symbol sizes. *P < 0.05; wild type (DMSO) vs. *L-Them1Tg* (DMSO). **(C)** Relative glucose production compared to wild type controls in serum-starved primary hepatocytes were determined by appearance of glucose in media following addition of compound **U1** (33 µM) or **W1** (20 µM), and pyruvate (2 mM) and lactate (20 mM) as gluconeogenic substrates. Data in the graphs represent the mean ± s.e.m. of 2 independent experiments. ***P < 0.001; wild type (DMSO) vs. *L-Them1Tg* (DMSO); P = 0.08; *L-Them1Tg* (DMSO) vs. *L-Them1Tg* (**U1**); ***, P < 0.001; ^##^, *L-Them1Tg* (DMSO) vs. *L-Them1Tg* (**W1**).

### Chemical synthesis of novel U1 structural derivatives

Based on its promising chemical characteristics and its effectiveness *in vitro* and in cell culture, we focused on further development of compound **U1**. In order to exclude the possibility of interference in the *in vitro* assay by reactive impurities and non-specific activity (Dahlin and Walters, 2016), compound **U1** was re-synthesized, and the structure and purity was confirmed by high resolution mass spectrometry, ^1^H-NMR and ^13^C-NMR (Supplemental methods). The re-synthesized molecule demonstrated similar potency in inhibiting Them1 activity (IC_50_: 32.9 μM). The inhibition was confirmed to be reversible by overnight dialysis, resulting in the recovery of enzymatic activity (Supplemental figure 6).

To build upon the SAR from commercially available structural analogs, structural derivatives of the parent **U1** compound were explored (Supplemental methods) (Kalgutkar and Dalvie, 2015). The SAR of the parent **U1** compound focused on regions of the molecule that included the piperazine (red) or thiophene (green) rings and indole moiety (blue) (Figure 5). Using the IC_50_ of the resynthesized parent **U1** compound as a benchmark, SAR analyses building upon the piperazine heterocycle either improved [i.e. increasing the steric bulk of **U10** with an isopropyl group (**U11**) or replacement with an N-benzyl derivative (**U12**)], remained comparable [i.e. replacing the piperazine ring with piperidine (**U6**) with a matched pair isonipecotic acid ethyl ester (**U7**), as well as removal (**U9**) or replacement of the carbamate with an acetamide (**U10**) or benzaldehyde (**U13**)], or resulted in complete loss of activity [i.e. extension of the carbamate with a benzoyl group (**U8**), suggesting that this group was sterically too large to accommodate the binding pocket]. SAR expansion off the indole moiety either improved IC_50_ potencies by extending the 2-position alkyl substituent (**U14**) or remained comparable to the parent **U1** compound by substitution off the indole nitrogen (**U15**). Structural derivatives designed around the thiophene ring remained similar [i.e. addition of a methyl group (**U16**)] or negated [i.e. substitution with a phenyl ring (**U17**)] inhibition. Reversibility of inhibition was confirmed by dialysis for all compounds (Supplemental figure 7). Each of these compounds exhibited a critical aggregation concentration > 125 μM, negating the possibility Them1 inhibition might have attributable to compound aggregation (Supplemental figure 8).

Compound **U1** bound more tightly to the START domain (K_d_: 1.03 μM) compared to full-length Them1 (K_d_: 8.00 μM) and lacked binding to Them1-ΔSTART (Figure 6). SAR-dependent differences in IC_50_ values relative to the parent **U1** compound were associated with tighter (e.g. **U11, U12** and **U14**), comparable (e.g. **U7**, **U13** and **U16**) or negligible (e.g. **U8**) binding affinities to full-length Them1 and the START domain, but consistently did not bind Them1-ΔSTART. In further support as a START domain-dependent inhibitor selective for Them1, the parent **U1** compound and its structural derivatives lacked the capacity to bind Acot12, a related Acot isoform also containing a C-terminal START domain.

**Figure 6.**
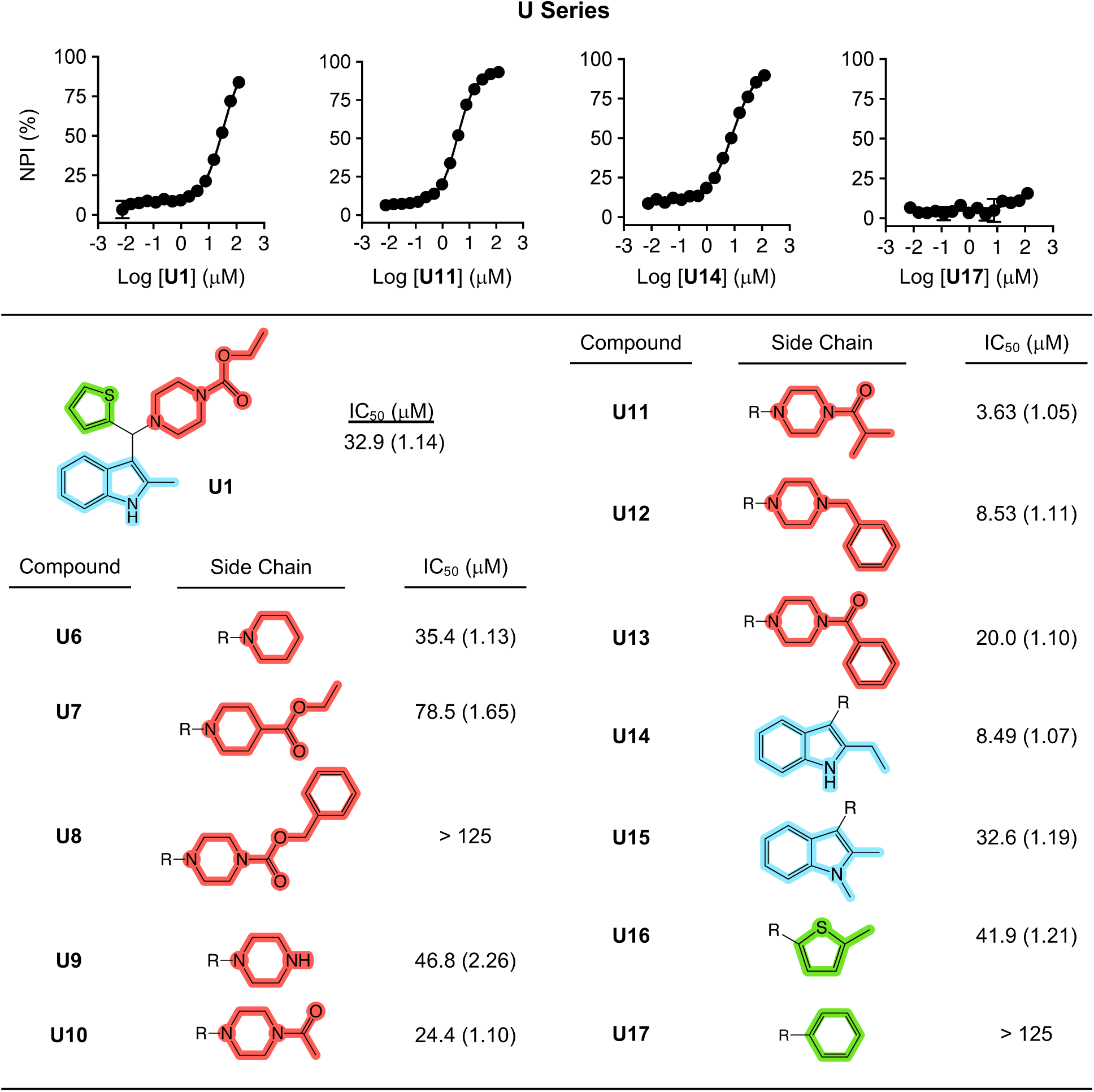
START domain-dependent inhibition of Them1 activity by freshly synthesized compound U1 and novel structural derivatives. Chemical features of freshly synthesized compound U1 and novel structural derivatives (U6 – U17). Reactions were performed in 384-well microplates with recombinant His-tagged human Them1 (Them1; 125 nM), myristoyl-CoA (C14-CoA; 25 μM) and compounds incubated for 60 min at 22 °C. Data represent the mean (s.e.m.) of triplicate determinations. Where not visible, error bars are contained within the symbol sizes.

The potency of structural derivatives of the parent **U1** compound were tested at their *in vitro* IC_50_ concentrations using the brown adipocyte-based OCR assay. *A-Them1Tg* brown adipocytes treated with structural derivatives (i.e. **U6**, **U9**, **U11** and **U12)** reversed the suppressive effects of Them1 on values of OCR compared to wild type brown adipocytes treated with DMSO (Figure 8A – C). By contrast, OCR values in *A-Them1Tg* brown adipocytes remained suppressed upon treatment with compounds **U8**, **U14** and **U15** (Figures 8D – E). The possibility of compound cellular toxicity was discounted by demonstrating that compounds **U1** and **W1** and their structural derivatives exhibited half-maximal cellular lethal concentrations (LC_50_) > 125 μM (Supplemental Figure 9).

## DISCUSSION

Our interest in Them1 as a therapeutic target evolved based on evidence toward its contribution to the pathogenesis of NAFLD in a common high-fat diet-fed mouse model (Desai et al., 2018; Zhang et al., 2012). Support for a pathogenic role of Them1 in human NAFLD can be gleaned from re-analyses of unbiased gene expression studies of human tissues (Desai et al., 2018; Okada et al., 2016; Zhang et al., 2012; unpublished findings). Upregulation of Them1 in human BAT in response to cold exposure (Figure 7A) is consistent with observations in mice (Adams et al., 2001; Zhang et al., 2012). Upregulation of Them1 in livers of patients with NAFLD relative to lean subjects (Figure 7B) is in keeping with its maladaptive role in promoting steatosis and hepatic glucose production (Desai et al., 2018), and upregulation in livers of non-alcoholic steatohepatitis relative to NAFLD patients (Figure 7C) is consistent with its contribution to hepatic inflammation (Kazankov et al., 2019; Zhang et al., 2012). Increased expression of Them1 in both WAT of obese relative to lean human subjects (Figure 7D) and visceral WAT relative to subcutaneous WAT (Figure 7E) suggest its contribution to WAT inflammation (Kazankov et al., 2019; Zhang et al., 2012). When taken together, these observations prompted us to explore whether pharmacological interventions using small molecule inhibitors to inhibit Them1 enzymatic activity could recapitulate the NAFLD-resistant phenotypes of *Them1^-/-^*mice under conditions of overnutrition and to serve as proof-of-concept for a novel therapeutic modality in the management of NAFLD.

**Figure 7.**
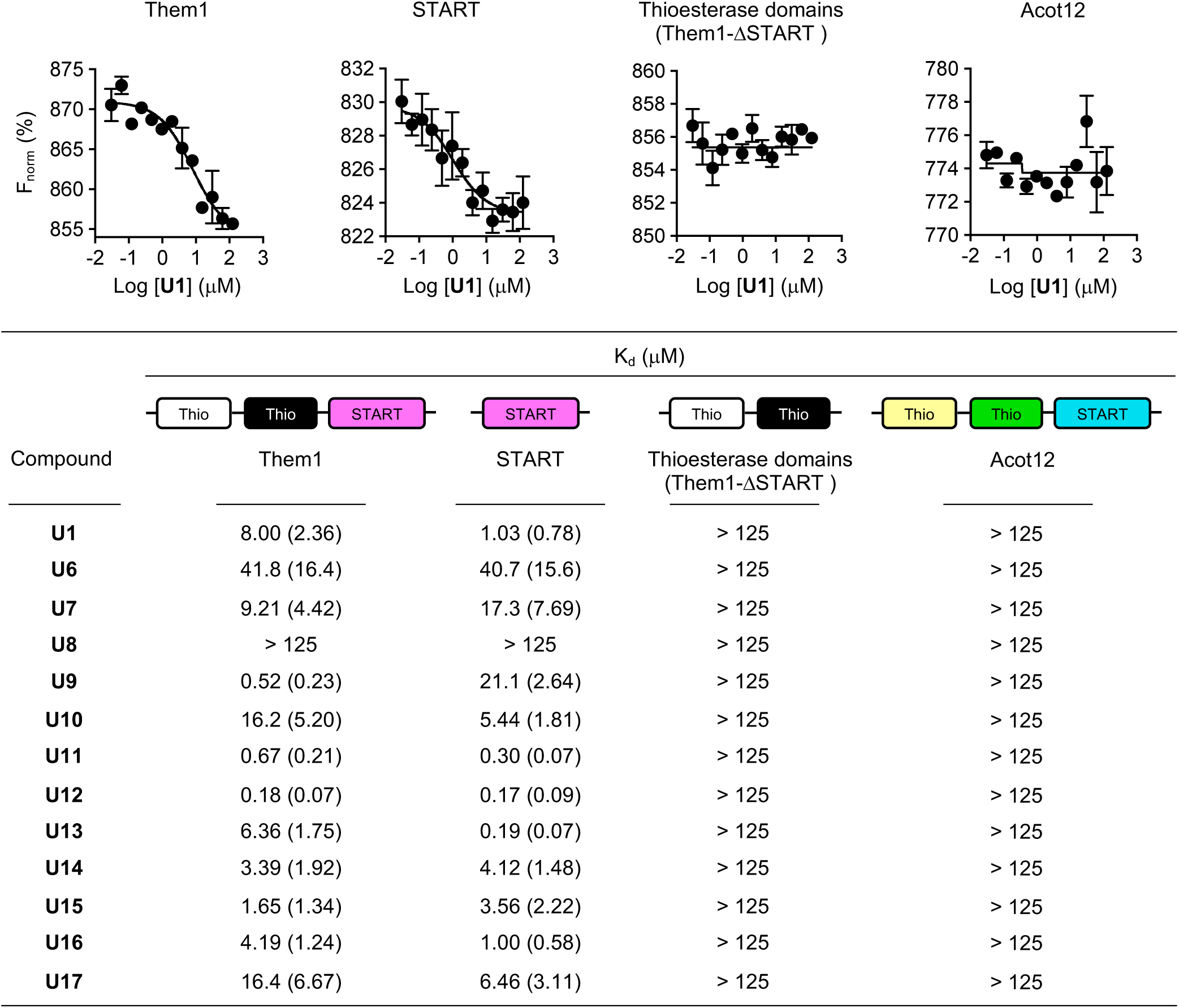
Compound U1 and structural derivatives selectively bind the Them1 START domain. Binding interactions were quantified as K_d_ values by microscale thermophoresis using recombinant His-tagged proteins labeled with Monolith RED-Tris-NTA (100 nM) (i.e. full-length Them1, a truncated Them1 either containing only the START domain but lacking the 2 thioesterase domains [START], only the 2 thioesterase domains but lacking the START domain [Them1-ΔSTART] or full-length human Acot12 [Acot12]) and compounds. Data represent the mean (s.e.m.) of triplicate determinations. Where not visible, error bars are contained within the symbol sizes.

**Figure 8.**
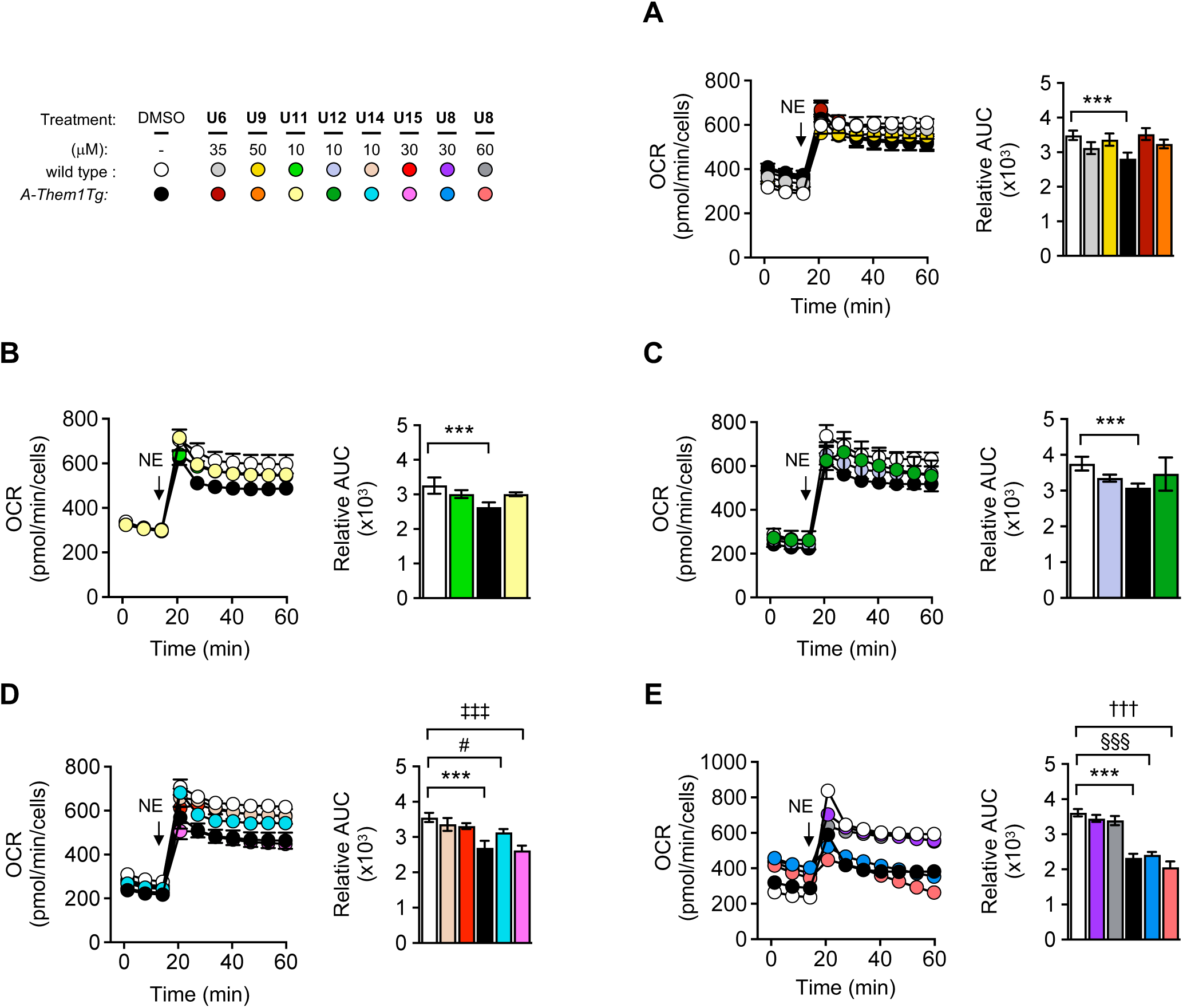
Small molecule inhibitors increase oxygen consumption rates (OCR) in cultured primary brown adipocytes. **(A - E)** OCR values in primary brown adipocytes cultured from wild type and transgenic adipose tissue-specific Them1 overexpression (*A-Them1Tg*) mice following stimulation with norepinephrine (NE; 1 μM), and treatment for 30 min with the following compounds: **(A) U6** and **U9**, **(B) U11 (C) U12**, **(D) U14** and **U15** and **(E) U8**. The bar graphs in each panel represent relative values of area under the curve (AUC) following NE stimulation. Data in the graphs represent the mean ± s.e.m. of 2 independent experiments. Where not visible, standard error bars are contained within the symbol sizes. ***, P < 0.001; wild type (DMSO) vs. *A-Them1Tg* (DMSO); #, P < 0.01; wild type (DMSO) vs. *A-Them1Tg* (**U14**); ‡‡‡, P < 0.001; wild type (DMSO) vs. *A-Them1Tg* (**U15**); §§§, P < 0.001; wild type (DMSO) vs. *A-Them1Tg* (**U8**; 30 μM); †††, P < 0.001; wild type (DMSO) vs. *A-Them1Tg* (**U8**; 60 μM).

**Figure 9.**
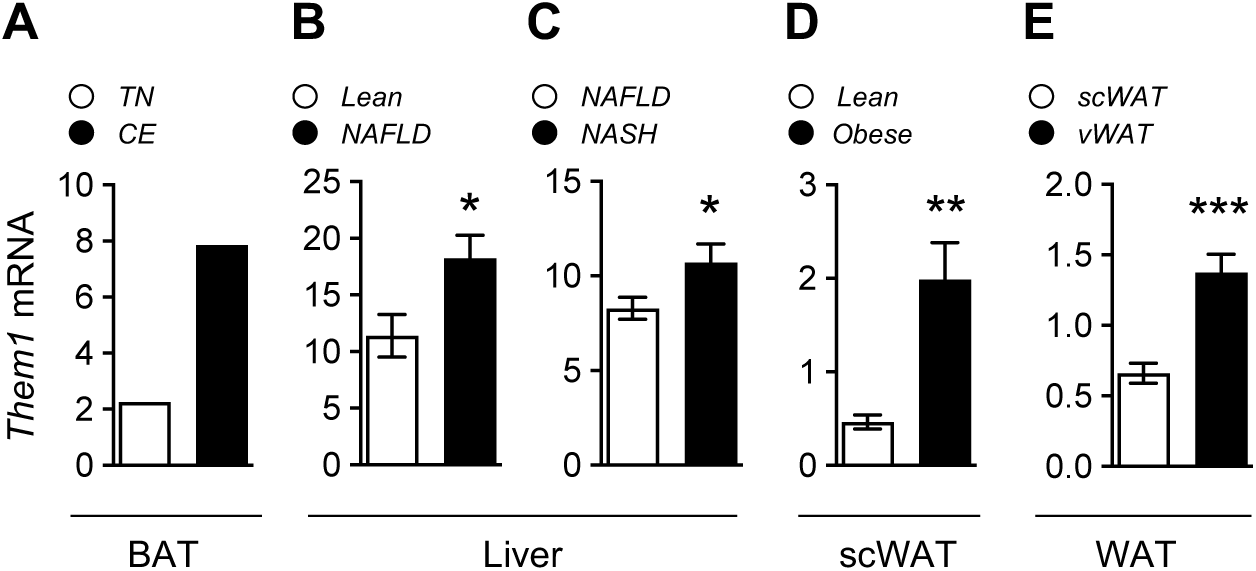
Evidence supporting a pathogenic role of Them1 in human NAFLD. *Them1* mRNA expression levels measured in brown adipose tissue (BAT), liver and white adipose tissue from human subjects were gleaned from the NCBI GEO or ArrayExpress databases: (**A**) BAT samples from a human subject who volunteered to underdo biopsy at thermoneutrality (26 ± 1.2 °C) and under cold exposure (18.2 ± 2.1 °C) (Accession #: E-MTAB-4031). (**B**) Liver samples from lean subjects and patients with non-alcoholic fatty liver disease (NAFLD). Lean, n = 10; NAFLD, n = 10. (Accession #: GSE135776). (**C**) Liver samples from patients NAFLD and non-alcoholic steatohepatitis (NASH). NAFLD, n = 51; NASH, n = 47. (Accession #: GSE167523). (**D**) Subcutaneous white adipose tissue (scWAT) from lean and obese human subjects. Lean, n = 10; Obese, n = 10. (Accession #: GSE162653). (**E**) scWAT and visceral WAT (vWAT) in obese human subjects. scWAT, n = 10; vWAT, n = 83. (Accession #: GSE135251). Data in the graphs represent the mean ± s.e.m. * P < 0.05; *Lean* vs. *NAFLD* and *NAFLD* vs. N*ASH*; ** P < 0.01; *Lean* vs. *Obese*; *** P < 0.001; *scWAT* vs. *vWAT*.

Identification of small molecule inhibitors targeting Them1 enzymatic activity necessitated the development of an Acot activity assay suitable for a HTS. Preliminary attempts using a colorimetric-based assay (Bernson, 1976) proved challenging, due to a lack of sensitivity and reproducibility upon scaling down to the requisite 384-well microplate-based format. Contributing factors were assay conditions that required microgram quantities of recombinant protein, cross reactivity with Ellman’s reagent with library compounds and low Z’ factors. These technical constraints were addressed by the development of a highly sensitive 384-well microplate-based fluorometric Acot activity assay in which free CoA liberated by Them1 activity covalently bound to a non-fluorescent reagent, generating a fluorescent product. The assay proved suitable for the HTS with consistently high Z’ factors among microplates and reproducibility on 2 independent days in pilot small molecule screens.

The helix-grip fold of the Them1 lipid-sensing START domain forms a hydrophobic tunnel that accommodates a lipid molecule (Schrick et al., 2004). Binding of long chain fatty acids allosterically enhances the enzymatic activity of the thioesterase domains (Han and Cohen, 2012; Tillman et al., 2020). The notion that an allosteric START domain-dependent small molecule inhibitor could be developed is in line with the observation that the lipid metabolite 18:1 lysophosphatidylcholine inhibits Them1 activity allosterically by binding the START domain (Tillman et al., 2020).

Based on our prior observation of allosteric regulation of Them1 activity through the START domain (Tillman et al. 2020), we selected compounds **U1** and **W1** from the HTS. This choice was further supported based on compound potency (Bickerton et al., 2012; Holdgate et al., 2017). For compound **W1**, the possibility of irreversible inhibition by covalent acetylation was discounted based on reversibility in response to dialysis. In keeping with our findings *in vitro* (i.e. enhanced potency and selectivity without covalent modification of Them1), compound **W1** reduced rates of glucose production in primary hepatocytes cultured from *L-Them1Tg* mice to levels comparable to wild type control hepatocytes. By contrast, this small molecule was apparently ineffective at inhibiting Them1 in primary brown adipocytes. We cannot exclude the possibility of compound **W1** instability in primary brown adipocytes, which could have led to the hydrolysis of the acetate side chain, resulting in an inactive product.

Because it exhibited activity in both primary brown adipocytes and hepatocytes, Compound **U1** was selected for more detailed SAR strategies through substitution or expansion of critical functional groups within the chemical structure. Values of OCR in primary brown adipocytes cultured from *A-Them1Tg* mice treated with structural analogs and derivatives of compound **U1** either remained suppressed or were restored to levels comparable to wild type control brown adipocytes. Because these effects were observed within an acute treatment period of 30 min, it is likely that compound **U1** and its structural analogs function through reversibly bound protein-inhibitor complexes to inhibit Them1 activity, rather than irreversible covalent inhibition (Singh et al., 2011), in keeping with observations *in vitro* using dialysis.

The current study identified several potent allosteric small molecule inhibitors with selectivity toward inhibiting Them1 activity through the START domain. These allosteric small molecule inhibitors should be well positioned for development as lead therapeutic compounds with promise for the management of NAFLD.

## ACKNOWLEDGEMENTS

The authors thank the Fisher Drug Discovery Resource Center at the Rockefeller University for providing equipment and assistance in the high-throughput and counter small molecule screens, and microscale thermophoresis and seahorse assays, the Bioexpression and Fermentation Facility at the University of Georgia for assistance in the production of recombinant Them1, the Proteomics and Metabolomics Core at Weill Cornell Medicine for verification of recombinant Acot isoforms and chemical compounds by high-resolution mass spectrometry, the Metabolic Phenotyping Center at Weill Cornell Medicine and the Seahorse Core at Brigham and Women’s Hospital for assistance with OCR measurements, the Mouse Genetics Core Facility at Memorial Sloan Kettering Cancer Center for assistance in the generation of the conditional transgenic tissue-specific Them1 overexpression, Drs. Doug Mashek (University of Minnesota; Minneapolis, Minnesota) and Baran Ersoy (Weill Cornell Medicine; New York, NY) for sharing the ds-RED-Acot1 and pMAL-c5X-Acot9 plasmids, respectively. This work was supported by the National Institute of Diabetes and Digestive and Kidney Diseases (R37 DK048873, R01 DK103046, R01 DK056626 and T32 DK116970 to D.E.C.), a Daedalus Fund for Innovation of Weill Cornell Medicine (to D.E.C.) and a Sanofi Innovation Awards Program Funding Award (to D.E.C.); C.S.K. acknowledges support by the Multidisciplinary Training in Gastroenterology & Hepatology Postdoctoral Fellowship from NIH T32 DK116970 (to D.E.C.); X.L. was supported by an American Heart Association Career Development Award [848388]. Support from the Weill Cornell Metabolic Phenotyping Center is gratefully acknowledged. The content is solely the responsibility of the authors and does not necessarily represent the official views of the National Institutes of Health.

## AUTHOR CONTRIBUTIONS

Conceptualization, C.S.K. and D.E.C.; Methodology, C.S.K., L.R.E. R.S.L., C.A., X.L., M.A., Y.X., X.X., M.C.T., J.D.G., J.F.G., E.A.O. and D.E.C.; Validation, C.S.K., L.R.E., R.S.L, C.A..; Investigation, C.S.K. L.R.E. R.S.L.; C.A., X.L., M.A., Y.X., X.X., M.C.T. and Y.L.; Resources, J.F.G. E.A.O. and D.E.C.; Writing – Original Draft, C.S.K. and D.E.C.; Writing – Review & Editing, C.S.K., L.R.E., R.S.L., C.A., X.L., M.A., Y.X., X.X., M.C.T., Y.L., J.D.G., J.F.G., E.A.O. and D.E.C.; Visualization, C.S.K. and D.E.C.; Supervision, E.A.O. and D.E.C.; Funding Acquisition, D.E.C.

## STAR METHODS

### RESOURCE AVAILABILITY

#### Lead contact

Requests for reagents and resources should be directed to the Lead Contact, David E. Cohen.

#### Materials availability

All data reported in this paper will be shared by the lead contact upon request.

#### Data and code availability

This paper analyzes existing, publicly available data. These accession numbers for the datasets are listed in the key resources table. This paper does not report original code. Any additional information required to reanalyze the data reported in this paper is available from the lead contact upon request.

### EXPERIMENTAL MODEL AND SUBJECT DETAILS

#### Animals

Generation of conditional transgenic tissue-specific Them1 overexpression (wild type) mice was previously described with modifications (Madisen et al., 2010). A cDNA encoding mouse Them1 was fused to a C-terminal FLAG-tag (FLAG) as a marker for Cre recombination and cloned into a Rosa26 expression vector consisting of a CMV promoter, a STOP cassette flanked by loxP sites and a polyA tail (pA). The plasmid was then linearized and microinjected into pronuclei of eggs from C57BL/6 female mice (Mouse Genetics Core Facility; Memorial Sloan Kettering Cancer Center; New York, NY). Conditional transgenic adipose tissue-specific Them1 overexpression (*A*-*Them1Tg)* mice were generated by crossing wild type mice to transgenic mice expressing Cre recombinase driven by the adiponectin gene promoter (B6.FVB-Tg(Adipoq-Cre)1Evdr/J; Strain #: 028020; Jackson Laboratory; Bar Harbor, ME) on a congenic C57/BL6 background (Eguchi et al., 2011). Mice with liver-specific Them1 overexpression (*L-Them1Tg*) and wild type controls, along with Them1 knockout (*Them1^-/-^*) controls were generated by intravenously injecting 6- to 14-w old wild type, C57BL/6 and *Them1^-/-^* mice (Zhang et al., 2012), respectively, with adeno-associated virus 8 (AAV8) expressing Cre recombinase driven by the human thyroid hormone-binding globulin (TBG) promoter (AAV8.TBG.Cre). Adiponectin- and TBG-Cre mediated recombination resulted in excision of the STOP cassette to bring Them1 under control of the CMV promoter in adipose tissue and hepatocytes, respectively. Litters were genotyped for integration of the Them1 transgene by PCR analysis from ear genomic DNA. Tissues were harvested, immediately snap frozen in liquid nitrogen and stored at -80 °C. Mice were same sex housed in mixed genotype groups (3 to 5 mice per cage) in a barrier facility on a 12 h light/dark cycle. Animal use and euthanasia protocols were performed using approved guidelines by Weill Cornell Medical College.

#### Cultures of primary brown adipocytes

Primary brown adipocytes from 4- to 6-w old wild type and *A-Them1Tg* mice were isolated, cultured and differentiated as described (Okada et al., 2016). In brief, BAT tissues harvested from 5- to 7-w old mice were pooled, minced and digested with collagenase B (Sigma-Aldrich; St. Louis, MO), and dispersed in growth medium [DMEM/F12 containing 4.5 g/L glucose, 0.1 mM pyruvate, 10 mM HEPES (Thermo Fisher; Waltham, MA) supplemented with 1% penicillin/streptomycin, 1% GlutaMAX (Thermo Fisher; Waltham, MA) and 20% fetal bovine serum]. Cells were seeded at 2,000 per well into XF96 cell culture plates (Seahorse Bioscience; North Billerica, MA) pre-coated with rat tail collagen (Sigma Aldrich; St. Louis, MO). Upon achieving confluence, pre-adipocytes were induced to differentiate for 8 d by culturing for the first 48 h in differentiation medium [DMEM/F12 containing 4.5 g/L glucose, 0.1 mM pyruvate, 10 mM HEPES (Thermo Fisher; Waltham, MA) supplemented with 1% penicillin/streptomycin, 1% GlutaMAX (Thermo Fisher; Waltham, MA), 10% fetal bovine serum, bovine insulin (5 μg/mL; Thermo Fisher; Waltham, MA) and rosiglitazone (1 μM; Sigma Aldrich; St. Louis, MO)] supplemented with dexamethasone (5 μM; Sigma Aldrich; St. Louis, MO) and 3-isobutyl-1-methylxanthine (0.5 mM; Sigma Aldrich; St. Louis, MO) and differentiation medium thereafter. On differentiation d 6 to 8, the adipocyte differentiation media was supplemented with L-(-)-norepinephrine (+) bitartrate (Calbiochem; EMD Millipore; Billerica, MA). Primary brown adipocytes were maintained in a cell culture incubator at 37 °C with 5% CO_2_.

#### Cultures of primary hepatocytes

Primary hepatocytes cultured were prepared from 6- to 14-w old wild type, *L-Them1Tg* and *Them1^-/-^* mice following anesthesia with ketamine and xylazine. Livers were perfused through the portal vein with 20 mL of liver perfusion medium (Thermo Fisher; Walham, MA) followed by 40 mL of liver digestion medium (Thermo Fisher; Walham, MA). Primary hepatocytes were gently disrupted from the liver capsule into hepatocyte wash medium (Thermo Fisher; Waltham, MA), filtered through a 70 μm cell strainer and then spun down at 30 x g for 4 min at 4 °C. Primary hepatocytes were cultured in William’s Medium E (Thermo Fisher; Waltham, MA) supplemented with 1% penicillin/ streptomycin and 10% fetal bovine serum. For the fatty acid oxidation and glucose production assays, primary hepatocytes were seeded at 2.5 x10^5^ cells per well into XF24 cell culture microplates (Seahorse Bioscience; North Billerica, MA) pre-coated with rat tail collagen (Sigma Aldrich; St. Louis, MO) or at 5 x10^5^ cells/well in 6-well Primaria plates (Corning Inc.; Corning NY), respectively. Cultured primary hepatocytes were maintained in a cell culture incubator at 37 °C with 5% CO_2_.

### METHOD DETAILS

#### Expression and purification of recombinant Acot isoforms

Human Them1 consists of two splice variants (Them1a and Them1b) that are distinguished by an additional 13 amino acids at the C-terminus of the ‘a’ isoform (Adams et al., 2001). For human Them1 (Them1) and a truncated Them1 containing the two thioesterase domains, but lacking the START domain (Them1-ι1START), a synthetic gene encoding Them1b (the human ortholog of mouse Them1) was codon optimized to achieve maximal recombinant protein expression (Thermo Fisher; Waltham, MA) and subcloned into a pET19b bacterial expression vector (Novagen, EMD Biosciences; Madison, WI), which introduced an in-frame N-terminal His-tag. Sufficient quantities of recombinant Them1 required for the HTS were obtained as a service of the University of Georgia Bioexpression and Fermentation Core Facility (Athens, GA). Cultures of *E. coli* Shuffle T7 competent cells (New England Biolabs, Ipswich, MA) transformed with pET19b-Them1 plasmid were grown to an A_600_ of ∼ 0.5 – 0.7 in terrific broth followed by induction of recombinant Them1 by the addition of 0.5 mM isoproyl β-D-thiogalactoside with 16 h of shaking (250 rpm) at 18 °C. The bacteria were harvested by centrifugation and then lysed with 20 mM Tris-Cl (pH 7.4), 500 mM NaCl, 150 mM imidazole and 1 mM β-mercaptoethanol. The soluble fraction following centrifugation of the bacterial lysate was purified by fast liquid protein chromatography (FPLC) using a HisTrap affinity column (HisTrap HP column; GE Healthcare, Waukesha, WI) after equilibration with 20 mM Tris-Cl (pH 7.4), 500 mM NaCl, 150 mM imidazole and 1 mM β-mercaptoethanol, and then washed with the same buffer. Recombinant Them1 was eluted from the HisTrap HP column using 20 mM Tris-Cl (pH 7.4), 500 mM NaCl, 500 mM imidazole and 1 mM β-mercaptoethanol. To further increase purity, recombinant Them1 was applied to a HisTrap HP affinity column (HisTrap HP column; GE Healthcare, Waukesha, WI) and re-purified as described above followed by dialysis into buffer containing 20 mM Tris-Cl (pH 7.4), 500 mM NaCl and 10% glycerol using a slide-a-lyzer dialysis cassette (Thermo Fisher; Waltham, MA). Purity of recombinant Them1 was analyzed by SDS-PAGE followed by Coomassie Brilliant Blue staining. The concentration of recombinant Them1 was determined according to the molar extinction coefficient at 280 nm, which was calculated based on the amino acid sequence (www.expasy.org). Single use aliquots of recombinant Them1 were stored at -80 °C until use to prevent protein precipitation during purification. The pH values of all buffers were adjusted to remain at least two units removed from the pI of recombinant Them1.

Molecular cloning, and bacterial expression and purification of recombinant Acot isoforms were prepared as previously described with modifications (Han and Cohen, 2012). The open reading frames of mouse Acot1 (Acot1), mouse Them1 and human Acot12 (Acot12) were amplified by PCR using the template plasmids ds-RED-Acot1, pCMV-SPORT6-Them1 and pcDNA-Acot12, respectively. The open reading frames of mouse Acot2 (Acot2) and human Acot13 (Acot13) were amplified by PCR using mouse liver tissue. The genes were subcloned into pET19b or pET29b bacterial expression vectors (Novagen, EMD Biosciences; Madison, WI), which introduced an in-frame N- or C-terminal His-tag, respectively. Maltose-binding protein (MBP)-tagged human Acot9 (Acot9) plasmid was prepared as previously described (Tillander et al., 2014). These plasmids were transformed into *E. coli* strains BL21 (DE3) (New England Biolabs; Ipswich, MA) or Shuffle T7 competent cells (New England Biolabs, Ipswich, MA), grown to an A_600_ of 0.5 – 0.7 in terrific broth and then induced by the addition of 0.5 mM isoproyl β-D-thiogalactoside with 16 h of shaking (250 rpm) at 18 °C. The bacteria were harvested by centrifugation at 10,000 x g for 20 min at 4 °C, and then lysed with 20 mM Tris-Cl, (pH 7.4) 500 mM NaCl, 150 mM imidazole, 1 mM β-mercaptoethanol and ethylenediaminetetraacetic acid-free protease inhibitors (Sigma Aldrich, St. Louis, MO). Following centrifugation of the bacterial lysate at 20,000 x g for 20 min at 4 °C, recombinant Acot isoforms were purified by FPLC using a HisTrap affinity column (HisTrap HP column; GE Healthcare, Waukesha, WI) unless otherwise specified. The soluble fractions containing recombinant Acot isoforms were applied to the column after equilibration with 20 mM Tris-Cl (pH 7.4), 500 mM NaCl, 150 mM imidazole, 1 mM β-mercaptoethanol and ethylenediaminetetraacetic acid-free protease inhibitors (Sigma Aldrich, St. Louis, MO), and then washed with the same buffer. His-tagged recombinant Acot isoforms were then eluted from the HisTrap HP column using 20 mM Tris-Cl (pH 7.4), 500 mM NaCl, 500 mM imidazole, 1 mM β-mercaptoethanol and ethylenediaminetetraacetic acid-free protease inhibitors (Sigma Aldrich, St. Louis, MO). The soluble fraction containing MBP-tagged recombinant Acot9 was applied to a MBPTrap HP column after equilibration with 20 mM Tris-Cl, (pH 7.4) 200 mM NaCl, 1 mM EDTA and protease inhibitors (Sigma Aldrich, St. Louis, MO), washed with the same buffer and then eluted with 20 mM Tris-Cl (pH 7.4), 200 mM NaCl, 1 mM EDTA, 10 mM Maltose and protease inhibitors (Sigma Aldrich, St. Louis, MO). Following purification, all recombinant Acot isoforms were subjected to dialysis into buffer containing 20 mM Tris-Cl (pH 7.4), 500 mM NaCl and 10% glycerol using a slide-a-lyzer dialysis cassette (Thermo Fisher; Waltham, MA). Assessment of purity, protein concentration and storage of single use aliquots of recombinant Acot isoforms, and adjustment of buffer pH values were performed as described above for recombinant Them1.

Recombinant human Them1 START domain (START; amino acids 339 – 594 of Them1b) was prepared as previously described (Tillman et al. 2020). A synthetic gene encoding the START domain was cloned into a pNIC28-Bsa4 bacterial expression vector (Novagen, EMD Biosciences; Madison, WI), which introduced an in-frame N-terminal His_6_ fusion containing a tobacco etch virus protease cleavage site to facilitate tag removal. The pNIC28-Bsa4-START domain plasmid was co-transformed into *E. coli* BL21 (DE3) (New England Biolabs; Ipswich, MA) competent cells with a pG-Tf2 vector (encoding groES-gorEL-tig chaperones) and then grown to an A_600_ of 0.5 – 0.7 in terrific broth. Under conditions of shaking (250 rpm) at 18 °C, chaperones were induced by the addition of 5 ng/mL tetracycline HCl for 60 min followed by START domain induction upon the addition of 0.5 mM isoproyl β-D-thiogalactoside for ∼ 18 h. The bacteria were harvested by centrifugation, and then lysed by sonication with 20 mM Tris-Cl (pH 7.4), 500 mM NaCl, 25 mM imidazole, 5% glycerol, lysozyme, Dnase A, 0.1% Triton X-100, 5 mM β-mercaptoethanol and 100 μM phenylmethylsulfonyl fluoride. Following centrifugation of the bacterial lysate, recombinant START was purified by FPLC using a His affinity column (HisTrap HP column; GE Healthcare; Waukesha, WI). The soluble fraction containing recombinant START was applied to the column after equilibration with 20 mM Tris-Cl (pH 7.4), 500 mM NaCl, 25 mM imidazole and 5% glycerol, and then washed with the same buffer. Recombinant START was eluted from the HisTrap HP column using 20 mM Tris-Cl (pH 7.4), 500 mM NaCl, 500 mM imidazole and 5% glycerol followed by tobacco etch virus protease-mediated His-tag cleavage at 4 °C overnight with simultaneous dialysis into 20 mM Tris-Cl (pH 7.4), 500 mM NaCl and 5% glycerol. Recombinant START was further purified using a Superdex column (HiLoad 16/60 Superdex 75 column; GE Healthcare; Waukesha, WI) in 20 mM Tris-Cl (pH 7.4), 500 mM NaCl and 5% glycerol. Assessment of purity, protein concentration and storage of single use aliquots of recombinant START were performed as described above for recombinant Them1.

#### Small molecule library

The compound library of the Fisher Drug Discovery Resource Center at Rockefeller University consisted of approximately 361,000 compounds including chemical compounds in parenthesis from the following companies: Amri (50,000; Albany, NY), AnalytiCon (700; Lichtenfels, Germany), BioFocus DPI (10,150; Essex, United Kingdom), Chem-X-Infinity (4,000; Palo Alto, CA), ChemBridge Corporation (65,638; San Diego, CA), ChemDiv (126,000; San Diego, CA), Enamine (79,921; Monmouth Junction; NJ), Edelris (2,000; Lyon, France), Greenpharma (240; 45100 Orlans, France), Life Chemicals (30,272; Ontario, CA) and SPECS (4,051; Bleiswijkseweg 55, The Netherlands). Small molecules generally adhered to Lipinski’s rules and contained a low proportion of known toxicophores and unwanted functionalities. A test library of 1,056 pharmacologically active compounds (LOPAC; Sigma Aldrich; St. Louis, MO) was used as a pilot small molecule screen. Commercially available structural analogs to the parent compounds **U1** (i.e. **U2**, **U3**, **U4** and **U5**) and **W1** (i.e. **W2**, **W3**, **W4** and **W5**) were purchased from MolPort (Beacon, NY). Compound **U1** structural derivatives **U6** - **U17** (Supplemental methods) were chemically synthesized as a service of WuXi App Tec (Beijing, China).

#### Acot activity assay

A 96-well fluorometric Acot activity assay was developed using a commercial kit detecting free thiols (Thermo fisher; Waltham, MA), and then scaled down and optimized in polystyrene 384-well microplates. Free CoA liberated by Acot activity covalently binds to a non-fluorescent detection reagent, resulting in the generation of a fluorescent product. Upon the addition of C14-CoA, time-dependent increases in fluorescence intensity reflected free CoA covalently bound to detection reagent.

The HTS was carried out in 20 μL final volumes in black 384-well flat bottom polystyrene microplates (GreinerBio One International AG; Kremsmünster, Austria) at 22 °C. A single 1000 x g, 30 s centrifugation step was performed upon the addition of any reagents to ensure that the reaction mixture collected at the bottom of the well. Reactions comprised of recombinant Them1 (125 nM), C14-CoA (25 μM) and compounds (12.5 μM). Assay buffer (Thermo Fisher; Waltham, MA) was dispensed into wells using a liquid handling Thermo Multidrop Combi dispenser (Thermo Fisher; Waltham, MA). Five mM stock compounds dissolved in DMSO were dispensed with a Janus 384 MDT NanoHead (PerkinElmer; Waltham, MA). Preliminary experiments demonstrated that 0.5% DMSO did not interfere with liberation of free CoA from C14-CoA. On each plate, recombinant Them1 was added to wells in columns 1 to 23 using a Multidrop Combi dispenser (Thermo Fisher; Waltham, MA), while Assay buffer (Thermo Fisher; Waltham, MA) was added in column 24. Reactions were initiated by the addition of C14-CoA and were incubated for 60 min at 22 °C, and then stopped by the addition of 0.1 M HCl. Upon adding the detection reagent, reactions were incubated for 30 min. Fluorescence was measured at 510 nm emission and 390 nm excitation using a NEO plate reader (Bio Tek; Winooski, VT). In the wells serving as (+) or (-) controls, recombinant Them1 or compound, respectively, were substituted with buffer. Acot activity was normalized to the (+) control (column 23) and (-) control (column 24) as follows: NPI (%) = 100 x (Relative fluorescent units (RFU)_average (-) control_ – RFU_sample_) / (RFU_average (-) control_ -RFU_average (+) control_). IC_50_ values were determined by fitting the calculated values of NPI to a sigmoidal curve using GraphPad Prism GraphPad Software; San Diego, CA).

Compounds that inhibited Them1 activity with a NPI ≥ 30 were re-tested using the Acot activity assay by concentration response experiments to determine IC_50_ values. Compounds exhibiting IC_50_ values ≤ 20 μM were further tested for selectivity using a counter screen against other Acot isoforms (i.e. Acot1, Acot2, Acot9 and Acot13), as well as Them1-1′START. Reactions were performed in 384-well microplates at 22 °C for 60 min with recombinant Acot isoforms (125 nM) and C14-CoA (25 μM). Compounds **U1** and **W1** were tested for activity targeting mouse Them1 using the Acot activity assay by concentration response experiments to determine IC_50_ values. Reactions were performed in 384-well microplates at 22 °C for 60 min with recombinant mouse Them1 (125 nM) and C14-CoA (25 μM).

The most promising compounds showing selectivity toward Them1 activity were ordered from available vending sources, dissolved in DMSO at 50 mM and tested for concentration-dependent inhibition. IC_50_ values derived from the Acot activity assay were calculated using GraphPad Prism (San Diego, CA) from 3 replicate experiments. The Z’ factor, which is a measure of suitability of an assay for the HTS, was calculated as Z’(60’) = 1-3 (S.D._-_ + S.D._+_) / [F(60’)_-_ -F(60’)_+_] (Zhang et al., 1999).

#### Kinetic characterization of Acot activity

Steady-state kinetic parameters were determined as previously described with modifications (Han and Cohen, 2012). Acot activity was determined in black 384-well flat bottom polystyrene microplates (GreinerBio One International AG; Kremsmünster, Austria) for 60 min at 22 °C as a function of time after mixing recombinant Acot isoform (125 nM) with substrate C14-CoA (25 μM). Initial rates (V_0_) were determined using GraphPad Prism (GraphPad Software; San Diego, CA). Concentrations of C14-CoA were varied to create saturation curves, and values of V_0_ were fitted to the Michaelis-Menten equation using GraphPad Prism GraphPad Software; San Diego, CA). Nonlinear analysis of the Michaelis-Menten equation provided satisfactory curve-fits with an average R^2^ = 0.99 and a minimum R^2^ > 0.95. K_cat_ values were determined by fitting the calculated values of V_0_ and substrate concentrations at 0, 15, 30, 45, 60 and 90 min timepoints to a sigmoidal concentration-response curve using GraphPad Prism (GraphPad Software; San Diego, CA).

#### Assay for reversibility of Them1 inhibition

Them1 activity was first measured in reactions consisting of recombinant Them1 (125 nM), C14-CoA (25 μM) and compounds (125 μM) incubated in 384-well microplates for 60 min at 22 °C. Reactions were dialyzed at 4 °C in buffer containing 20 mM Tris-Cl (pH 7.4) and 500 mM NaCl using slide-a-lyzer dialysis cassettes (Thermo Fisher; Waltham, MA) with the pH adjusted to remain at least 2 units removed from the pI of recombinant Them1. Three buffer changes were performed over the extent of the 18 h dialysis. Post-dialysis reactions were incubated with fresh C14-CoA (25 μM) for 60 min at 22 °C. Ebselen (Sigma Aldrich; St. Louis, MO) was used as a (-) control.

#### Microscale thermophoresis

Reactions labeling recombinant full-length Them1, Them1-ΔSTART, START or Acot12 with Monolith RED-Tris-NTA (NanoTemper Technologies; San Francisco, CA) were performed according to the manufacturer’s instructions in buffer containing PBS plus 0.05% Tween-20 for 60 min at 22 °C with a molar dye: protein ratio = 1:1. Compounds starting at 50 mM were serially diluted in DMSO. Working concentrations of compounds were prepared by diluting in microscale thermophoresis buffer. Reactions consisting of 100 nM of fluorescently labeled recombinant protein (full-length Them1, Them1-ΔSTART, START or Acot12) and compounds were loaded onto standard Monolith NT.115 Capillaries (NanoTemper Technologies; San Francisco, CA). Fluorescence was measured using a Monolith NT.115 instrument (NanoTemper Technologies; San Francisco, CA) at 22 °C with instrument parameters adjusted to 40% LED power and medium microscale thermophoresis power. Data were fitted by non-linear regression in GraphPad Prism (GraphPad Software; San Diego, CA).

#### Cytotoxicity assay

Cytotoxicities of compounds were determined using a CellTiter-Glo® Luminescent Cell Viability Assay, which measures concentrations of ATP liberated from viable cells (Promega; Madison, WI). Primary brown adipocytes cultured from C57BL/6 mice were seeded at 2,000 per well into white 384-well transparent bottom opaque microplates (GreinerBio One International AG; Kremsmünster, Austria) pre-coated with rat tail collagen (Sigma Aldrich; St. Louis, MO) at 22 °C and treated for 48 h with DMSO (Control) or compounds (250 μM to 7.7 nM) in growth medium. Tamoxifen (50 μM) was used as a (+) control. CellTiter-Glo reagent (10 μL) was added to each well and incubated on an orbital shaker for 10 min at 22 °C to induce cell lysis and stabilize the luminescent signal. Relative luminescent units (RLU) were determined using a NEO plate reader (BioTek, Winooski, VT).

#### Critical aggregation concentration

Critical aggregation concentrations of compounds were determined using dynamic light scattering in black 96-well flat bottom polystyrene microplates (GreinerBio One International AG; Kremsmünster, Austria) at 22 °C using a DynaPro plate reader (Wyatt Technology; Santa Barbara, CA). 50 mM stock compounds dissolved in pure DMSO were serial diluted 2-fold (250 to 62.5 μM) in PBS and then spun down at 14,000 x g for 10 min at 22 °C followed by filtration through a 0.22 μm filter unit. Aggregation of compounds were assessed by measuring the amplitude of the autocorrelation intensity calculated from the average of 3 to 5 acquisitions per replicate.

#### Oxygen consumption rates (OCR)

In primary brown adipocytes on d 8 of differentiation, values of OCR were measured in the presence or absence of compounds using a Seahorse XF96 extracellular flux analyzer (Agilent Technologies; Santa Clara, CA). Primary brown adipocytes were treated for 30 min with DMSO (Control) or compounds in serum-free growth medium followed by incubation in the absence of CO_2_ for 60 min at 37 °C in Krebs-Henseleit buffer (pH 7.4) containing 0.45 g/L glucose, 111 mM NaCl, 4.7 mM KCl, 2 mM MgSO_4_^-^7H_2_O, 1.2 mM Na_2_HPO_4_, 5 mM HEPES and 0.5 mM carnitine (Sigma-Aldrich; St. Louis, MO). OCR values were measured before and after the exposure of cells to NE (1 μM) and normalized with total live cell count calculated through staining with NucRed Live probe (Thermo Fisher; Waltham, MA) measured at 715 nm emission and 623 nm excitation using a SpectraMax i3X plate reader (Molecular Devices; San Jose; CA).

In primary hepatocytes, values of OCR were measured using a Seahorse XF24 extracellular flux analyzer (Agilent Technologies; Santa Clara, CA). Following 4 to 5 h of incubation in William’s Medium E (Thermo Fisher; Waltham, MA) supplemented with 10% FBS and 1% penicillin-streptomycin, primary hepatocytes were treated for 30 min with DMSO (Control) or compounds in serum-free Medium 199 (Thermo Fisher; Waltham, MA) followed by incubation in the absence of CO_2_ for 60 min at 37 °C in Krebs-Henseleit buffer (pH 7.4) containing 0.45 g/L glucose, 111 mM NaCl, 4.7 mM KCl, 2 mM MgSO_4_^-^7H_2_O, 1.2 mM Na_2_HPO_4_, 5 mM HEPES and 0.5 mM carnitine (Sigma-Aldrich; St. Louis, MO). OCR values were measured before and after the exposure of cells to palmitic acid conjugated with fatty acid-free BSA (300 μM) and normalized with total live cell count calculated through staining with NucRed Live probe (Thermo Fisher; Waltham, MA) measured at 715 nm emission and 623 nm excitation using a SpectraMax i3X plate reader (Molecular Devices; San Jose; CA).

#### RNA extraction and analysis of gene expression

Total RNA was extracted from mouse WAT, BAT and kidney and primary brown adipocytes cultured from wild type and *A-Them1Tg* mice using QIAzol lysis reagent (Qiagen; Valencia, CA), and used to synthesize cDNA with a High-Capacity cDNA Reverse Transcription Kit (Applied Biosystems; Foster City, CA). Gene expression was analyzed with quantitative real-time PCR assays using Power SYBR Green Mix (Applied Biosystems; Foster City, CA). Real-time PCR assays were performed in triplicate with a total reaction volume of 25 μL containing 500 nM concentrations of each primer and cDNA (25 ng). mRNA expression levels were normalized to the housekeeping gene *18S*.

#### Glucose production

Rates of glucose production were determined as previously described with modifications (Kawano et al., 2014). Primary hepatocytes cultured from wild type, *L-Them1Tg* and *Them1^-/-^* mice were serum-starved in Medium 199 (Thermo Fisher; Waltham, MA) for 16 h and then washed twice with PBS. Cells were then incubated for 6 h in glucose-, L-glutamine- and phenol red-free DMEM (Thermo Fisher; Waltham, MA) supplemented with compounds, 2 mM sodium pyruvate and 20 mM sodium lactate. Glucose concentrations in the media were measured enzymatically through a commercial kit according to the manufacturer’s instructions using a NEO plate reader (BioTek, Winooski, VT).

#### Immunoblot analysis

Tissue extracts from mouse brain, heart, lung, WAT, skeletal muscle (muscle) and kidney, and total cellular extracts from primary brown adipocytes and hepatocytes were prepared in 10 mM Tris-HCl (pH 7.4), 150 mM NaCl, 1% Nonidet P-40, 1 mM phenylmethylsulfonyl fluoride, 1 mM EDTA, 1 mM NaF, 0.25% sodium deoxycholate, and 10% glycerol, supplemented with protease and phosphatase inhibitors (Thermo Fisher; Waltham, MA). Protein extracts were separated on 10 to 12% polyacrylamide gels and transferred onto nitrocellulose membranes (Protan, Schleicher, and Schuell Bioscience; Dassel, Germany). Membranes were blocked in Tris-buffered saline with Tween-20 (0.05 M Tris-HCl (pH 7.4), 0.2 M NaCl, and 0.1% Tween-20) containing (5% wt/vol) nonfat dried skim milk. Membranes were then immunodecorated with primary antibodies against mouse Them1 (Han and Cohen, 2012; Zhang et al., 2012), FLAG (BioLegend; San Diego, CA) and Hsp90 (Santa Cruz Biotechnology; Santa Cruz, CA), and diluted in blocking solution. Signals were developed with goat anti-rabbit or goat anti-rat secondary antibodies (Thermo Fisher; Waltham, MA), and visualized with the ProteinSimple system (ProteinSimple; San Jose, CA).

#### Quantification and statistical analysis

Data were analyzed by a mixed model using the fit model procedure of JMP Pro 11.0 statistical software (SAS Institute; Cary, NC). For experiments measuring OCR in primary brown adipocytes and hepatocytes and corresponding values of area under the curve (AUC), data were analyzed by two-way ANOVA accounting for genotype and compound. For analyses measuring *Them1* mRNA abundance in human tissues (BAT, WAT and liver), data were analyzed by pair-wise comparisons accounting for physiological state or tissue). For the reversibility washout experiments, data were analyzed by pair-wise comparisons accounting for dialysis. For experiments measuring *Them1* mRNA abundance in tissues from wild type and *A-Them1Tg* mice, data were analyzed by one-way ANOVA accounting for genotype. Statistical significance was determined using two-tailed unpaired Student t-test or two-way ANOVA followed by Tukey’s post-hoc test for comparisons between two or three groups. Differences were considered significant at *P* < 0.05. All statistical analyses were performed using JMP Pro 11.0 statistical software.

## Supplemental figures

**Supplemental figure 1.**
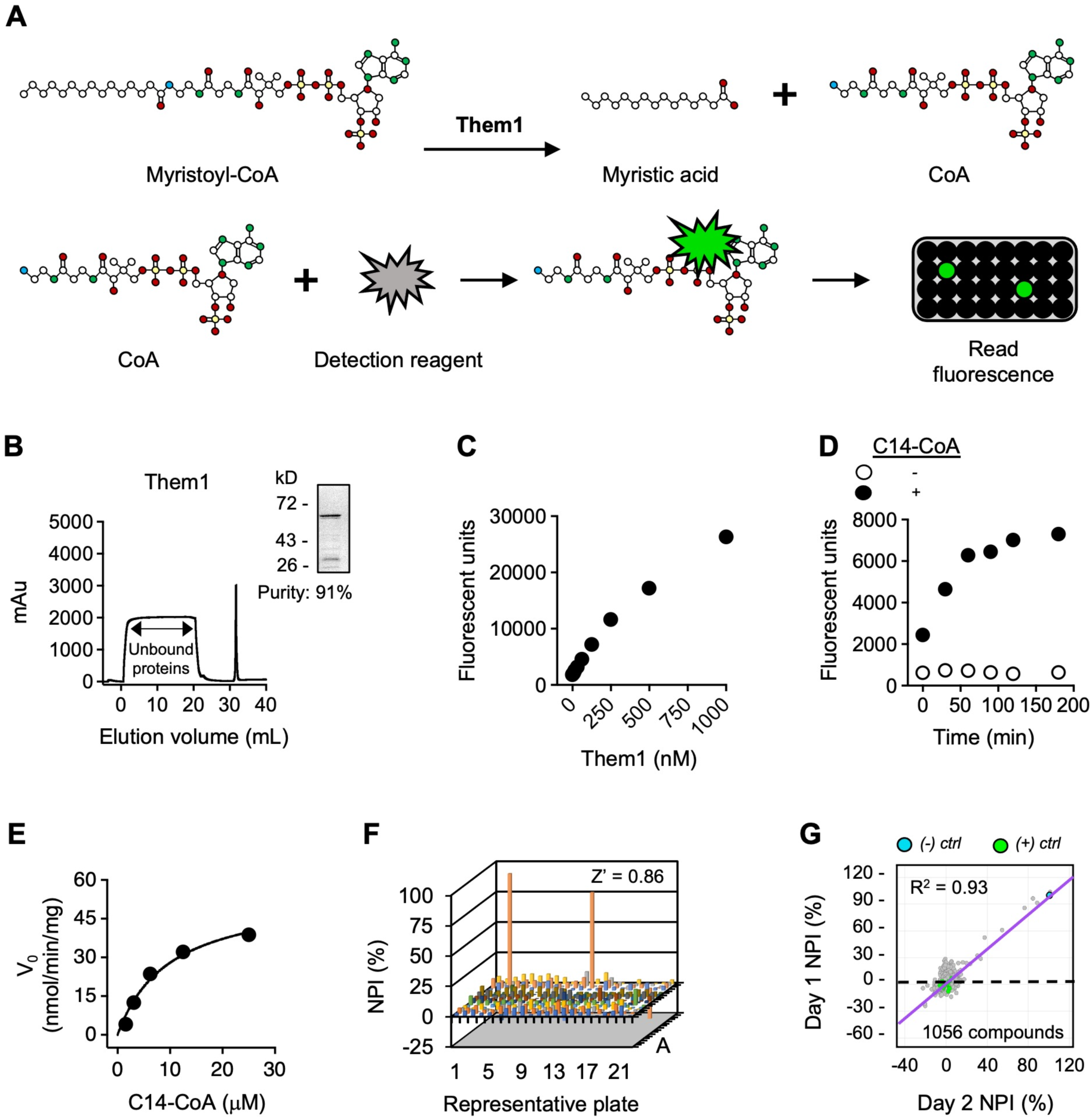
Optimization of an Acot activity assay for identification of small molecule inhibitors targeting Them1 activity. **(A)** Schematic showing Them1-mediated de-activation of myristoyl (C14)-CoA into myristic acid plus CoA. Free CoA liberated by Them1 activity covalently binds to a non-fluorescent detection reagent, generating a fluorescent product. **(B)** Recombinant His-tagged human Them1 (Them1) purified from Shuffle T7 competent cells on a HisTrap HP column (Black line). *Upper right panel*, SDS-PAGE of Them1 (67 kD). **(C)** Recombinant Them1 and C14-CoA (25 μM) reactions (black dots). **(D)** Reactions consisting of Them1 and C14-CoA (white dots, 0 μM; black dots, 25 μM) were incubated for the indicated timepoints. **(E)** Saturation curve of Them1 and C14-CoA reactions (black dots) were used to determined steady state kinetic constants. **(F)** A pilot small-molecule screen was performed using the library of pharmacologically active compounds (LOPAC, 12.5 μM, 1,056 compounds) with Them1 and C14-CoA as substrate. Compounds were considered as potential inhibitors when Them1 activity was inhibited based on a normalized percent inhibition (NPI) ≥ 30. A representative plate with Z’ factor and 2 compounds fulfilling the inhibitor criteria are shown. Negative values are attributable to the absorbance of a minority of compounds. **(G)** Reproducibility of the pilot small molecule screen was assessed by NPI values of compounds over 2 independent days. Correlation between d 1 and 2 is indicated by the purple line; R^2^ = 0.93. Data represent the mean (s.e.m.) of triplicate determinations. Where not visible, standard error bars are contained within the symbol sizes.

**Supplemental figure 2:**
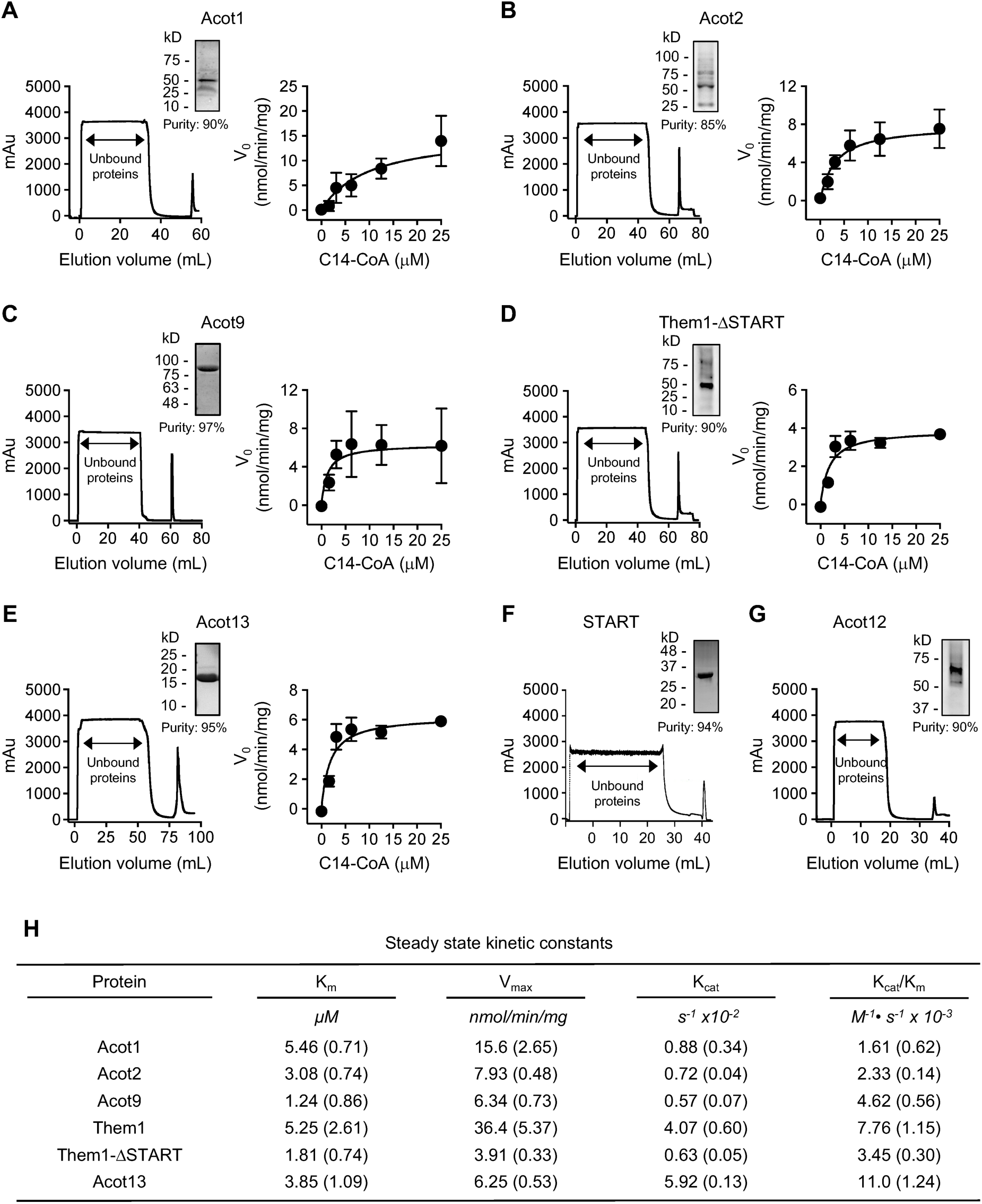
Recombinant Acot purification and steady state kinetic constants. **(A-G)** *Acot isoform panels*: *Left panels*, recombinant Acot isoform enzymes including type I [Acot1, Acot2] and type II [Acot9, Acot12 and Acot13], as well as a truncated Them1 either containing only the 2 thioesterase domains but lacking the START domain (Them1-ΔSTART) or the START domain but lacking the 2 thioesterase domains (START) were purified from Shuffle T7 or BL21 (DE3) *E. coli* competent cells on a HisTrap HP column unless otherwise specified. Recombinant START was further purified using a Superdex 75 column. Recombinant Acot9 was purified on a MBPTrap HP column. *Upper right panel*, SDS-PAGE of recombinant Acot isoform enzymes. *Right panel*, saturation curves of recombinant Acot isoform enzymes and myristoyl (C14)-CoA reactions (black dots) were used to determine steady state kinetic constants. Where not visible, standard error bars are contained within the symbol sizes. (**H**) *Table*, steady state kinetic constants (K_m_, V_max_, K_cat_ and K_cat_/K_m_) were determined as described in the text. Data represent the mean (s.e.m.) of triplicate determinations.

**Supplemental figure 3.**
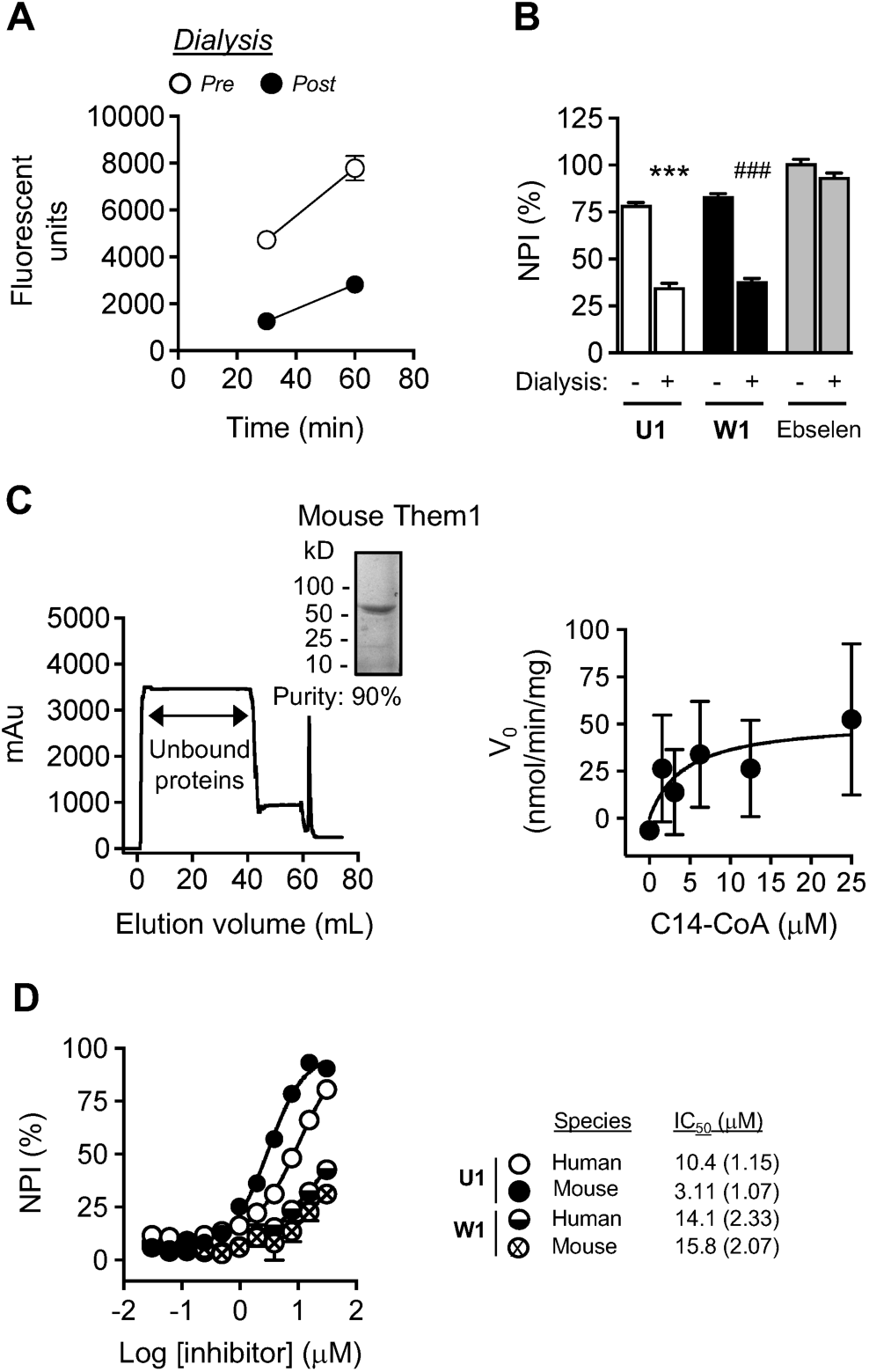
Them1 inhibitors bind reversibly and similarly inhibit mouse and human enzymes. Pre-dialysis reactions were performed with recombinant His-tagged human Them1 (Them1) and myristoyl (C14)-CoA. Reactions subjected to an overnight dialysis were incubated with C14-CoA. (**A**) Time course of Pre- and Post- dialysis reactions for the indicated timepoints. (**B**) Pre- and Post-reactions consisting of compounds **U1**, **W1** or Ebselen [(-) control]. Each graph is representative of four independent experiments. (**C**) *Left panels*, recombinant His-tagged mouse Them1 was purified from BL21 (DE3) *E. coli* competent cells on a HisTrap HP column. *Upper right panel*, SDS-PAGE of recombinant mouse Them1. *Right panel*, saturation curve of recombinant mouse Them1 (125 nM) and C14-CoA reactions (black dots) was used to determine steady state kinetic constants. (**D**) All reactions were performed with either recombinant mouse Them1 or recombinant human Them1. Acot activities quantified as IC_50_ values are shown. Data represent the mean ± s.e.m. of triplicate determinations. Where not visible, standard error bars are contained within the symbol sizes. ***, P<0.001; *Pre-* (**U1**) vs. *Post-* (**U1**); ###, P<0.001; *Pre-* (**W1**) vs. *Post-* (**W1**).

**Supplemental figure 4.**
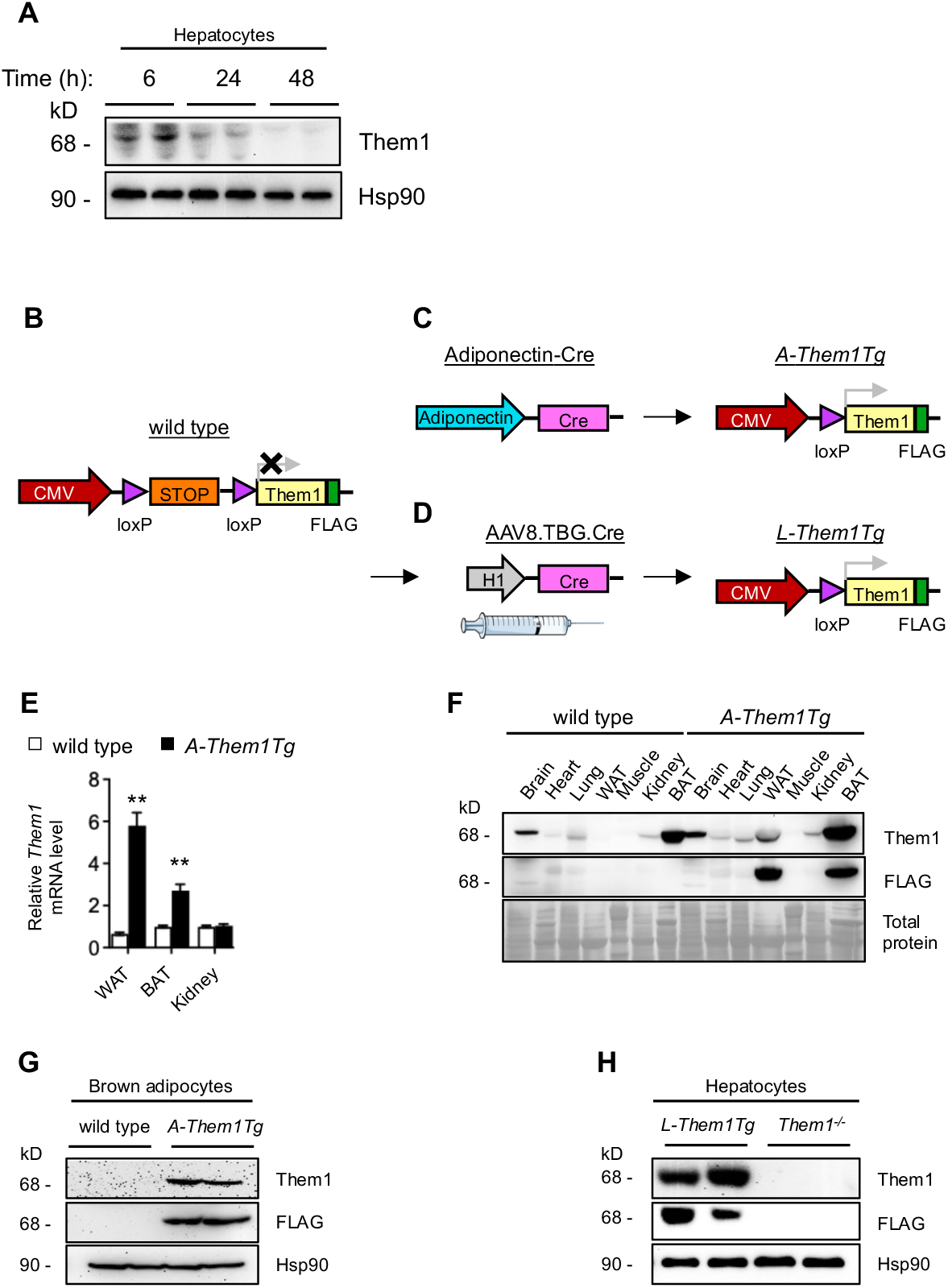
Generation of transgenic adipose tissue (*A-Them1Tg*) and liver (*L-Them1Tg*)-specific Them1 overexpression mice. **(A)** Protein expression levels of Them1 in primary hepatocytes were analyzed as functions of time in culture, with Hsp90 utilized to control for unequal loading. **(B)** A mouse Them1 (yellow boxes) cDNA fused to a C-terminal FLAG-tag (green; pink box) as a marker for Cre recombination was cloned into a Rosa26 expression vector consisting of a CMV promoter (red arrows) and a STOP cassette (yellow box) flanked by loxP sites (purple arrows) to generate conditional transgenic tissue-specific Them1 overexpression (wild type) mice. **(C - D)** Adiponectin- and human thyroid hormone-binding globulin (TBG)-Cre recombination excises out the STOP cassette and brings Them1 under control of the CMV promoter in adipose tissue or hepatocytes, respectively. **(C)** Conditional transgenic adipose tissue-specific Them1 overexpression (*A*-*Them1Tg)* mice were generated by crossing wild type mice to transgenic mice expressing Cre recombinase (pink boxes) driven by the adiponectin gene promoter (light blue box). **(D)** Conditional transgenic liver-specific Them1 overexpression (*L*-*Them1Tg)* mice were generated by intravenously injecting wild type mice with adeno-associated virus 8 (AAV8) expressing Cre recombinase (Cre; pink boxes) driven by the human thyroid hormone-binding globulin (TBG) promoter (gray box). **(E)** Relative mRNA expression level of *Them1* in white adipose tissue (WAT), brown adipose tissue (BAT) and kidney extracts of *A-Them1Tg* were analyzed by quantitative real-time PCR. **(F)** Relative protein expression levels of Them1 and FLAG in brain, heart, lung, WAT, skeletal muscle (muscle), kidney and BAT of *A-Them1Tg* mice were analyzed by immunoblotting with total protein utilized to control for unequal loading. wild type, n = 5; *A-Them1Tg*, n = 5. **(G)** Relative protein expression levels of Them1 and FLAG in primary brown adipocytes cultured from wild type and *A-Them1Tg* mice were analyzed by immunoblotting with Hsp90 utilized to control for unequal loading. wild type, n = 6; *A-Them1Tg*, n = 6. **(H)** Relative protein expression levels of Them1 and FLAG in primary hepatocytes cultured from *L-Them1Tg* and *Them1^-/-^* mice were analyzed by immunoblotting with Hsp90 utilized to control for unequal loading. *L-Them1Tg*, n = 6; *Them1^-/-^*, n = 6. Data represent the mean ± s.e.m. **, P < 0.01; ***, P < 0.001; wild type vs. *A-Them1Tg*.

**Supplemental figure 5.**
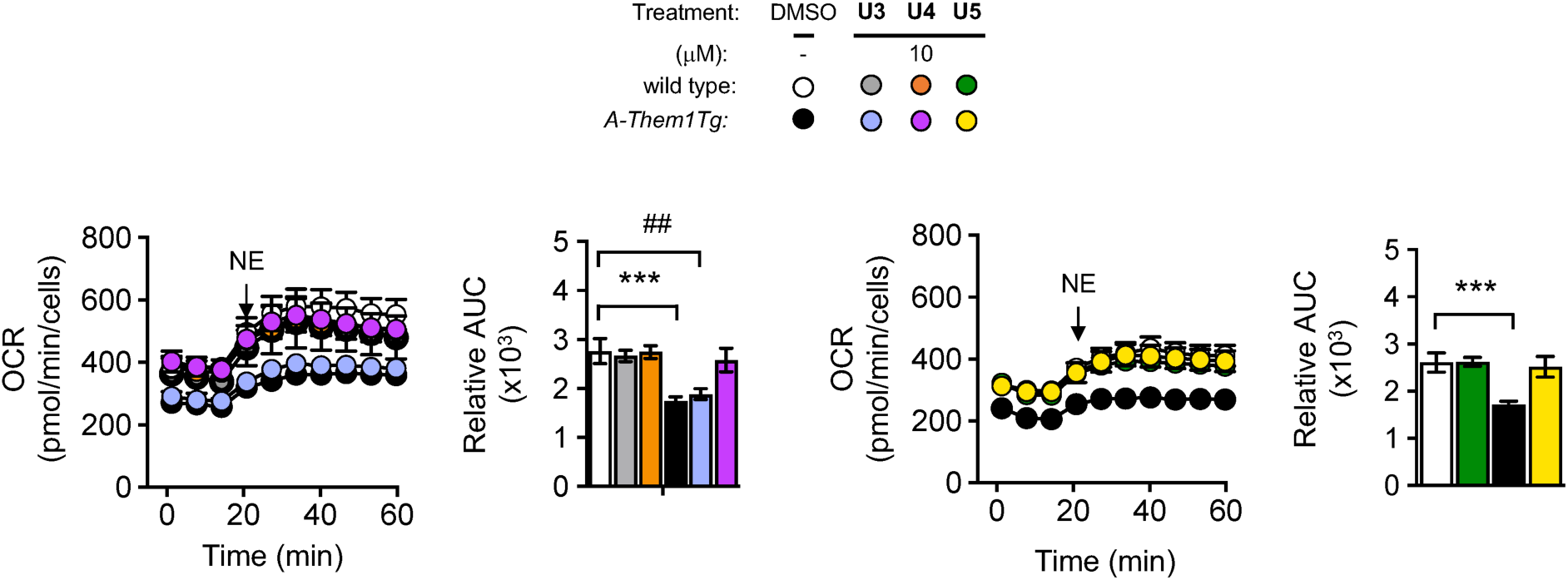
Small molecule inhibitors increase OCR in primary brown adipocytes. OCR values in primary brown adipocytes cultured from wild type and transgenic adipose tissue-specific Them1 overexpression (*A-Them1Tg*) mice following stimulation with norepinephrine (NE; 1 μM), and treatment for 30 min with the following compounds at 10 μM: **U3**, **U4** and **U5**. The bar graphs in each panel represent relative NE-stimulated values of area under the curve (AUC). Data in the graphs represent the mean ± s.e.m. of 2 to 3 independent experiments. Where not visible, standard error bars are contained within the symbol sizes. ***, P < 0.001; wild type (DMSO) vs. *A-Them1Tg* (DMSO); ^##^, P < 0.01; wild type (DMSO) vs. *A-Them1Tg* (**U3**).

**Supplemental figure 6.**
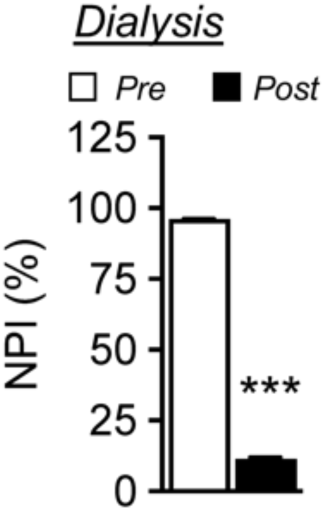
Reversibility of inhibition by synthesized compound U1. Pre-dialysis reactions were performed with recombinant His-tagged human Them1 (Them1), myristoyl (C14)-CoA and synthesized compound **U1**. Reactions subjected to an overnight dialysis were incubated with C14-CoA. Data in the graph is representative of 2 independent experiments. Data represent the mean ± s.e.m. of triplicate determinations. Where not visible, standard error bars are contained within the symbol sizes. ***, P<0.001; *Pre-* (Compound **U1**) vs. *Post-* (Compound **U1**).

**Supplemental figure 7.**
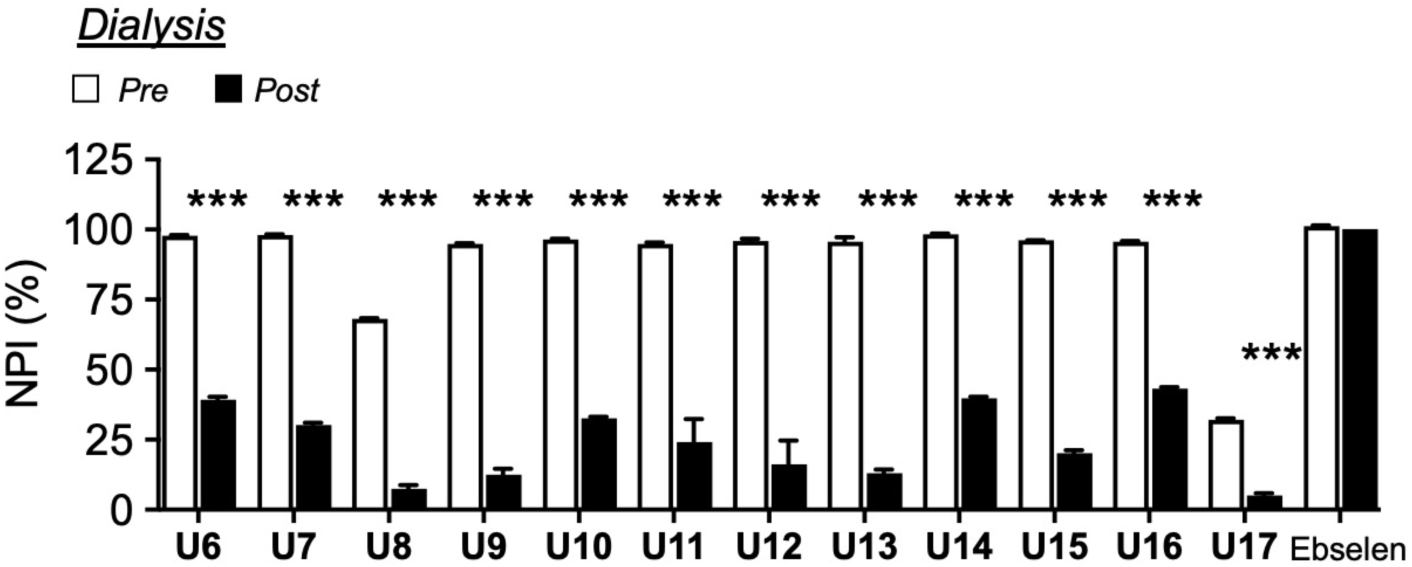
Reversibility of inhibition by compound U1 structural derivatives. Pre-dialysis reactions were performed with recombinant His-tagged human Them1 (Them1), myristoyl (C14)-CoA as well as compound **U1** structural derivatives or Ebselen [(-) control. Reactions subjected to an overnight dialysis were incubated with C14-CoA. Data in the graph is representative of 2 independent experiments. Data represent the mean ± s.e.m. of triplicate determinations. Where not visible, standard error bars are contained within the symbol sizes. ***, P<0.001; *Pre-* (Compound **U**1) vs. *Post-* (Compound **U1**).

**Supplemental figure 8.**
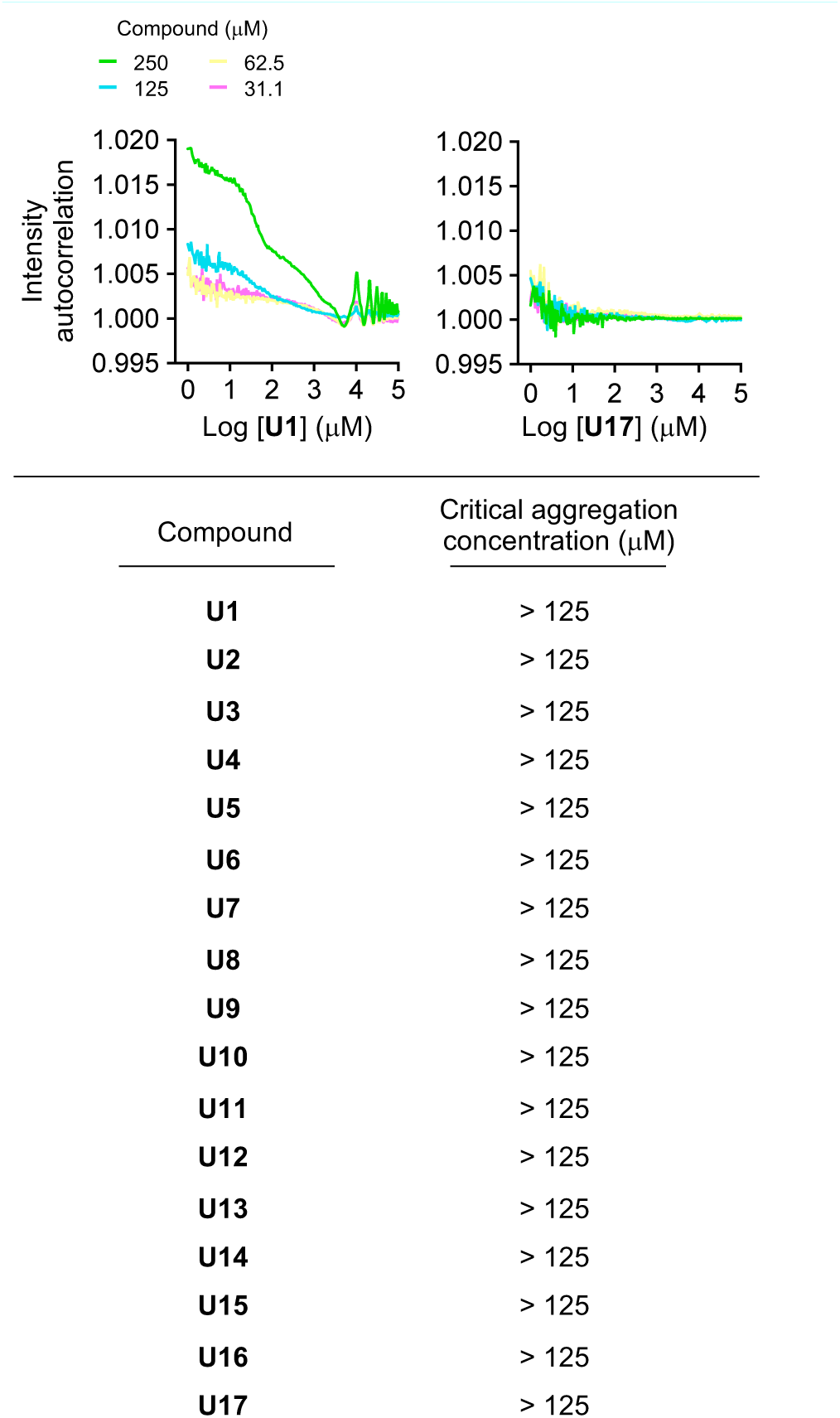
Assessment of the solubility of compound U1 and its structural derivatives. Critical aggregation concentration values of compound **U1** and its structural derivatives were determined by dynamic light scattering using serial 2-fold dilutions of compounds (250 μM – 31.5 μM) in PBS. Data represent triplicate determinations on 2 independent days.

**Supplemental figure 9.**
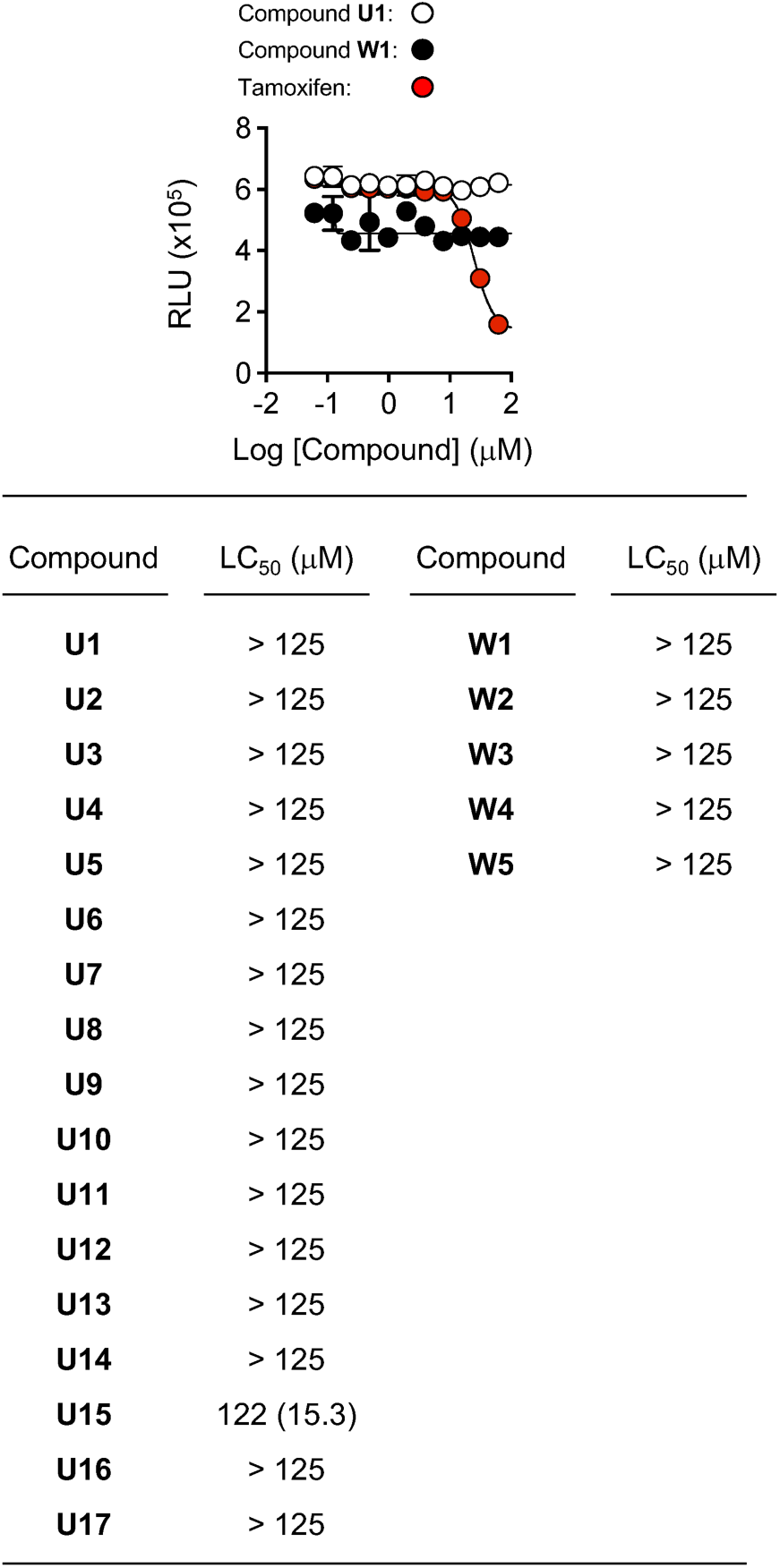
Cytotoxicity of compounds U1, W1 and their structural analogs and derivatives. Figure 16. Cytotoxicity of small molecule inhibitors with START-domain binding in primary brown adipocytes. Cell viability for compounds U1 and W1 following a 48 h treatment period in primary brown adipocytes cultured from wild type mice. Lethal concentrations (LC_50_) were > 125 μM for compounds U1 – U17 and W1 – W5. Tamoxifen served as a (+) control for the assay. Data in the graphs represent the mean of 2 independent experiments. Where not visible, standard error bars are contained within the symbol sizes.

## Supplemental methods

### Synthesis of small molecule inhibitors. Exemplary synthetic for the Compound U series

Due to facile fragmentation under electrospray conditions, no parent mass was observed by high-resolution mass spectrometry (HRMS) for all **U** series compounds. Instead, a consistent fragmentation pattern was observed that corresponds to the theoretical m/z for each compound. The observed fragmentation ions are reported for each **U** series compound below.

#### Synthesis of intermediate **I-2**

**Step 1:**

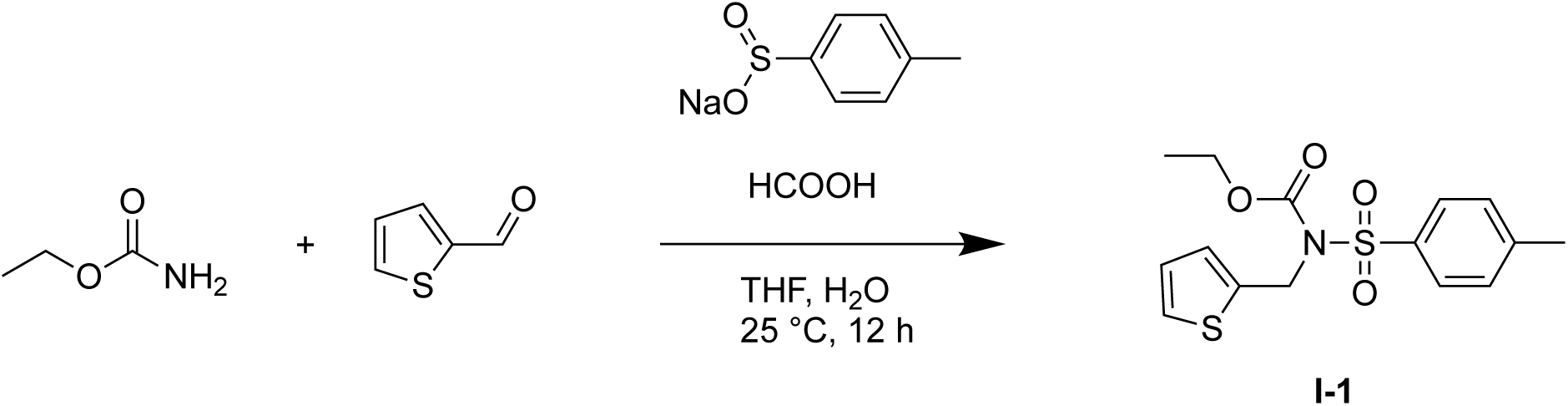

To a solution of ethyl carbamate (500 mg, 5.61 mmol, 1.00 eq) in a mixture of THF (2.00 mL) and H_2_O (5.00 mL), was added sodium 4-methylbenzenesulfinate (1.00 g, 5.61 mmol, 1.00 eq) followed by thiophene-2-carbaldehyde (**3**) (577 μL, 6.17 mmol, 1.10 eq). To this mixture was added formic acid (1.20 mL, 5.61 mmol, 1.00 eq) dropwise at 25 °C. The mixture was stirred for 12 h at 25 °C then concentrated under vacuum to obtain ethyl (thiophen-2-ylmethyl)(tosyl)carbamate (**I-1**) (1.00 g) as a yellow solid. The crude product was used without further purification.

**Step 2:**

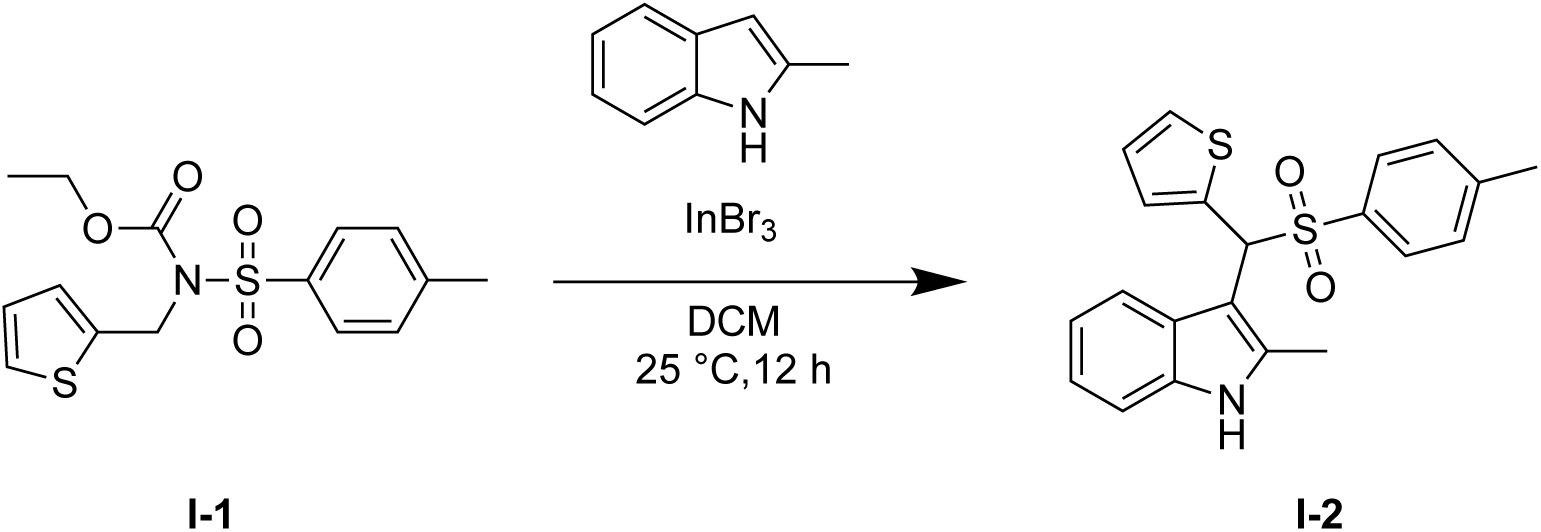

To a solution of **I-1** (1.00 g, 2.95 mmol, 1.00 *eq*) and 2-methyl indole (425 mg, 3.24 mmol, 1.10 *eq*) in DCM (10.0 mL) was added InBr_3_ (104 mg, 295 μmol, 0.10 *eq*) at 25 °C and the mixture was stirred for 12 h. The reaction mixture was diluted with water (20.0 mL) and then extracted with DCM (2X 20.0 mL each). The combined organic phase was washed with brine (2X 30.0 mL) then dried over anhydrous Na_2_SO_4_, filtered, and concentrated under vacuum. The crude product was purified by column chromatography (SiO_2_, petroleum ether: ethyl acetate = 15:1 to 1:1, petroleum ether: ethyl acetate = 2:1, R_f_ = 0.70) to obtain 2-methyl-3-(thiophen-2-yl(tosyl)methyl)-1H-indole (**I-2**) (700 mg, 1.52 mmol, 32.9% yield) as a yellow solid.

**^1^H-NMR of I-2**: (400 MHz, DMSO-*d_6_*) *δ* 11.1 (s, 1H), 7.39 (d, *J* = 8.0 Hz, 1H), 7.52 (dd, *J* = 5.2, 1.2 Hz, 1H), 7.42 (d, *J* = 8.4 Hz, 2H), 7.28 (dt, *J* = 3.6, 1.2, 0.8 Hz, 1H), 7.22 (d, *J* = 8.0 Hz, 3H), 7.03 - 6.97 (m, 2H), 6.96 - 6.91 (m, 1H), 6.22 (d, *J* = 0.8 Hz, 1H), 2.31 (s, 3H), 2.04 (s, 3H).

#### Alternate synthesis of compound **I-2.**

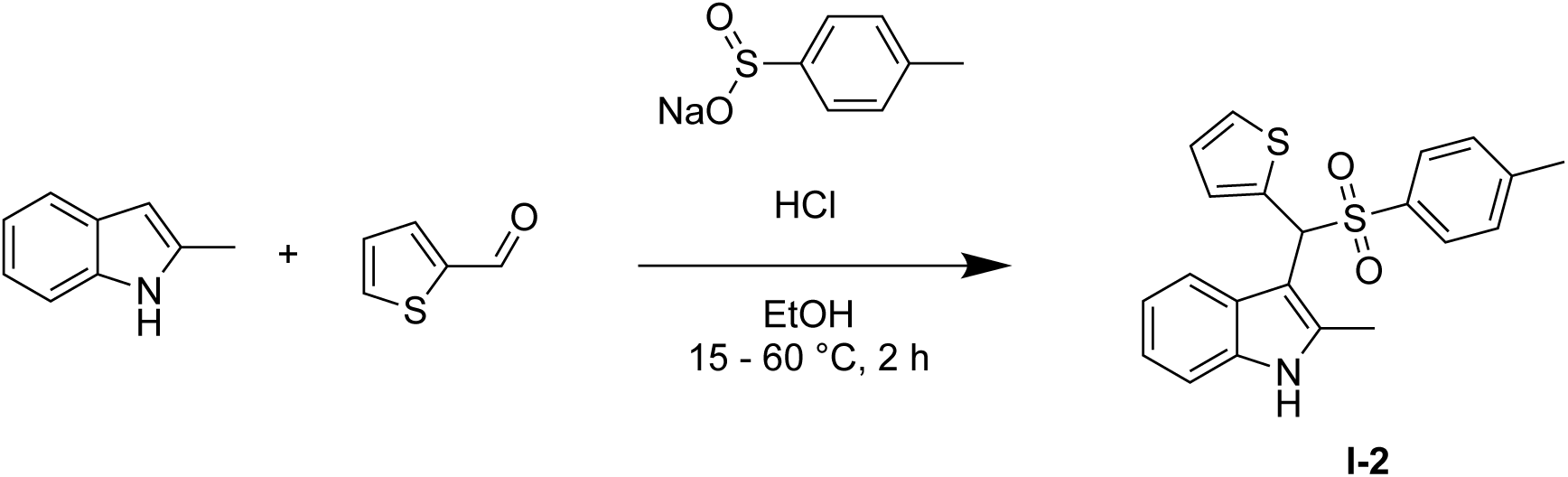

To a solution of HCl (12.0 M, 6.80 mL, 2.42 eq) in EtOH (50.0 mL) was added 2-methyl indole (5.00 g, 38.1 mmol, 1.13 eq) followed by thiophene-2-carbaldehyde (3.15 mL, 33.7 mmol, 1.00 eq) and sodium 4-methylbenzenesulfinate (7.21 g, 40.4 mmol, 1.20 eq) under N_2_ at 15 °C. The mixture was heated to 60 °C and stirred for 2 h then cooled to room temperature and extracted with ethyl acetate (3X 200 mL). The combined organic phase was washed with brine (2X 100 mL), dried over anhydrous Na_2_SO_4_, filtered, and concentrated under vacuum. The crude product was purified by column chromatography (SiO_2_, R_f_ = 0.40, petroleum ether/ethyl acetate = 1/0 to 2/1) to obtain 2-methyl-3-(thiophen-2-yl(tosyl)methyl)-1H-indole (**I-2**) (6.87 g, 13.7 mmol, 40.7 % yield) as a red solid.

**_1_H-NMR of I-2:** (400 MHz, DMSO). *δ* 11.0 (s, 1H), 7.73 (s, 1H), 7.52 (d, *J* = 0.8 Hz, 1H), 7.43 (d, *J* = 8.0 Hz, 2H), 7.28 (d, *J* = 3.2 Hz, 1H), 7.22 (d, *J* = 8.0 Hz, 3H), 7.02 - 6.93 (m, 3H), 6.22 (s, 1H), 2.32 (d, *J* = 12.4 Hz, 3H), 2.04 (s, 3H).

#### Synthesis of Compound **U1**

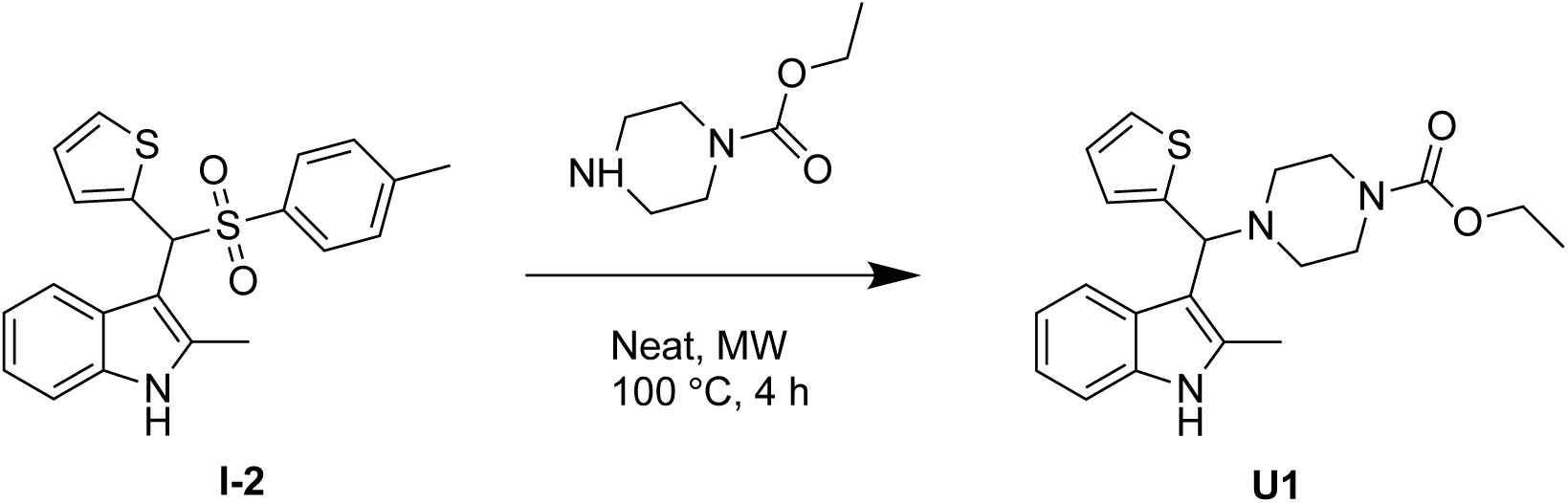

A mixture of **I-2** (150 mg, 393 μmol, 1.00 *eq*) and ethyl piperazine-1-carboxylate (926 μL, 6.32 mmol, 16.1 *eq*) was heated to 100 °C in a microwave reactor. The mixture was stirred for 4 h then cooled to room temperature and diluted with MeOH (2.00 mL). A solid was collected by filtration and purified by prep-HPLC (column, waters Xbridge C18 150 * 50mm * 10 μm; mobile phase: [water (NH_4_HCO_3_) - ACN]; B%: 27%-57%, 10 min) then further purified by pre-HPLC (column: Welch ultimate XB-SiOH 250 * 50 * 10 μm; mobile phase: [Hexane-EtOH]; B%: 1% - 30%, 15 min) to obtain ethyl 4-((2-methyl-1H-indol-3-yl)(thiophen-2-yl)methyl)piperazine-1-carboxylate (**U1**) (12.4 mg, 23.5 μmol, 5.98% yield) as a light-yellow solid.

**^1^H-NMR of U1:** (400 MHz, DMSO+D_2_O) *δ* 7.69 (d, *J* = 7.6 Hz, 1H), 7.28 - 7.20 (m, 2H), 6.97 - 6.85 (m, 4H), 4.95 (s, 1H), 4.00 - 3.94 (q, *J* = 7.2 Hz, 2H), 3.40 - 3.20 (m, 4H), 2.40 - 2.30 (m, 7H), 1.13 - 1.09 (t, *J* = 7.2 Hz, 3H).

**^13^C-NMR of U1:** (100 MHz, DMSO+D_2_O) *δ* 155.2, 148.3, 135.5, 133.5, 126.9, 126.6, 125.1, 124.9, 120.7, 120.0, 119.0, 110.0, 62.7, 61.3, 51.4, 44.1, 15.0, 12.2.

**HRMS of observed fragmentation pattern [C_14_H_12_NS]^+^ and [C_7_H_15_N_2_O_2_]^+^:** calculated m/z. 226.0685 and 159.1139; observed m/z: 226.0688 and 159.1130.

#### Synthesis of compound **U6**

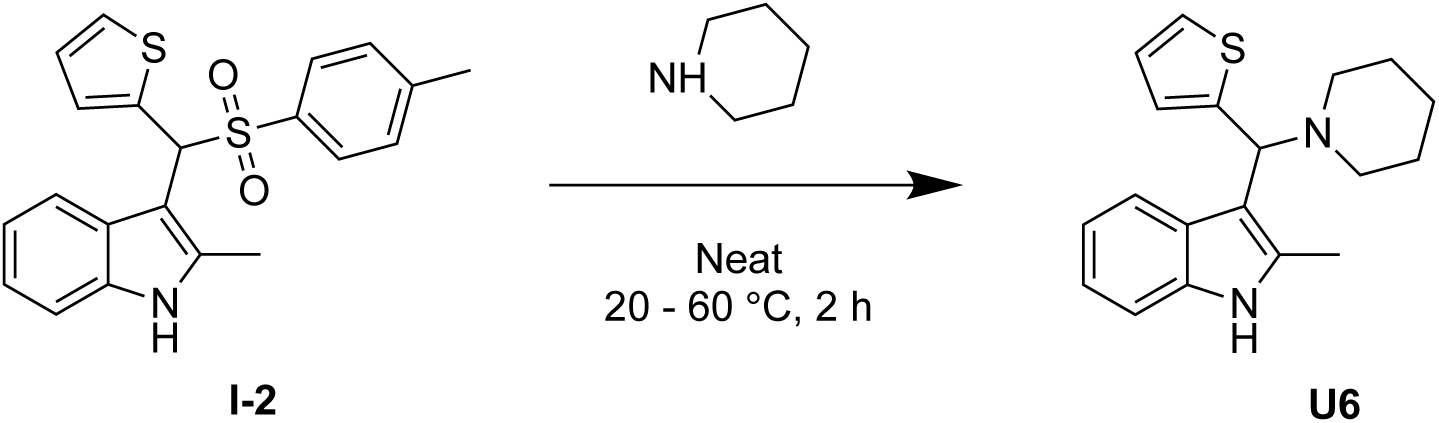

**I-2** (100 mg, 235 μmol, 1.00 eq) and piperidine (0.50 mL, 5.1 mmol, 21 eq) were mixed in a 100 mL three neck-bottomed flask at 20 °C and then heated to 60 °C for 2 h. The mixture was cooled to room temperature, diluted with MeOH (2.00 mL), and purified by pre-HPLC (column: Welch Xtimate C18 150 * 25 mm * 5 μm; mobile phase: [water (NH_3_H_2_O)-ACN]; B%: 50%-80%, 8 min) to obtain compound **U6** (53.12 mg, 171 μmol, 72.7 % yield,) as a yellow solid.

**_1_H-NMR of U6:** (400 MHz, DMSO) *δ* 10.8 (s, 1H), 7.72 (d, *J* = 8.00 Hz, 1H), 7.27 - 7.20 (m, 2H), 6.97 - 6.83 (m,4H), 4.90 (s, 1H), 2.38 (d, *J* = 25.6 Hz 7H), 1.43 (d, *J* = 58.8 Hz, 6H).

**_13_C-NMR of U6:** (100 MHz, DMSO) *δ* 149.6, 135.7, 133.3, 127.3, 126.4, 124.7, 124.2, 120.4, 120.2, 118.7, 110.8, 110.6, 63.4, 52.8, 26.4, 24.7, 12.4.

**HRMS of observed fragmentation pattern [C_14_H_12_NS]^+^ and [C_7_H_15_N_2_O_2_]^+^:** calculated m/z. 226.0685 and 159.1139; observed m/z: 226.0688 and 159.1130.

#### Synthesis of compound **U7**

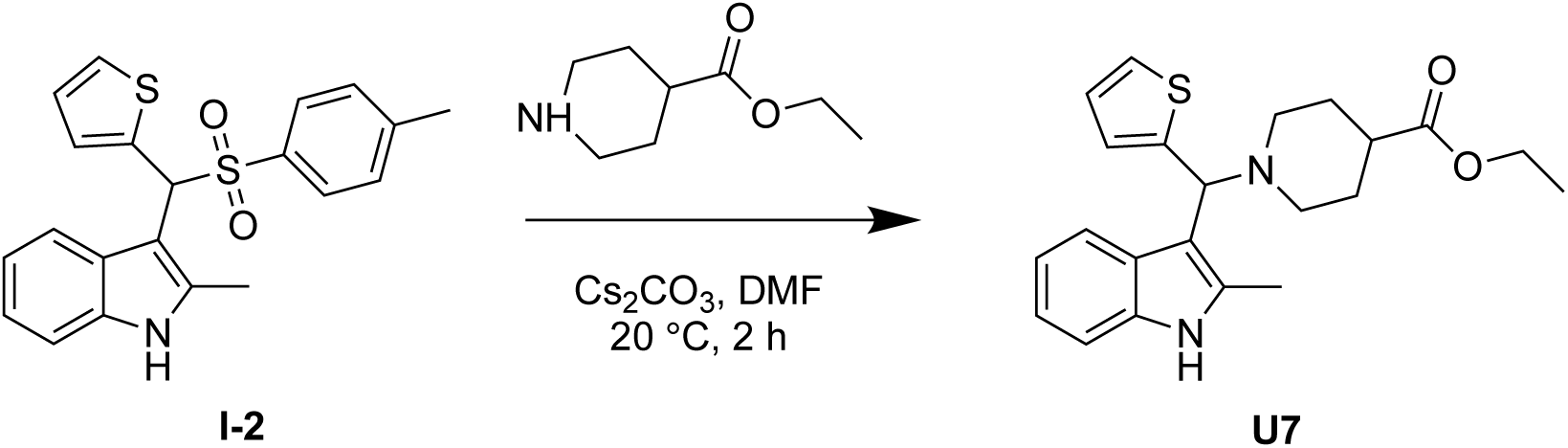

To a solution of **I-2** (100 mg, 235 μmol, 1.00 eq) in DMF (1.00 mL) at 20 °C was added iso-nipecotic acid ethyl ester (72.5 μL, 470 μmol, 2.00 eq) followed by Cs_2_CO_3_ (230 mg, 706 μmol, 3.00 eq). The mixture was stirred for 2 h at 20 °C then diluted with EtOH (2.00 mL) and purified by reverse phase HPLC (column: Welch Xtimate C18 150 * 25 mm * 5 μm; mobile phase: [water (NH_3_H_2_O)-ACN]; B%: 50%-80%, 8 min) to obtain ethyl 1-((2-methyl-1H-indol-3-yl)(thiophen-2-yl)methyl)piperidine-4-carboxylate (**U7**) (12.8 mg, 33.4 μmol, 14.2 % yield) as a brown solid.

**_1_H-NMR of U7:** (400 MHz, DMSO) *δ* 10.8 (s, 1H), 7.71 (d, *J* = 7.60 Hz, 1H), 7.27, 7.21 (dd, *J* = 5.20, 7.60 Hz, 2H), 6.97 - 6.92 (m, 3H), 6.91-6.84 (m, 1H), 4.92 (s, 1H), 4.07 - 4.01 (m, 2H), 2.91 (d, *J* = 4.40 Hz, 2H), 2.32 (d, *J* = 58.0 Hz, 3H), 2.00 (s, 1H), 1.85 (s, 1H), 1.81 (d, *J* = 13.2 Hz, 2H), 1.73 (d, *J* = 12.0 Hz, 1H), 1.63 - 1.58 (m, 2H).

**_13_C-NMR of U7:** (100 MHz, DMSO) *δ* 175.0, 149.3, 135.7, 133.2, 127.1, 126.5, 125.0, 124.4, 120.5, 120.1, 118.8, 110.8, 110.5, 62.4, 60.2, 51.8, 50.6, 28.8, 14.6, 12.4.

**HRMS of observed fragmentation pattern [C_14_H_12_NS]^+^ and [C_7_H_15_N_2_O_2_]^+^:** calculated m/z. 226.0685 and 159.1139; observed m/z: 226.0688 and 159.1130.

#### Synthesis of compound **U8**

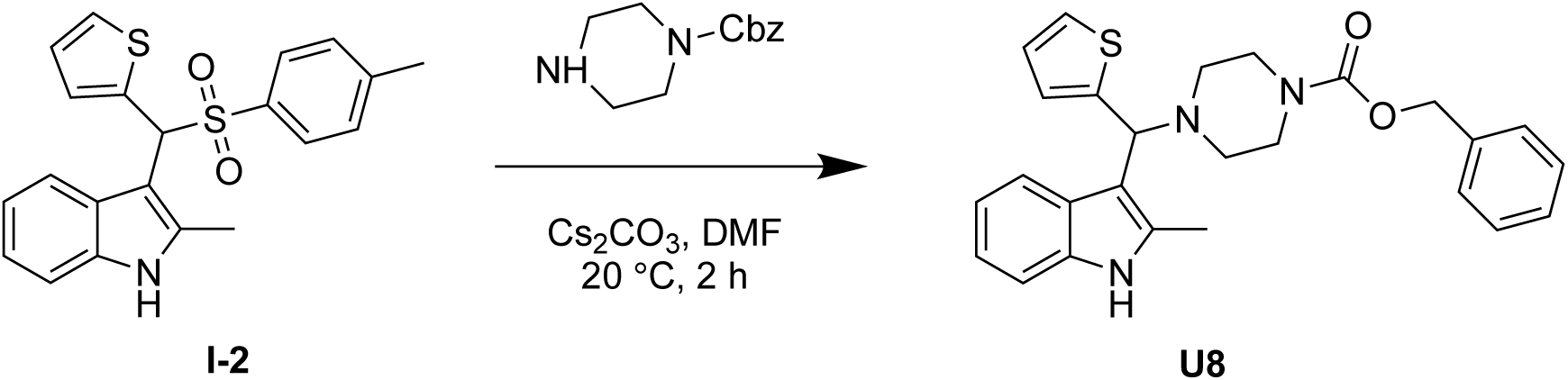

To a solution of **I-2** (100 mg, 235 μmol, 1.00 eq) in DMF (1.00 mL) at 20 °C was added cbz-piperazine (90.9 μL, 470 μmol, 2.00 eq) followed by Cs_2_CO_3_ (230 mg, 706 μmol, 3.00 eq). The mixture was stirred for 2 h at 20 °C then diluted with MeOH (2.00 mL) and purified by prep-HPLC (column: Welch Xtimate C18 150 * 25 mm * 5 μm; mobile phase: [water (NH_3_H_2_O)-ACN]; B%: 50%-80%, 8 min) to obtain benzyl 4-((2-methyl-1H-indol-3-yl)(thiophen-2-yl)methyl)piperazine-1-carboxylate (**U8**) (92.0 mg, 144 μmol, 61.4% yield) as a pink solid.

**_1_H-NMR of U8:** (400 MHz, DMSO) *δ* 10.8 (s, 1H), 7.73 (d, *J* = 7.6 Hz, 1H), 7.35 - 7.32 (m, 6H), 7.31 (d, *J* = 2.8 Hz, 1H), 7.29 - 6.97 (m, 3H), 6.92 (d, *J* = 38.8 Hz, 1H), 5.04 (s, 2H), 4.97(s, 1H), 3.41 (s, 5H), 2.42 - 2.32 (m, 7H).

**_13_C-NMR of U8:** (100 MHz, DMSO) *δ* 154.8, 148.4, 137.4, 135.7, 133.6, 128.8, 128.2, 127.9, 127.0, 126.6, 125.2, 124.8, 120.6, 120.0, 118.9, 110.9, 110.0, 66.5, 62.7, 51.5, 44.2, 12.4.

**HRMS of observed fragmentation pattern [C_14_H_12_NS]^+^ and [C_7_H_15_N_2_O_2_]^+^:** calculated m/z. 226.0685 and 159.1139; observed m/z: 226.0688 and 159.1130.

#### Synthesis of compound **U9**

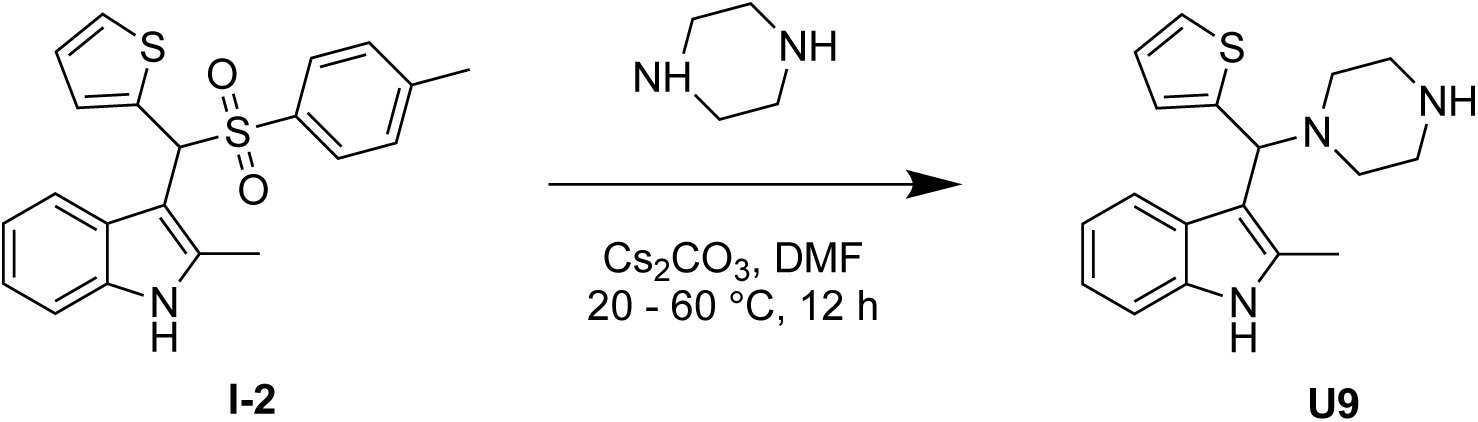

To a solution of piperazine (160 μL, 1.63 mmol, 10.0 eq) in DMF (1.00 mL) at 20 °C was added **I-2** (100 mg, 162 μmol, 1.00 eq) followed by Cs_2_CO_3_ (159 mg, 488 μmol, 3.00 eq). The mixture was heated to 60 °C and stirred for 12 h. The reaction mixture was cooled to room temperature, diluted with MeOH (1.00 mL), and purified by pre-HPLC (column: Welch Xtimate C18 150 * 25 mm * 5 μm; mobile phase: [water (NH_3_H_2_O)-ACN]; B%: 50%-80%, 8 min) to obtain 2-methyl-3-(piperazin-1-yl(thiophen-2-yl)methyl)-1H-indole (**U9**) (30 mg, 54.3 μmol, 33.3 % yield) as a yellow solid.

**_1_H-NMR of U9:** (400 MHz, DMSO) *δ* 10.9 - 10.8 (m, 1H), 7.66 (d, *J* = 8.0 Hz, 1H), 7.23 - 7.19 (m, 2H), 6.95 - 6.88 (m, 3H), 6.81 - 6.79 (m, 1H), 4.94 (d, *J* = 3.2 Hz, 1H), 2.64 - 2.57 (m, 4H), 2.40 (d, *J* = 3.2 Hz, 7H).

**_13_C-NMR of U9:** (400 MHz, DMSO) *δ* 148.9, 135.6, 133.4, 127.1, 126.4, 124.9, 124.5, 120.5, 120.1, 118.8, 117.1, 110.8, 110.2, 107.4, 62.8, 12.3.

**HRMS of observed fragmentation pattern [C_14_H_12_NS]^+^ and [C_7_H_15_N_2_O_2_]^+^:** calculated m/z. 226.0685 and 159.1139; observed m/z: 226.0688 and 159.1130.

#### Synthesis of compound **U10**

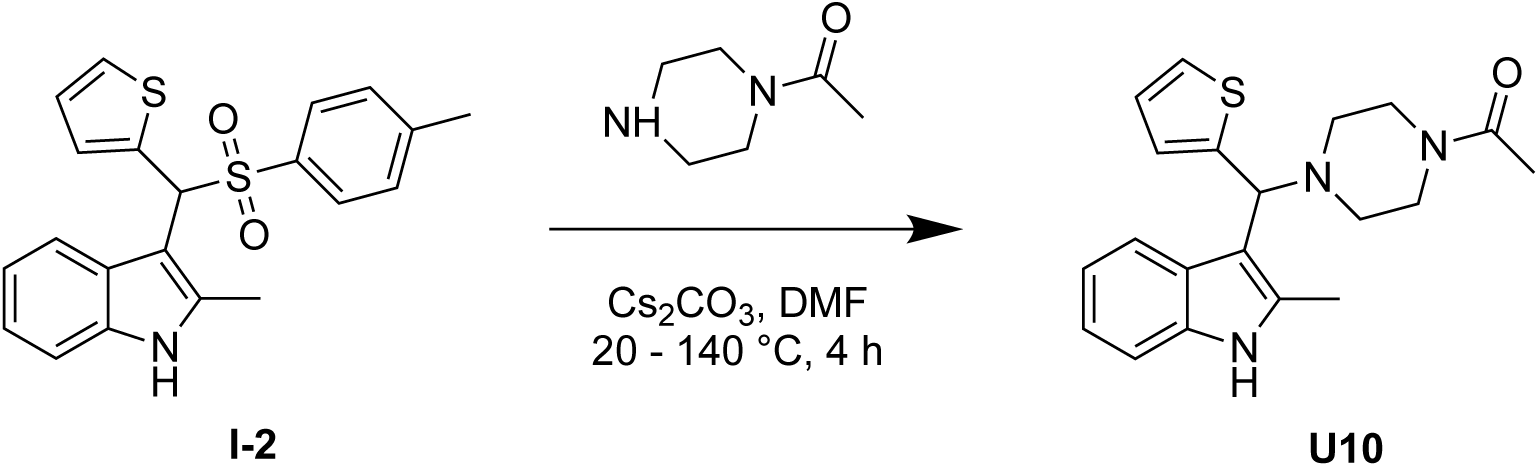

To a solution of **I-2** (100 mg, 162 μmol, 1.00 eq) in DMF (1.00 mL) at 20 °C was added 1- (piperazin-1-yl)ethan-1-one (41.7 mg, 325 μmol, 2.00 eq) followed by Cs_2_CO_3_ (159 mg, 488 μmol, 3.00 eq). The mixture heated to 140 °C and stirred for 4 h then cooled to room temperature and diluted with MeOH (1.00 mL). The mixture was purified by pre-HPLC (column: Welch Xtimate C18 150 * 25 mm * 5 μm; mobile phase: [water (NH_3_H_2_O)-ACN]; B%: 50%-80%, 8 min) to obtain 1-(4-((2-methyl-1H-indol-3-yl)(thiophen-2-yl)methyl)piperazin-1-yl)ethan-1-one (**U10**) (12.7 mg, 28.3 μmol, 17.4% yield) as a pink solid.

**_1_H-NMR of U10:** (400 MHz, DMSO) *δ* 10.9 (s, 1H), 7.73 (d, *J* = 7.6 Hz, 1H), 7.31 (d, *J* = 1.2 Hz, 1H), 7.22 (d, *J* = 7.6 Hz, 1H), 6.97 (d, *J* = 3.2 Hz, 2H), 6.87 - 6.86 (m, 2H), 4.97 (s, 1H), 3.44 - 3.41 (m, 4H), 2.45 - 2.35 (m, 7H), 1.93 (s, 3H).

**_13_C-NMR of U10:** (400 MHz, DMSO) *δ* 168.5, 148.4, 133.6, 133.2, 127.0, 126.6, 125.2, 125.0, 120.6, 120.0, 118.9, 110.9, 110.0, 62.7, 52.0, 51.5, 21.6, 12.4.

**HRMS of observed fragmentation pattern [C_14_H_12_NS]^+^ and [C_7_H_15_N_2_O_2_]^+^:** calculated m/z. 226.0685 and 159.1139; observed m/z: 226.0688 and 159.1130.

#### Synthesis of compound **U11**

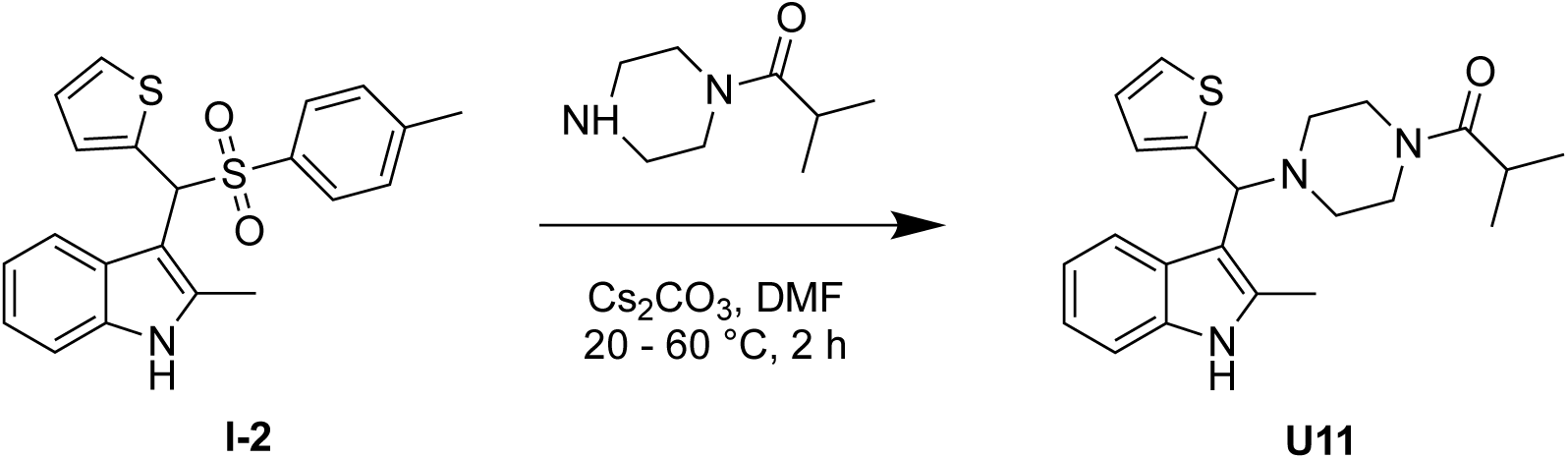

To a solution of 2-methyl-1-(piperazin-1-yl)propan-1-one (101 mg, 651 μmol, 2.00 eq) in DMF (1.00 mL) at 15 °C was added Cs_2_CO_3_ (318 mg, 976 μmol, 3.00 eq) followed by dropwise addition of **I-2** (200 mg, 325 μmol, 1.00 eq) as a solution in in DMF (2.00 mL). The mixture was heated to 60 °C and stirred for 2 h. The reaction mixture was cooled to 15 °C, filtered, and the filtrate purified by pre-HPLC (column: Welch Xtimate C18 150 * 25 mm * 5 μm; mobile phase: [water (NH_3_H_2_O)-ACN]; B%: 50%-80%, 8 min) to obtain 2-methyl-1-(4-((2-methyl-1H-indol-3-yl)(thiophen-2-yl)methyl)piperazin-1-yl)propan-1-one (**U11**) (72.9 mg, 170 μmol, 52.3 % yield) as a yellow solid.

**_1_H-NMR of U11:** (400 MHz, DMSO) *δ* 10.7 (s, 1H), 7.76 (d, *J* = 8.0 Hz, 1H), 7.30 (d, *J* = 5.2 Hz, 1H), 7.22 (d, *J* = 8.0 Hz, 1H), 6.98 - 6.93 (m, 3H), 6.92 - 6.86 (m, 1H), 4.95 (s, 1H), 3.48 (d, *J* = 8.0 Hz, 4H), 2.82 - 2.75 (m, 1H), 2.38 (d, *J* = 21.6 Hz, 7H), 0.944 - 0.928 (m, 6H).

**_13_C-NMR of U11:** (100 MHz, DMSO) *δ* 174.5, 148.4, 135.7, 133.5, 126.9, 126.5, 125.2, 124.8, 120.6, 120.0, 118.9, 110.9, 110.2, 62.8, 52.2, 51.7, 29.3, 19.8, 12.3.

**HRMS of observed fragmentation pattern [C_14_H_12_NS]^+^ and [C_7_H_15_N_2_O_2_]^+^:** calculated m/z. 226.0685 and 159.1139; observed m/z: 226.0688 and 159.1130.

#### Synthesis of compound **U12**

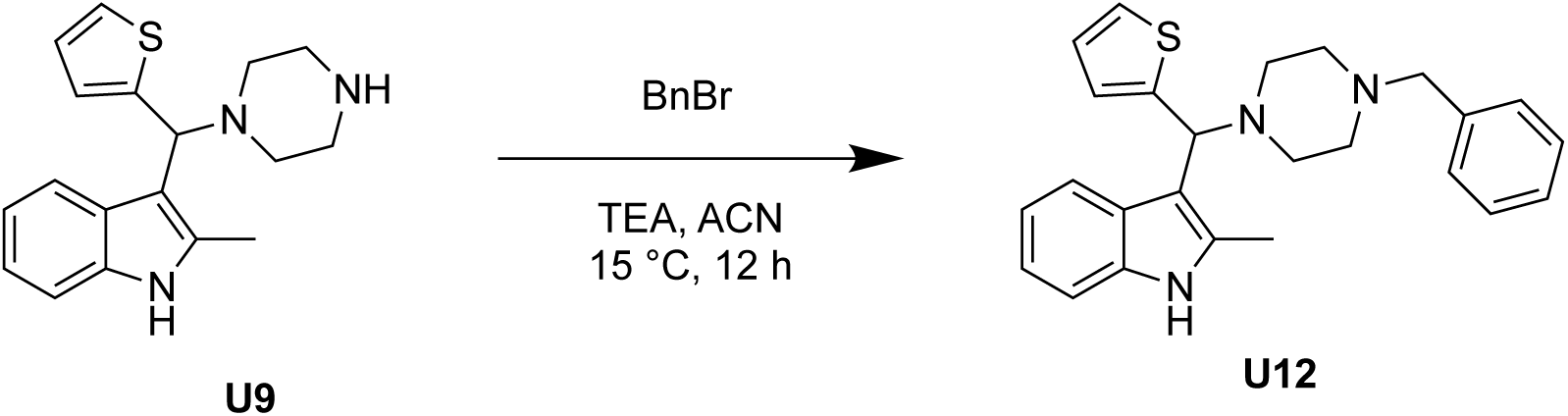

To a solution of **U9** (60 mg, 192 μmol, 1.00 eq) in ACN (1.00 mL) at 15 °C was added bromobenzene (45.7 μL, 385 μmol, 2.00 eq) followed by TEA (80.4 μL, 577 μmol, 3.00 eq). The mixture was stirred for 12 h at 15 °C then filtered and purified by pre-HPLC (column: Welch Xtimate C18 150 * 25 mm * 5 μm; mobile phase: [water (NH_3_H_2_O)-ACN]; B%: 50%-80%, 8 min) to obtain 3-((4-benzylpiperazin-1-yl)(thiophen-2-yl)methyl)-2-methyl-1H-indole (**U12**) (3.98 mg, 9.59 μmol, 4.98 % yield,) as a yellow gum.

**_1_H-NMR of U12:** (400 MHz, DMSO) *δ* 10.8 (s, 1H), 7.71 (d, *J* = 7.60 Hz, 1H), 7.28 - 7.19 (m, 7H), 6.94 - 6.84 (m, 4H), 4.92 (s, 1H), 3.44 (s, 2H), 2.52 - 2.32 (m, 11H).

**_13_C-NMR of U12:** (100 MHz, DMSO) *δ* 148.9, 138.7, 135.7, 133.3, 129.2, 128.6, 127.3, 127.1, 126.4, 124.9, 124.5, 120.5, 120.1, 118.7, 110.8, 110.4, 62.9, 62.5, 53.5, 51.7, 12.4.

**HRMS of observed fragmentation pattern [C_14_H_12_NS]^+^ and [C_7_H_15_N_2_O_2_]^+^:** calculated m/z. 226.0685 and 159.1139; observed m/z: 226.0688 and 159.1130.

#### Synthesis of compound **U13**

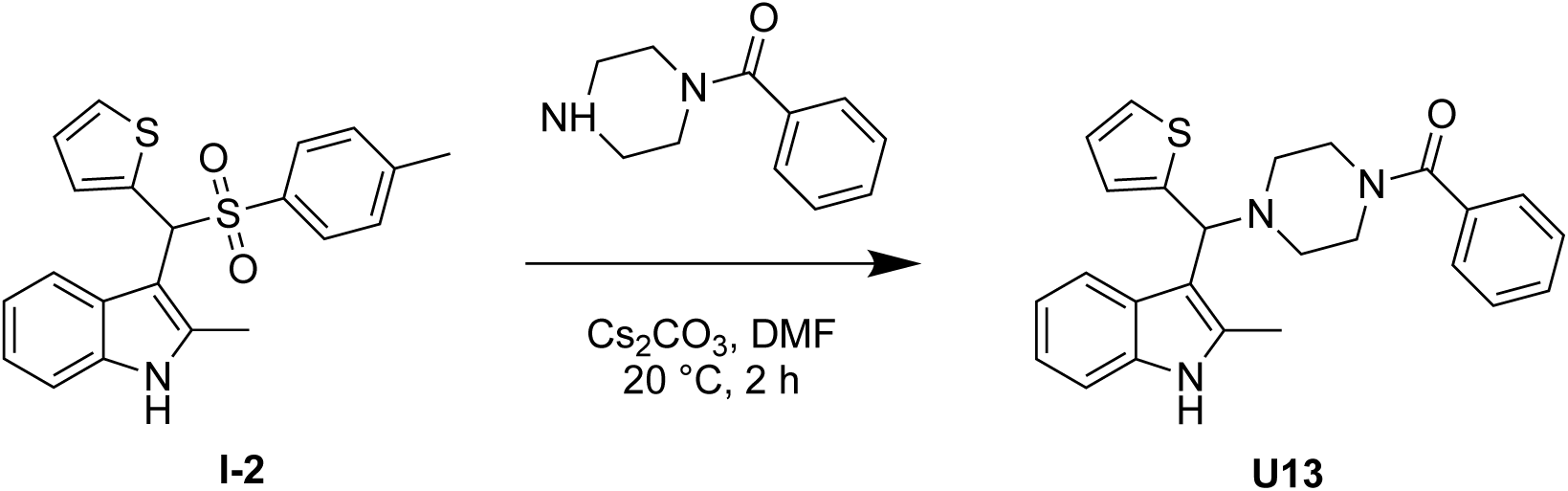

To a solution of **I-2** (100 mg, 162 μmol, 1.00 eq) in DMF (1.00 mL) at 20 °C was added phenyl(piperazin-1-yl)methanone (61.9 mg, 325 μmol, 2.00 eq) followed by Cs_2_CO_3_ (159 mg, 488 μmol, 3.00 eq). The mixture was stirred for 2 h at 20 °C then diluted with MeOH (1.00 mL) and purified by pre-HPLC (column: Welch Xtimate C18 150 * 25 mm * 5 μm; mobile phase: [water (NH_3_H_2_O)-ACN]; B%: 50%-80%, 8 min) to obtain (4-((2-methyl-1H-indol-3-yl)(thiophen-2-yl)methyl)piperazin-1-yl)(phenyl)methanone (**U13**) (37.6 mg, 87.2 μmol, 53.5 % yield) as a yellow solid.

**_1_H-NMR of U13:** (400 MHz, DMSO) *δ* 10.9 (s, 1H), 7.73 (d, *J* = 8.0 Hz, 1H), 7.40 - 7.20 (m, 7H), 6.97 - 6.86 (m, 4H), 5.00 (s, 1H), 3.64 - 3.57 (m, 4H), 2.41 (s, 7H).

**_13_C-NMR of U13:** (400 MHz, DMSO) *δ* 169.2, 148.3, 136.3, 135.7, 133.6, 129.9, 128.8, 127.4, 127.0, 126.6, 125.2, 124.9, 120.6, 120.0, 118.9, 110.9, 62.7, 51.7, 12.4.

**HRMS of observed fragmentation pattern [C_14_H_12_NS]^+^ and [C_7_H_15_N_2_O_2_]^+^:** calculated m/z. 226.0685 and 159.1139; observed m/z: 226.0688 and 159.1130.

#### Synthesis of compound **U14**

**Step 1:**

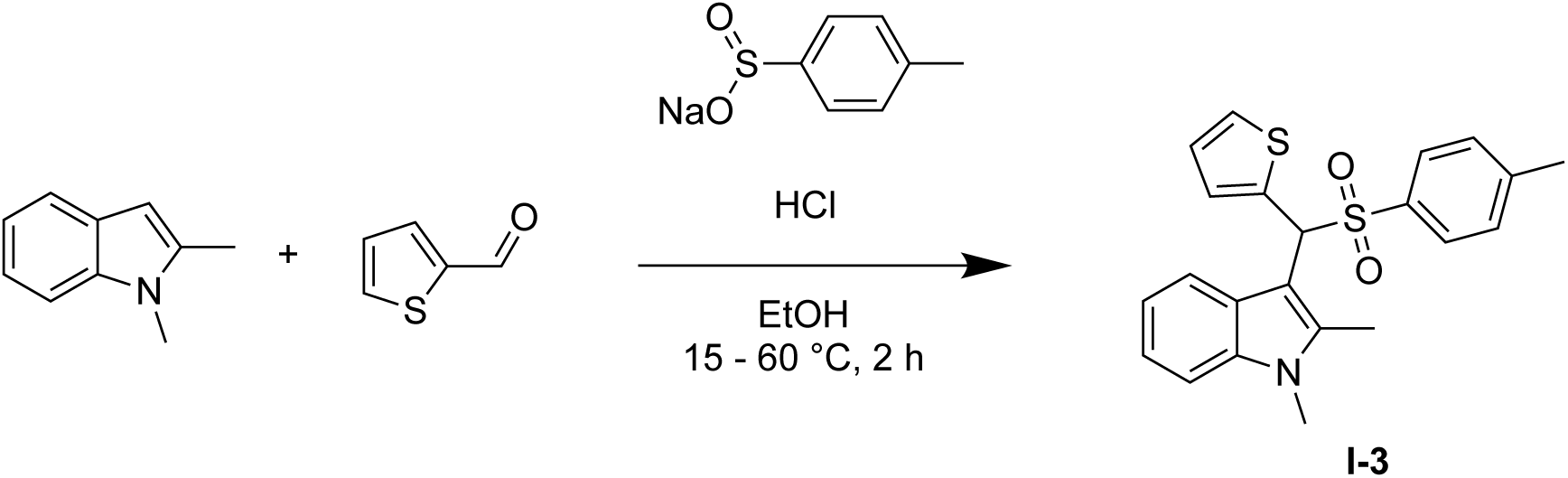

To a solution of HCl (12.0 M, 614 μL, 2.42 eq) in EtOH (25.0 mL) at 15 °C was added 1,2-dimethyl-1H-indole (500 mg, 3.44 mmol, 1.13 eq), thiophene-2-carbaldehyde (284 μL, 3.05 mmol, 1.00 eq) and sodium 4-methylbenzenesulfinate (651 mg, 3.66 mmol, 1.20 eq). The mixture was heated to 60 °C and stirred for 2 h. The reaction mixture was cooled to room temperature, diluted with water (100 mL) and extracted with ethyl acetate (3X 100 mL). The combined organic phase was washed with brine (100 mL), dried over anhydrous Na_2_SO_4_, filtered, and concentrated under vacuum to obtain 1,2-dimethyl-3-(thiophen-2-yl(tosyl)methyl)-1H-indole **(I-3)** (1.10 g, 1.52 mmol, 49.7 % yield) as a red solid.

**_1_H-NMR of I-3:** (400 MHz, DMSO) *δ* 7.74 (d, J = 8.0 Hz, 1H), 7.51 (d, *J* = 4.0 Hz, 1H), 7.44 (d, *J* = 8.0 Hz, 2H), 7.37 (d, *J* = 8.0 Hz, 1H), 7.27 - 7.20 (m, 3H), 7.10 - 7.06 (m, 1H), 7.02 - 6.91 (m, 2H), 3.56 (s, 3H), 2.34 - 2.30 (s, 3H), 2.12 (s, 3H).

**Step 2:**

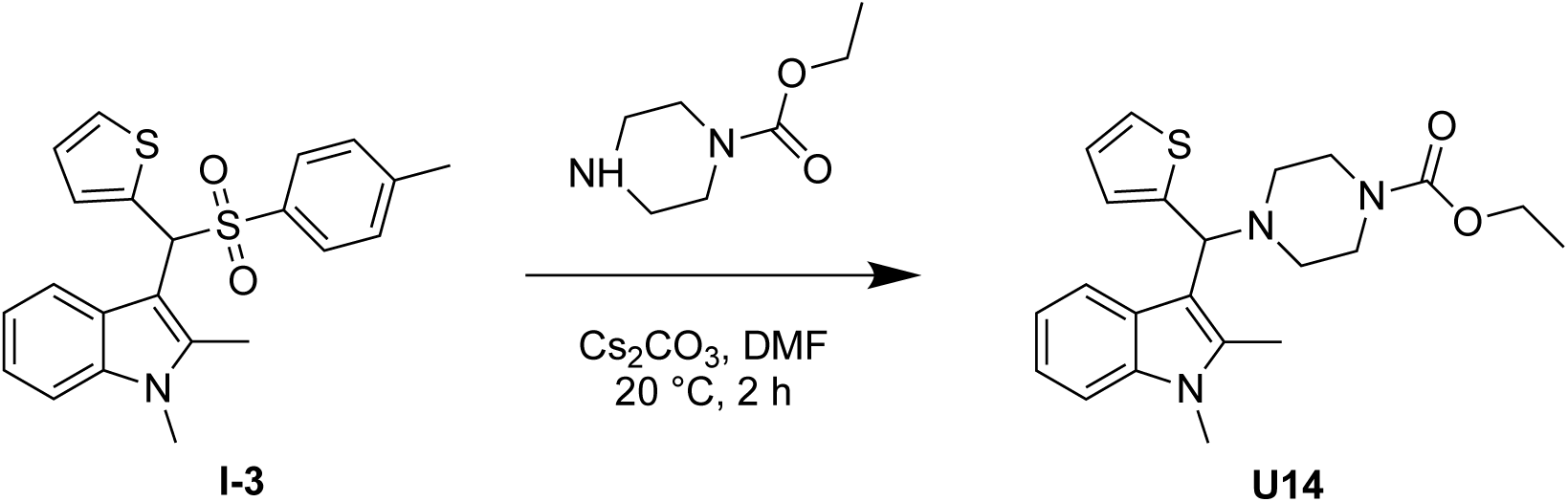

To a solution of **I-3** (1.03 g, 1.42 mmol, 1.00 eq) in DMF (10.00 mL) at 25 °C was added ethyl piperazine-1-carboxylate (414 μL, 2.83 mmol, 2.00 eq) followed by Cs_2_CO_3_ (1.38 g, 4.25 mmol, 3.00 eq). The mixture was stirred for 2 h at 25 °C then filtered and purified by pre-HPLC (column: Phenomenex C18 250 * 50 mm * 10 μm; mobile phase: [water (ammonia hydroxide v/v) - ACN]; B %: 52 % - 82 %, 10 min) to obtain ethyl 4-((1,2-dimethyl-1H-indol-3-yl)(thiophen-2-yl)methyl)piperazine-1-carboxylate (**U14**) (59.7 mg, 150 μmol, 10.6 % yield) as an off-white solid.

**_1_H-NMR of U14:** (400 MHz, DMSO) *δ* 7.79 (d, *J* = 7.6 Hz, 1H), 7.34 (d, *J* = 8.4 Hz, 1H), 7.30 (dd, *J* = 6.4, 1.2 Hz, 1H), 7.05 - 7.03 (m, 1H), 6.94 - 7.00 (m, 2H), 6.85 (m, 1H), 5.03 (s, 1H), 3.98 (q, *J* = 0.4 Hz, 2H), 3.62 (s, 3H), 3.36 (d, *J* = 5.6 Hz, 6H), 2.44 (s, 3H), 2.36 - 2.30 (m, 2H), 1.14 - 1.11 (m, 3H).

**_13_C-NMR of U14:** (100 MHz, DMSO) *δ* 155.0, 148.4, 135.7, 133.8, 135.0, 126.5, 126.0, 125.2, 124.9, 120.7, 120.2, 119.0, 110.1, 109.5, 63.0, 61.1, 51.5, 44.0, 29.8, 15.0, 10.9.

**HRMS of observed fragmentation pattern [C_14_H_12_NS]^+^ and [C_7_H_15_N_2_O_2_]^+^:** calculated m/z. 226.0685 and 159.1139; observed m/z: 226.0688 and 159.1130.

#### Synthesis of compound **U15**

**Step 1:**

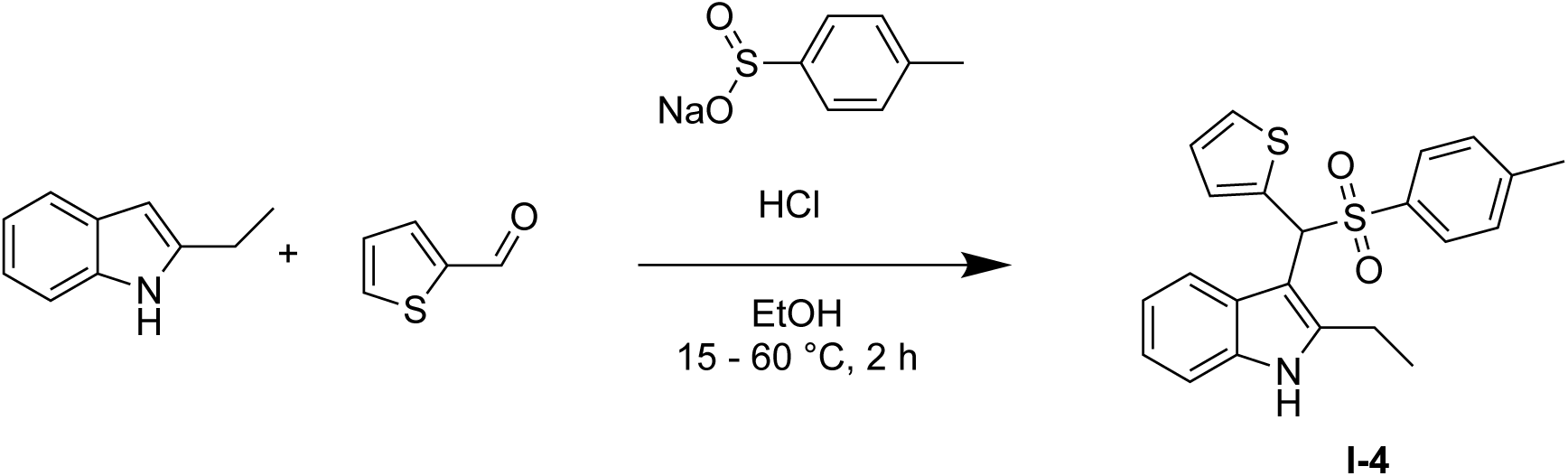

To a solution of HCl (12.0 M, 122 μL, 2.42 eq) in EtOH (5.00 mL) at 15 °C was added 2-ethyl-1H-indole (100 mg, 688 μmol, 1.13 eq) followed by thiophene-2-carbaldehyde (56.9 μL, 609 μmol, 1.00 eq) and sodium 4-methylbenzenesulfinate (130 mg, 731 μmol, 1.20 eq). The mixture was heated to 60 °C and stirred for 2 h. Upon cooling to 25 °C, water (20.0 mL) was added and then the mixture was extracted with ethyl acetate (2X 20.0 mL). The combined organic phase was washed with brine (20.0 mL), dried over anhydrous Na_2_SO_4_, filtered, and concentrated under vacuum to obtain 2-ethyl-3-(thiophen-2-yl(tosyl)methyl)-1H-indole (**I-4**) (89.0 g, 2.25 μmol, 36.9 % yield) as a red solid.

**_1_H-NMR of I-4:** (400 MHz, d4-MeOD) *δ* 7.78 (d, *J* = 8.0 Hz, 1H), 7.39 - 7.35 (m, 4H), 7.25 (d, *J* = 8.0 Hz, 1H), 7.14 (d, *J* = 8.0 Hz, 2H), 7.07 - 7.03 (m, 1H), 7.02 - 6.94 (m, 2H), 6.02 (s, 1H), 2.57 - 2.47 (m, 1H), 2.41 - 2.37(m, 1H), 2.33 (s, 3H), 0.98 - 0.95 (m, 3H).

**Step 2:**

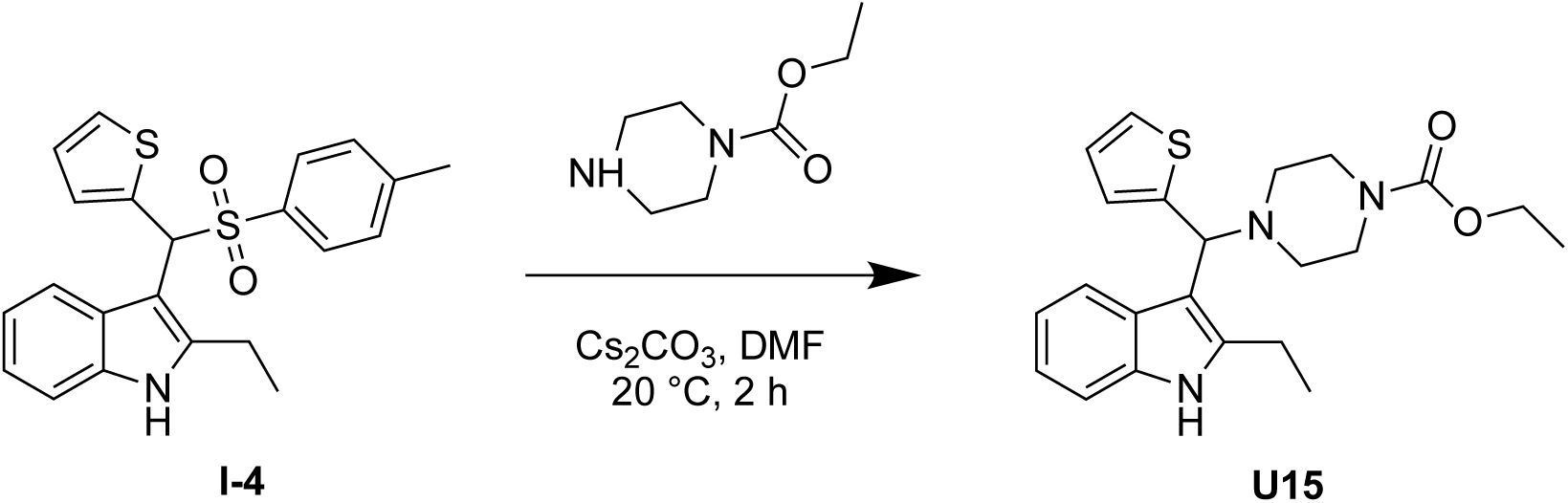

To a solution of **I-4** (89.0 mg, 225 μmol, 1.00 eq) in DMF (2.00 mL) at 15 °C was added ethyl piperazine-1-carboxylate (65.9 μL, 450 μmol, 2.00 eq) followed by Cs_2_CO_3_ (219 mg, 675 μmol, 3.00 eq). The mixture was stirred for 2 h at 15 °C then filtered and the filtrate purified by pre-HPLC (column: Welch Xtimate C18 150 * 25 mm * 5 μm; mobile phase: [water (NH_3_H_2_O) - ACN]; B %: 50 % - 80 %, 8 min) to obtain ethyl 4-((2-ethyl-1H-indol-3-yl)(thiophen-2-yl)methyl)piperazine-1-carboxylate (**U15**) (19.2 mg, 68.2 μmol, 30.3 % yield) as a white solid.

**_1_H-NMR of U15:** (400 MHz, DMSO) *δ* 10.8 (s, 1H), 7.80 (d, *J* = 8.0 Hz, 1H), 7.29 (d, *J* = 5.2 Hz, 1H), 7.23 (d, *J* = 8.0 Hz, 1H), 7.02 - 6.89 (m, 3H), 6.88 - 6.83 (m, 1H), 4.95 (s, 1H), 4.03 - 3.95 (m, 2H), 2.85 - 2.79 (m, 2H), 2.67 (s, 2H), 2.41 - 2.31 (m, 6H), 1.20 (t, *J* = 7.6 Hz, 3H), 1.13 (t, *J* = 7.2 Hz, 3H).

**_13_C-NMR of U15:** (100 MHz, DMSO) *δ* 155.0, 148.6, 139.2, 135.9, 126.6, 126.5, 125.1, 124.9, 120.7, 120.4, 118.8, 111.0, 109.5, 62.8, 61.1, 51.6, 44.1, 19.5, 15.0, 14.7.

**HRMS of observed fragmentation pattern [C_14_H_12_NS]^+^ and [C_7_H_15_N_2_O_2_]^+^:** calculated m/z. 226.0685 and 159.1139; observed m/z: 226.0688 and 159.1130.

#### Synthesis of compound **U16**

**Step 1:**

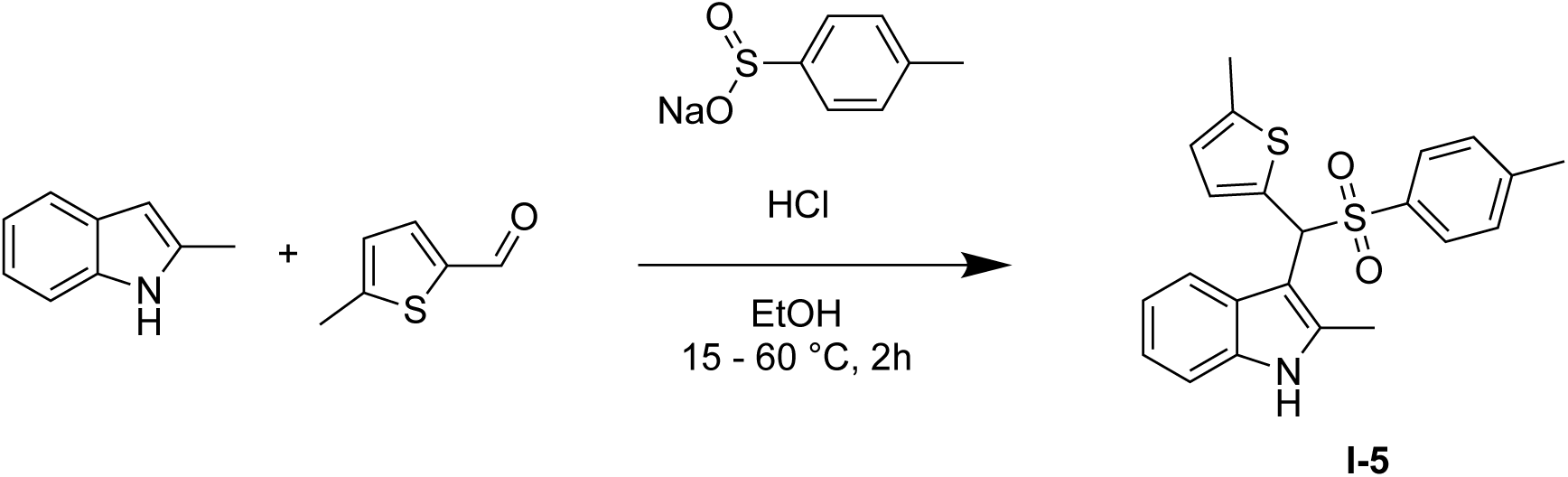

To a mixture of EtOH (5.00 mL) and HCl (12.0 M, 136 μL, 2.42 eq) at 15 °C was added sodium 4-methylbenzenesulfinate (144 mg, 809 μmol, 1.20 eq) followed by 2-methyl indole (100 mg, 762 μmol, 1.13 eq) and 5-methylthiophene-2-carbaldehyde (73.4 μL, 675 μmol, 1.00 eq). The mixture was heated to 60 °C and stirred for 2 h. Upon cooling to 15 °C, water (20.0 mL) was added, and the mixture was extracted with ethyl acetate (3X 20.0 mL). The combined organic phase was washed with brine (50.0 mL), dried over anhydrous Na_2_SO_4_, filtered, and concentrated under vacuum. The crude product was purified by pre-TLC (Petroleum ether: ethyl acetate = 2:1, R_f_ = 0.37) to obtain 2-methyl-3-((5-methylthiophen-2-yl)(tosyl)methyl)-1H-indole (**I-5**) (110 mg, 278 μmol, 41.2 % yield) as a red solid.

**^1^H-NMR of I-5:** (400 MHz, CDCl_3_) δ 7.87 (s, 1H), 7.69 (br d, *J* = 8.0 Hz, 1H), 7.41 (br d, *J* = 8.0 Hz, 2H), 7.24 (br d, *J* = 8.4 Hz, 2H), 7.16 - 7.02 (m, 7H), 6.63 (br d, *J* = 1.6 Hz, 1H), 5.75 (s, 1H), 2.46 (s, 3H), 2.33 (s, 3H).

**Step 2:**

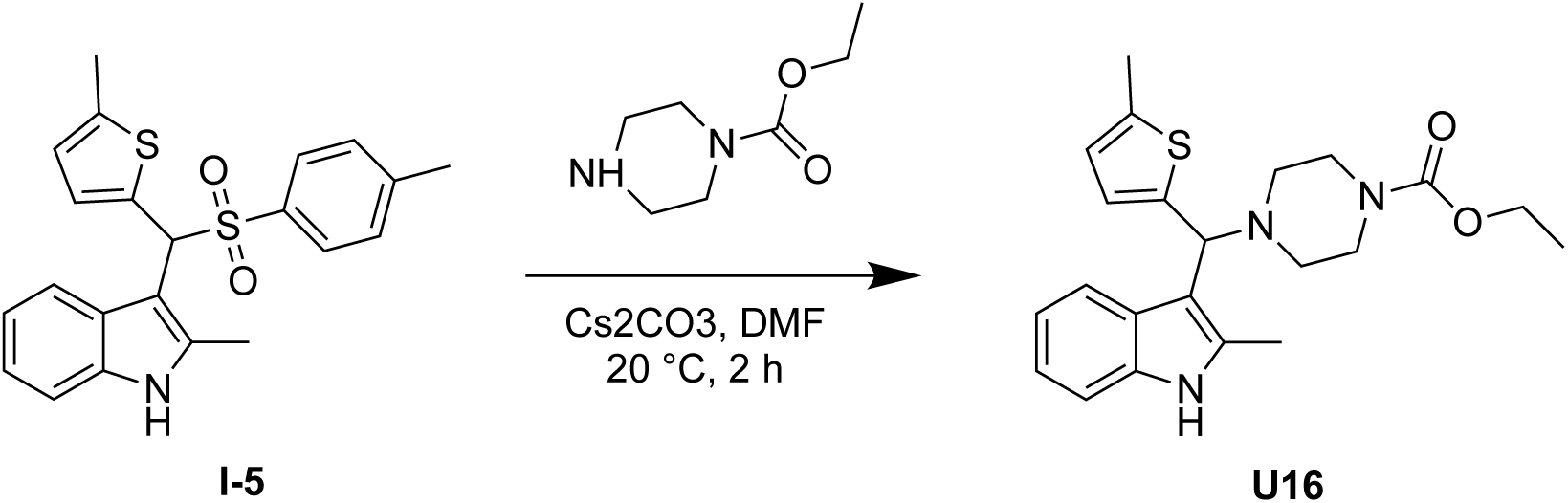

To a solution of **I-5** (110 mg, 278 μmol, 1.00 eq) in DMF (2.00 mL) at 15 °C was added ethyl piperazine-1-carboxylate (81.5 μL, 556 μmol, 2.00 eq) followed by Cs_2_CO_3_ (272 mg, 834 μmol, 3.00 eq). The mixture was stirred for 2 h at 15 °C then filtered and the filtrate purified by pre-HPLC (column: Welch Xtimate C18 150 * 25 mm * 5 μm; mobile phase: [water (NH_3_H_2_O) - ACN]; B%: 55% - 85%, 8 mins) to obtain ethyl 4-((2-methyl-1H-indol-3-yl)(5-methylthiophen-2-yl)methyl)piperazine-1-carboxylate (**U16**) (9.85 mg, 21.4 μmol, 7.72% yield) as a yellow solid.

**^1^H-NMR of U16:** (400 MHz, CDCl_3_) *δ* 7.98 - 7.91 (d, *J* = 8.4 Hz, 1H), 7.83 (s, 1H), 7.26 - 7.22 (dd, *J* = 1.2, 1.6 Hz, 1H), 7.10 (dt, *J* = 1.6, 6.8 Hz, 2H), 6.73 (d, *J* = 3.6 Hz, 1H), 6.51 - 6.46 (m, 1H), 4.80 (s, 1H), 4.11 (q, *J* = 7.2 Hz, 2H), 3.48 (br s, 4H), 2.55 - 2.47 (m, 3H), 2.45 (s, 4H), 2.38 (d, *J* = 0.8 Hz, 3H), 1.23 (t, *J* = 7.2 Hz, 3H).

**^13^C-NMR of U16:** (100 MHz, CDCl_3_) *δ* 155.5, 145.2, 139.0, 135.2, 132, 127, 124.2, 123.9, 121.2, 120.6, 119.3, 111.5, 110.1, 63.5, 61.2, 51.5,44.0, 15.4, 14.7, 12.4.

**HRMS of observed fragmentation pattern [C_14_H_12_NS]^+^ and [C_7_H_15_N_2_O_2_]^+^:** calculated m/z. 226.0685 and 159.1139; observed m/z: 226.0688 and 159.1130.

#### Synthesis of compound **U17**

**Step 1:**

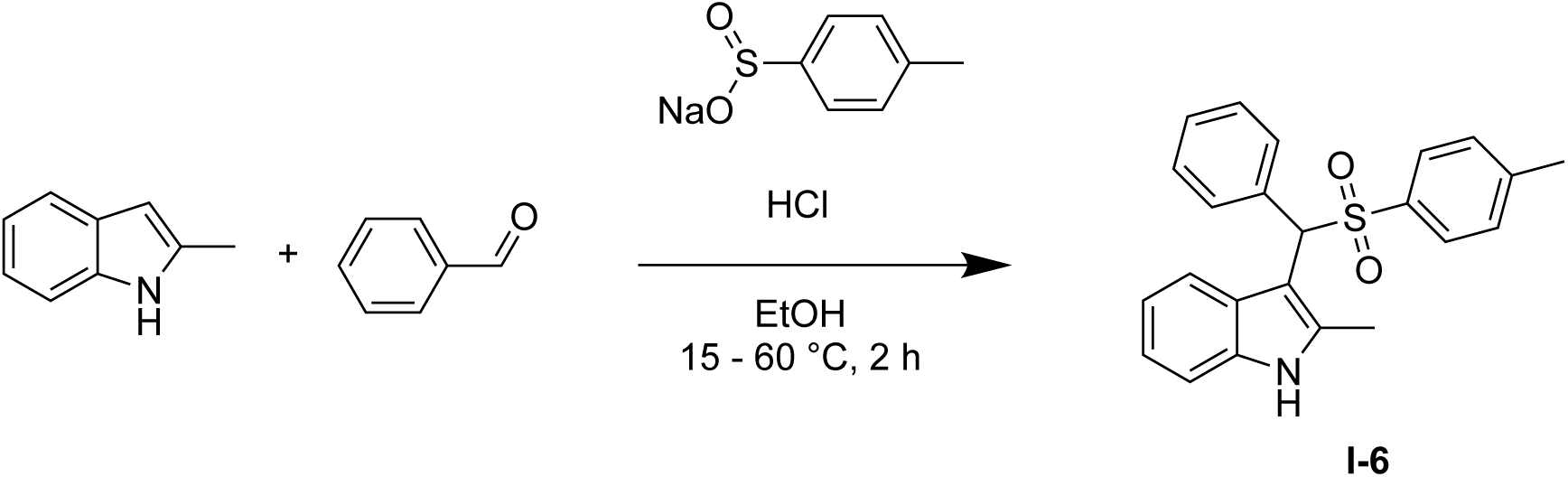

To a mixture of EtOH (5.00 mL) and HCl (12.0 M, 136 μL, 2.42 eq) at 15 °C was added sodium 4-methylbenzenesulfinate (144 mg, 809 μmol, 1.20 eq) followed by 2-methyl indole (100 mg, 762 μmol, 1.13 eq) and benzyaldehyde (68.2 μL, 675 μmol, 1.00 eq). The mixture was heated to 60 °C and stirred for 2 h. Upon cooling to 15 °C, water (20.0 mL) was added, and the mixture was extracted with ethyl acetate (3X 20.0 mL). The combined organic phase was washed with brine (50.0 mL), dried over anhydrous Na_2_SO_4_, filtered, and concentrated under vacuum. The crude product was purified by pre-TLC (Petroleum ether: ethyl acetate = 1:0 to 2:1, R_f_ = 0.35) to obtain 2-methyl-3-(phenyl(tosyl)methyl)-1H-indole (**I-6)** (135 mg, 87.7 μmol, 13.0% yield) as a red solid.

**^1^H-NMR of I-6:** (400 MHz, CDCl_3_) *δ* 7.85 (br s, 1H), 7.82 - 7.75 (m, 3H), 7.43 (d, *J* = 8.0 Hz, 2H), 7.33 (br d, *J* = 7.2 Hz, 3H), 7.23 (br d, *J* = 8.2 Hz, 2H), 7.16 - 7.09 (m, 2H), 7.07 (d, *J* = 7.6 Hz, 3H), 5.65 (s, 1H), 4.13 (q, *J* = 7.2 Hz, 1H), 2.33 (s, 3H), 2.10 (s, 3H), 2.06 (s, 1H), 1.27 (t, *J* = 7.2 Hz, 1H).

**Step 2:**

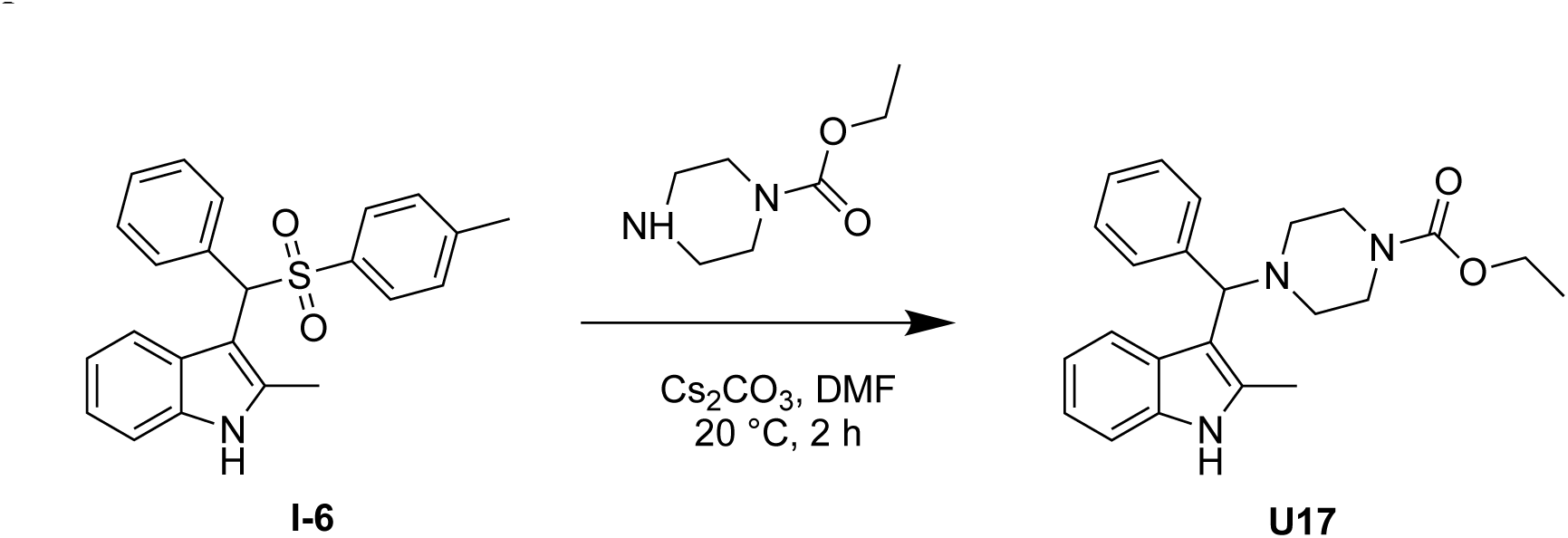

To a solution of **I-6** (135 mg, 87.7 μmol, 1.00 eq) in DMF (2.00 mL) at 15 °C was added ethyl piperazine-1-carboxylate (25.7 μL, 175 μmol, 2.00 eq) followed by Cs_2_CO_3_ (85.7 mg, 263 μmol, 3.00 eq). The mixture was stirred for 2 h at 15 °C then filtered and the filtrate purified by pre-HPLC (column: Welch Xtimate C18 150 * 25 mm * 5 μm; mobile phase: [water (NH_3_H_2_O) - ACN]; B%: 55% - 85%, 8 min) to obtain ethyl 4-((2-methyl-1H-indol-3-yl)(phenyl)methyl)piperazine-1-carboxylate (**U17**) (37.7 mg, 92.5 μmol) as a yellow solid.

**^1^H-NMR of U17:** (400 MHz, CDCl_3_) *δ* 8.05 - 7.99 (m, 1H), 7.76 (br s, 1H), 7.53 (d, *J* = 7.2 Hz, 2H), 7.26 - 7.20 (m, 3H), 7.16 (d, *J* = 7.6 Hz, 1H), 7.11 - 7.07 (m, 2H), 4.54 (s, 1H), 4.12 (q, *J* = 7.2 Hz, 2H), 3.50 (br s, 4H), 2.52 - 2.48 (m, 1H), 2.46 (s, 4H), 2.40 (br s, 2H), 1.24 (t, *J* = 7.2 Hz, 3H).

**^13^C-NMR of U17:** (400 MHz, CDCl_3_) *δ* 155.6, 143.2, 135.3, 131.7, 128.4, 127.6, 127.3, 126.6, 121.1, 120.3, 119.5, 112.5, 110.1, 68.4, 61.2, 51.9, 44.0, 14.7, 12.5.

**HRMS of observed fragmentation pattern [C_14_H_12_NS]^+^ and [C_7_H_15_N_2_O_2_]^+^:** calculated m/z. 226.0685 and 159.1139; observed m/z: 226.0688 and 159.1130.

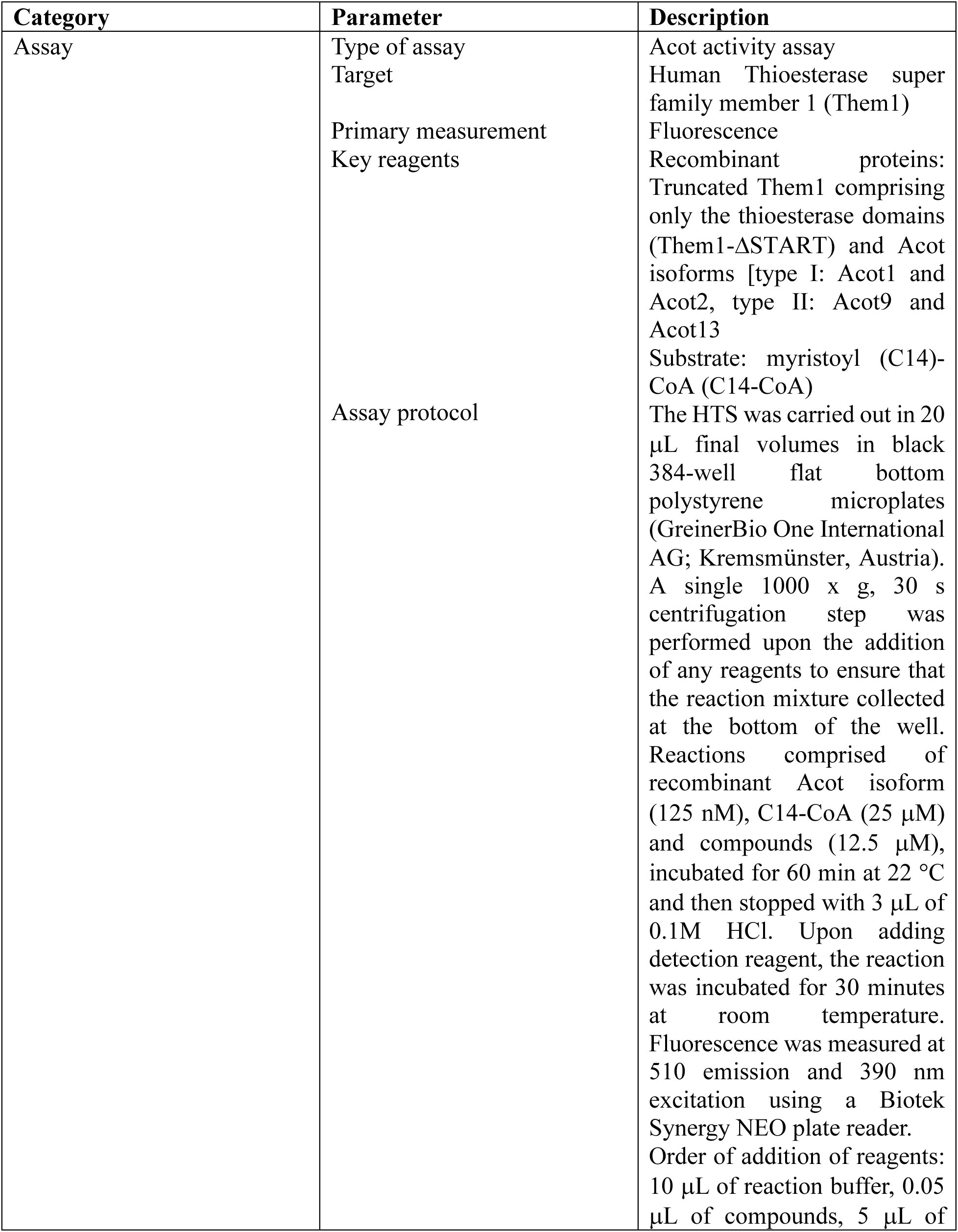
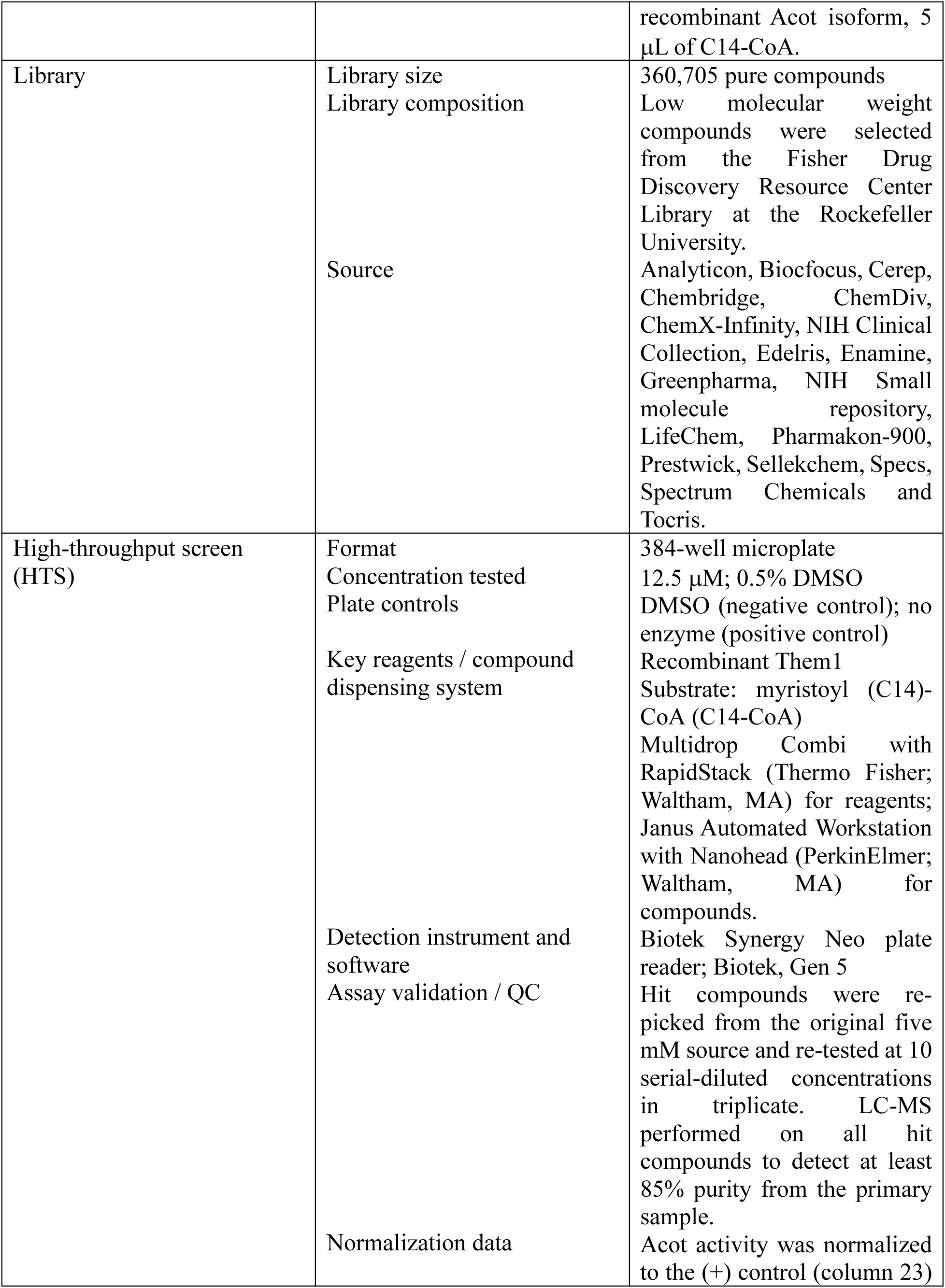
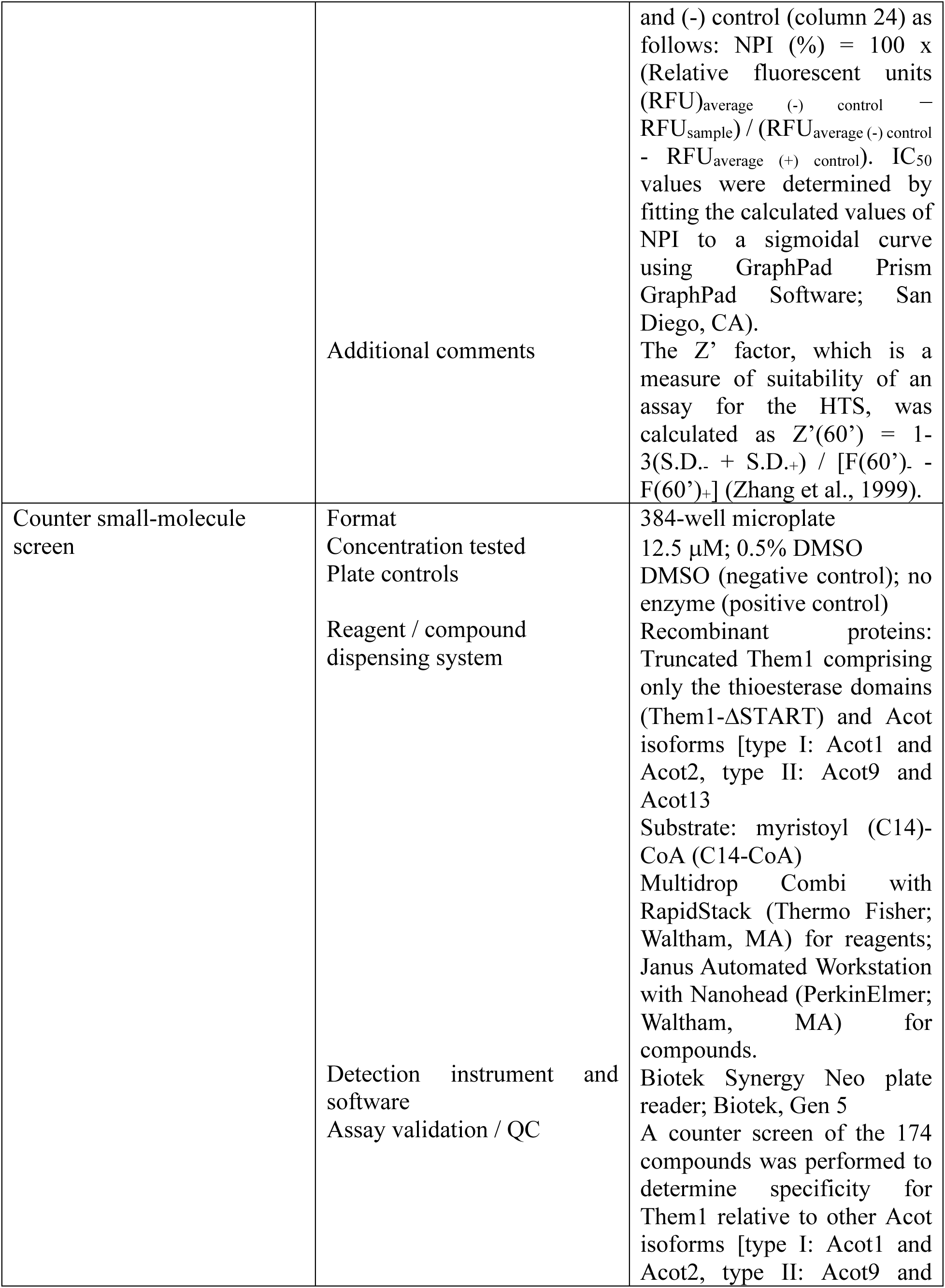
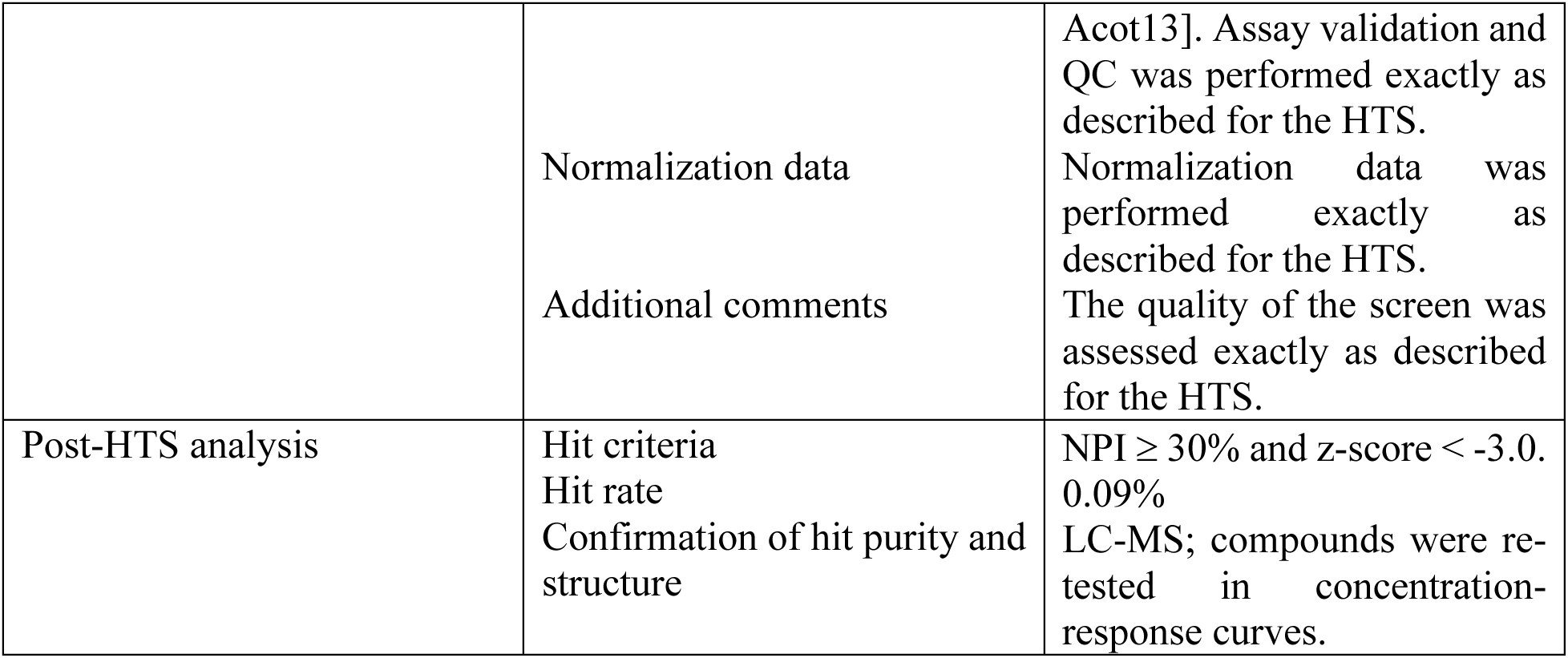
Supplemental table. Small molecule screening data.

